# Molecular diversity and phenotypic pleiotropy of genomic regulatory loci derived from human endogenous retrovirus type H (HERVH) promoter LTR7 and HERVK promoter LTR5_Hs and their impacts on pathophysiology of Modern Humans

**DOI:** 10.1101/2022.04.19.488808

**Authors:** Gennadi V. Glinsky

**Author notes:** Correspondence, Web: Guennadi V Glinskii (ucsd.edu).

## Abstract

Targeted DNA sequences conservation analyses of 17 primate species demonstrated that human endogenous retroviruses (HERV) LTR7/HERVH and LTR5_Hs/HERVK appear to have distinct evolutionary histories charted by evidence of the earliest presence and expansion of highly-conserved (HC) LTR sequences. HC-LTR7 loci were mapped to genomes of Old World Monkeys (18% of all HC-LTR7 loci), suggesting that LTR7/HERVH have entered germlines of primates in Africa after the separation of the New World Monkey lineage. HC-LTR5_Hs loci have been identified in the Gibbon’s genome (24% of all HC-LTR5_Hs loci), suggesting that LTR5_Hs/HERVK successfully colonized primates’ germlines after the segregation of Gibbons’ species. Subsequently, both LTR7 and LTR5_Hs undergo a marked ∼4-5-fold expansion in genomes of Great Apes. Timelines of quantitative expansion of both LTR7 and LTR5_Hs loci during evolution of Great Apes appear to replicate the consensus evolutionary sequence of increasing cognitive and behavioral complexities of non-human primates, which seems particularly striking for LTR7 loci and 11 distinct LTR7 subfamilies.

Consistent with previous reports, identified in this study 351 human-specific (HS) insertions of LTR7 (175 loci) and LTR5_Hs (176 loci) regulatory sequences have been linked to genes implicated in establishment and maintenance of naïve and primed pluripotent states and preimplantation embryogenesis phenotypes. Unexpectedly, HS regulatory LTRs appear linked with genes encoding markers of 12 distinct cells’ populations of fetal gonads, as well as genes implicated in physiology and pathology of human spermatogenesis, including Y-linked spermatogenic failure, oligo- and azoospermia.

Granular investigations of genes linked with eleven LTR7 subfamilies revealed that mammalian offspring survival (MOS) genes seem to remain one of consistent regulatory targets throughout ∼30 MYA of the divergent evolution of LTR7 loci. Differential GSEA of MOS versus non-MOS genes identified clearly discernable dominant enrichment patterns of phenotypic traits affected by MOS genes linked with LTR7 (562 MOS genes) and LTR5_Hs (126 MOS genes) regulatory loci across the large panel of genomics and proteomics databases reflecting a broad spectrum of human physiological and pathological traits. GSEA of LTR7-linked MOS genes identified more than 2200 significantly enriched records of human common and rare diseases and gene signatures of 466 significantly enriched records of Human Phenotype Ontology traits, including 92 genes of Autosomal Dominant Inheritance and 93 genes of Autosomal Recessive Inheritance.

One of the most informative categories of genes linked with LTR7 elements were genes implicated in functional and morphological features of central nervous system, including genes regulating synaptic transmission and protein-protein interactions at synapses, as well as gene signatures differentially regulated in cells of distinct neurodevelopmental stages and morphologically diverse cell types residing and functioning in human brain. These include Neural Stem/Precursor cells, Radial Glia cells, Bergman Glia cells, Pyramidal cells, Tanycytes, Immature neurons, Interneurons, Trigeminal neurons, GABAergic neurons, and Glutamatergic neurons. GSEA of LTR7-linked regulatory targets identified significantly enriched sets of genes encoding markers of more than 80 specialized types of neurons and markers of 521 human brain regions, most prominently, subiculum and dentate gyrus amongst top significantly enriched records. These observations were validated and extended by identification and characterization of 1944 genes comprising high-fidelity down-steam regulatory targets of LTR7 and/or LTR5_Hs loci, which are markedly enriched for genes implicated in neoplasm metastasis, intellectual disability, autism, multiple cancer types, Alzheimer’s, schizophrenia, and other brain disorders. Despite distinct evolutionary histories of retroviral LTRs, genes representing down-stream regulatory targets of LTR7 and LTR5_Hs elements exert the apparently cooperative and exceedingly broad phenotypic impacts on human physiology and pathology. Observations reported in this contribution highlight the need to accelerate the in-depth experimental and translational explorations of these important genomic determinants of Modern Humans’ health and disease states.

## Introduction

More than 3,000 DNA sequences derived from insertions of the human endogenous retrovirus type H (HERVH) are scattered across human genome at fixed non-polymorphic locations (Thomas et al., 2018). The long terminal repeats designated LTR7 harbor the promoter sequence of the HERVH and the HERVH/LTR7 family has been extensively investigated in the context of its regulatory functions and locus-specific differential expression in human preimplantation embryogenesis, human embryonic and pluripotent stem cells (Fort et al., 2014; Gemmell et al., 2015; Glinsky et al., 2018; Göke et al.,2015; Izsvák et al., 2016; Kelley and Rinn, 2012; Loewer et al., 2010; Lu et al., 2014; Ohnuki et al., 2014; Pontis et al., 2019; Römer et al., 2017; Santoni et al., 2012; Takahashi et al., 2021 Theunissen et al., 2016; Wang et al., 2014; Zhang et al., 2019),

It has been reported that HERVH/LTR7s harbor binding sites for master pluripotency transcription factors OCT4, NANOG, SP1, and SOX2 which bind HERVH/LTRs and activate their expression (Glinsky, 2015; Göke et al., 2015; Ito et al., 2017; Kelley and Rinn, 2012; Kunarso et al., 2010; Ohnuki et al., 2014; Pontis et al., 2019; Santoni et al., 2012).

Most of the previous studies have considered HERVH/LTR7 insertions in human genome as functionally homogenous genomic regulatory elements (Bao et al., 2015; Gemmell et al., 2019; Göke et al.,2015; Izsvák et al., 2016; Lu et al., 2014; Storer et al., 2021; Wang et al., 2014; Zhang et al., 2019) and regarded the entire family of >3,000 insertion sites as one monophyletic entity.

In contrast, application of a ‘phyloregulatory’ approach that integrates chromatin state features as well as regulatory and expression profiling genomics data to a phylogenetic analysis of LTR7 sequences facilitated discovery of new insights into the remarkable diversity of origin, evolution, and transcriptional activities of the HERVH/LTR7 family (Carter et al., 2022). This granular interrogation of fine molecular structures of LTR7 elements and their evolutionary history has revealed striking genetic and regulatory distinctions amongst LTR7 elements by demonstrating that LTR7 sequences represent a polyphyletic group composed of at least eight monophyletic subfamilies (Carter et al., 2022).

Collectively, these findings indicate that the HERVH/LTR7 family underwent the extensive diversification of LTR sequences during primate evolution through a combination of point mutations, indels, and recombination events facilitating the gain, loss, and exchange of multiple cis-regulatory modules. These processes are likely underlying the apparent functional partitioning of LTR7 transcription regulatory activities during primates’ preimplantation embryogenesis and maintenance of a pluripotency phenotype (Carter et al., 2022).

Human endogenous retrovirus type K [HERVK (HTML-2)] is he most recently endogenized primate-specific retrovirus and all human-specific and human-polymorphic HERVK insertions are associated with a specific LTR subtype designated LTR5_Hs (Hanke et al., 2016; Subramanian et al., 2011). It has been reported that LTR5_Hs/HERVK manifests transcriptional and biological activities in human preimplantation embryos and in naïve hESCs (Grow et al., 2015). Notably, LTR5_Hs elements acquire enhancer-like chromatin state signatures concomitantly with transcriptional reactivation of HERVK (Grow et al., 2015). Genome-wide CRISPR-guided activation and interference experiments targeting LTR5_Hs elements demonstrated global long-range effects on expression of human genes consistent with postulated functions of LTR5_Hs as distal enhancers (Fuentes et al., 2018).

In the present study, detailed analyses of evolutionary conservation patterns of eleven LTR7 subfamilies were carried out to understand when HERVH/LTR7 family has infiltrated the primates’ germline at different evolutionary time points and how they have achieved quantitatively and qualitatively various levels of genomic amplification in distinct primate species during evolution. To highlight connectivity patterns between the LTR7 structural diversity and the phenotypic pleiotropy of putative regulatory functions of LTR7 elements, the GREAT algorithm was employed to identify and characterize LTR7-linked genes. Comprehensive Gene Set Enrichment Analyses (GSEA) of LTR7-linked genes were performed to infer potential phenotypic impacts of LTR7 regulatory networks and findings were juxtaposed to results of analyses of LTR5_Hs elements.

## Methods

### Data source and analytical protocols

A total of 3354 LTR7 loci and 606 LTR5_Hs loci residing at fixed non-polymorphic locations in genomes of Modern Humans (hg38 human reference genome database) analyzed in this study were reported previously (Carter et al., 2022; Fuentes et al., 2018). Solely publicly available datasets and resources were used in this contribution. The significance of the differences in the expected and observed numbers of events was calculated using two-tailed Fisher’s exact test. Additional proximity placement enrichment and gene set enrichment tests were performed for individual classes of regulatory sequences taking into account the position and size in bp of corresponding genomic regions, size distributions in human cells of topologically associating domains, distances to putative regulatory targets, bona fide regulatory targets identified in targeted genetic interference and/or epigenetic silencing experiments, details of methodological and analytical approaches of which were reported previously [Barakat et al., 2018; Fuentes et al., 2018; Glinsky, 2015; 2016; 2018 - 2021; Guffanti et al., 2018; McLean et al., 2010; 2011; Pontis et al., 2019; Wang et al., 2014].

### Gene set enrichment and genome-wide proximity placement analyses

Gene set enrichment analyses (GSEA) were carried-out using the Enrichr bioinformatics platform, which enables the interrogation of nearly 200,000 gene sets from more than 100 gene set libraries. The Enrichr API (January 2018 through March 2022 releases) [Chen et al., 2013; Kuleshov et al., 2016; Xie et al., 2021] was used to test genes linked to regulatory LTRs of interest for significant enrichment in numerous functional categories. When technically feasible, larger sets of genes comprising several thousand entries were analyzed. Regulatory connectivity maps between LTRs and coding genes and additional functional enrichment analyses were performed with the GREAT algorithm [McLean et al., 2010; 2011] applying default settings at differing maximum extension thresholds as previously reported (Glinsky, 2020 - 2021). The reproducibility of the results was validated by implementing two releases of the GREAT algorithm: GREAT version 3.0.0 (2/15/2015 to 08/18/2019) and GREAT version 4.0.4 (08/19/2019) as well as two releases of the human genome reference database (hg19 and hg38). The GREAT algorithm allows investigators to identify and annotate the genome-wide connectivity networks of user-defined distal regulatory loci and their putative target genes. Concurrently, the GREAT algorithm performs functional annotations and analyses of statistical enrichment of annotations of identified genes, thus enabling the inference of potential biological significance of interrogated genomic regulatory networks (GRNs). Genome-wide Proximity Placement Analysis (GPPA) of distinct genomic features co-localizing with LTRs and human-specific regulatory sequences (HSRS) was carried out as described previously and initially implemented for interrogation of human-specific transcription factor binding sites and other candidate HSRS [Glnsky et al., 2018; 2019; Glinsky, 2015 – 2021; Guffanti et al., 2018].

Targeted differential GSEA were employed to infer the relative contributions of distinct subsets of genes on phenotypes of interest. In brief, to gain insights into biological effects of LTRs and infer potential mechanisms of biological activities, multiple sets of differentially-expressed genes (DEGs) and/or coding genes representing putative regulatory targets of LTRs were identified. These gene sets comprising from dozens to several thousand individual genetic loci were defined at multiple significance levels of corresponding statistical metrics and analyzed using differential GSEA applied to ∼30 genomics and proteomics databases. This approach was successfully implemented for identification and characterization of human-specific regulatory networks governed by human-specific transcription factor-binding sites [Glnsky et al., 2018; 2019; Glinsky, 2015 – 2021; Guffanti et al., 2018] and functional enhancer elements [Barakat et al., 2018; Glnsky et al., 2018; Glinsky and Barakat, 2019; Glinsky, 2015 – 2021], 13,824 genes associated with 59,732 human-specific regulatory sequences [Glinsky, 2020a], 8,405 genes associated with 35,074 human-specific neuroregulatory single-nucleotide changes [Glinsky, 2020b], different subsets of 8,384 genes regulated by stem cell-associated retroviral sequences [Glinsky, 2021], as well as human genes and medicinal molecules affecting the susceptibility to SARS-CoV-2 coronavirus [Glinsky, 2020c].

According to a standard analytical protocol, differential GSEA entail interrogations of specific sets of DEGs comprising LTR-regulated genes using distinct genomic databases, including comprehensive pathway enrichment Gene Ontology (GO) analyses. Upon completion, these analyses were followed by in-depth interrogations of the identified significantly-enriched genes employing selected genomic databases deemed most statistically informative during the initial GSEA. In all tables and plots (unless stated otherwise), in addition to the nominal p values and adjusted p values, the “combined score” calculated by Enrichr software is reported, which is a product of the significance estimate and the magnitude of enrichment (combined score c = log(p) * z, where p is the Fisher’s exact test p-value and z is the z-score deviation from the expected rank).

### Statistical Analyses of the Publicly Available Datasets

All statistical analyses of the publicly available genomic datasets, including error rate estimates, background and technical noise measurements and filtering, feature peak calling, feature selection, assignments of genomic coordinates to the corresponding builds of the reference human genome, and data visualization, were performed exactly as reported in the original publications and associated references linked to the corresponding data visualization tracks (http://genome.ucsc.edu/). Any modifications or new elements of statistical analyses are described in the corresponding sections of the Results. Statistical significance of the Pearson correlation coefficients was determined using GraphPad Prism version 6.00 software. Both nominal and Bonferroni adjusted p values were estimated and considered. The significance of the differences in the numbers of events between the groups was calculated using two-sided Fisher’s exact and Chi-square test, and the significance of the overlap between the events was determined using the hypergeometric distribution test [Tavazoie et al., 1999].

## Results

### LTR family-specific and granular analyses of evolutionary origin, expansion, conservation, and divergence of HERVH promoter LTR7 and HERVK promoter LTR5_Hs during primate evolution

Precise timelines of transitions of retroviruses from the state of exogenous infection agents to endogenous retroviral sequences integrated into host genomes remain unknown. This information could be inferred from the comparative analyses of highly conserved retrovirus-derived loci in genomes of multiple distinct primate species with known species’ divergence time from the last extinct common ancestor (ECA) during primates’ evolution. To this end, fixed non-polymorphic sequences of 3354 LTR7 loci and 606 LTR5_Hs loci residing in genomes of Modern Humans (hg38 human reference genome database) were retrieved and the number of highly conserved loci in genomes of sixteen non-human primates (NHP) were determined (**Figure 1**). Results of these analyses demonstrate that LTR7 and LTR5_Hs loci appear to have clearly distinguishable evolutionary histories. The consistent earliest presence of LTR7 loci could be mapped to genomes of Old World Monkeys (**Figure 1A**), suggesting that LTR7/HERVH retroviruses have entered germlines of the primate lineage after the separation of the New World Monkey lineage. Subsequently, endogenous LTR7/HERVH retroviruses appear to undergo a marked 4-5-fold expansion in genomes of Great Apes (**Figure 1A**). While small numbers (∼1%) of highly conserved LTR5_Hs loci could be traced to genomes of Old World Monkeys (**Figure 1B**), the consistent earliest presence of highly conserved LTR5_Hs loci has been observed in the Gibbon’s genome (143 loci; 24% of HC LTR5_Hs loci residing in human genomes). Taken together, these findings suggest that LTR5_Hs/HERVK retroviruses successfully colonized germlines of the primate lineage after the segregation of Gibbons’ species and subsequently underwent a marked expansion in genomes of Great Apes (**Figure 1B**).

**Figure 1.**
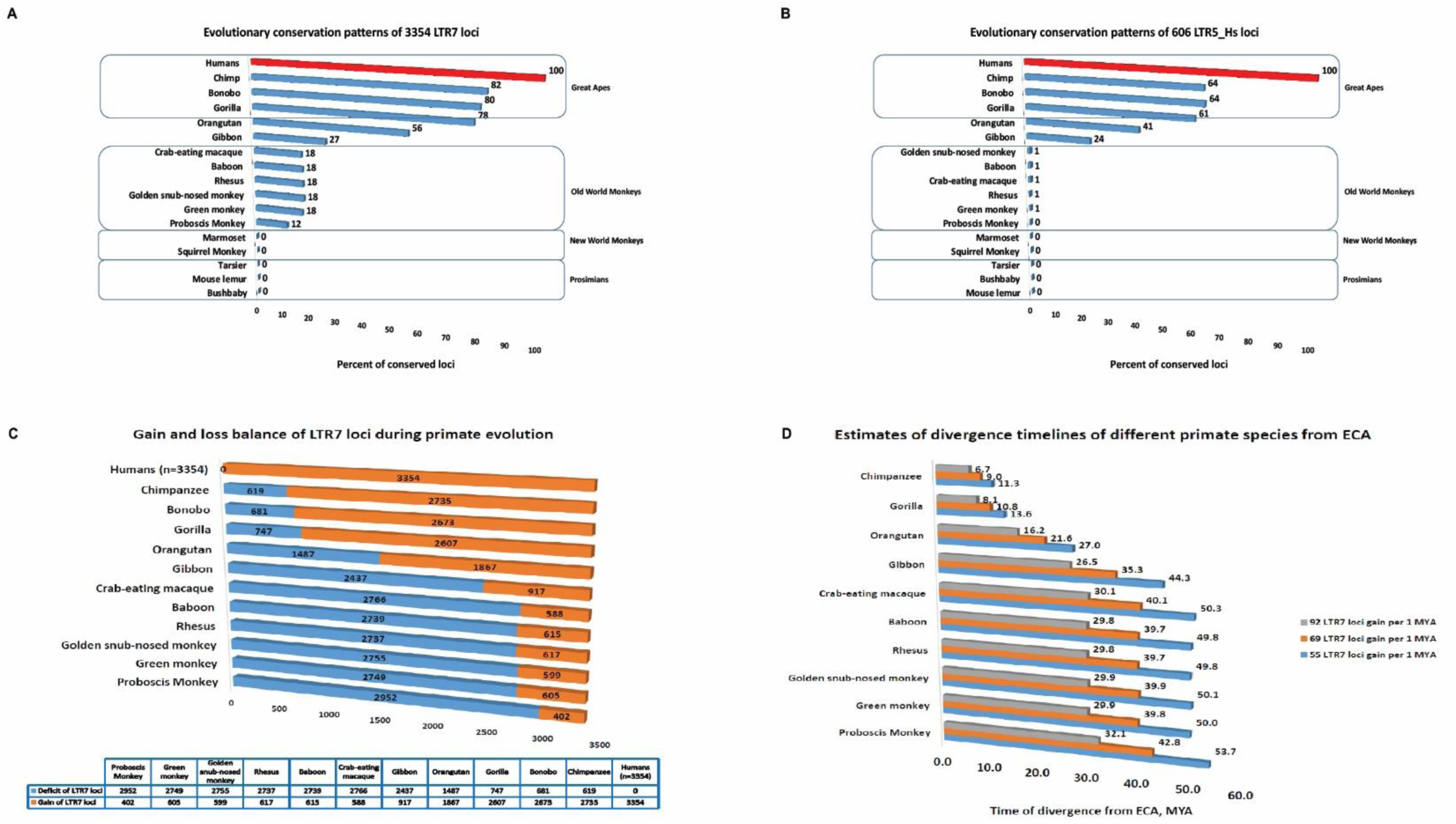
Evolutionary conservation analysis of regulatory LTR7 (A) and LTR5_Hs (B) loci and a model of species-specific expansion of regulatory LTR7 loci during primate evolution (C) and its putative associations with species segregation processes (D). The model is presented as a balance of gains and losses of LTR7 in genome of each primate species (C). Estimates of divergence timelines of different primate species from ECA based on estimated numbers of LTR7 loci acquisition per 1 MYA (D).

Interestingly, timeline sequences of quantitative expansion of both LTR7 and LTR5_Hs loci during primate evolution appear to replicate the consensus evolutionary sequence of increasing cognitive and behavioral complexities of NHP (**Figure 1**), which seems particularly striking for LTR7 loci (**Figure 1A**). This hypothesis was extended further by building a model of a putative species-specific expansion of regulatory LTR7 loci during primate evolution presented for genomes of eleven NHP as relative gains (a number of highly conserved LTR7 loci identified in a genome) and losses (a deficit of highly conserved LT7 loci with regard to human genome) of LTR7 loci vis-a-vis Modern Human’s genome (**Figure 1C**). A notable feature of this model is apparently similar numbers of gains and losses of LTR7 loci independently estimated for genomes of five Old World Monkeys’ species, providing a baseline for estimates of numbers of LTR7 loci gains per MYA during primate evolution (**Figure 1D**). Based on these estimates tailored to a presumed timeline of Old World Monkeys’ segregation from ECA, a hypothetical model defining species segregation timelines could be built, which reflect putative associations of LTR7 loci acquisitions in primate genomes with timelines of species segregation processes during primate evolution (**Figure 1D**).

Recent investigations of fine molecular structures of LTR7 elements and their genetic and regulatory distinctions demonstrated that LTR7 sequences represent a complex polyphyletic group composed of at least eight monophyletic subfamilies (Carter et al., 2022). Next, sequence conservation analyses of each individual monophyletic subfamilies of LTR7 loci in genomes of sixteen NHP have been performed. It has been observed that highly conserved sequences of all eleven monophyletic LTR7 subfamilies are present in genomes of all Old World Monkeys’ species analyzed in this contribution as well as in genomes of Gibbon, Orangutan, Gorilla, Bonobo, and Chimpanzee (**Figure 2**). These observations suggest that diversification of LTR7 loci into distinct genetic and regulatory subfamilies may have occurred early during primate evolution and subsequent cycles of LTR7 expansion appear to maintain this diversity.

**Figure 2.**
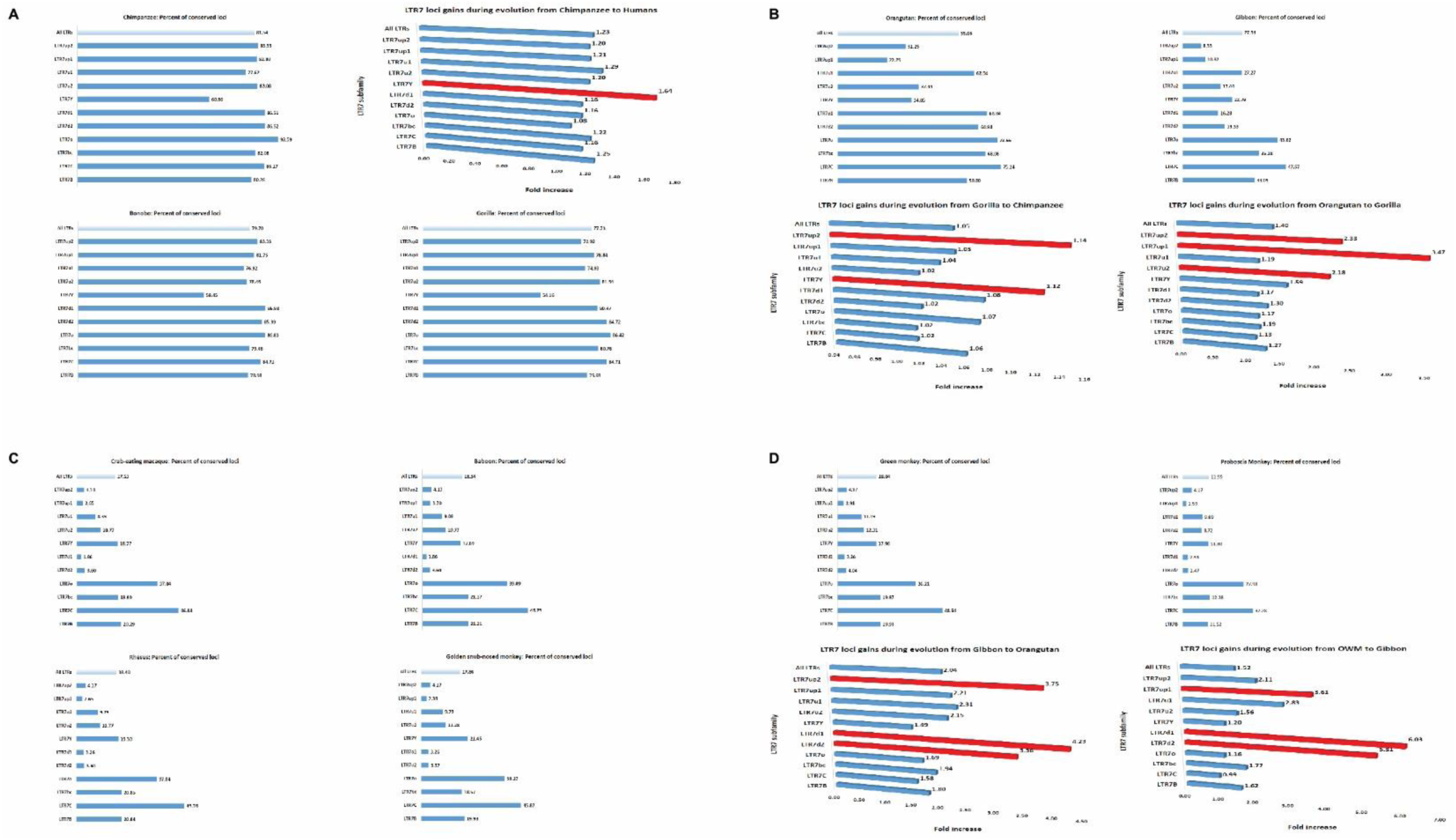
Granular evolutionary conservation analysis of eleven regulatory LTR7 subfamilies identifies subfamilies that are most rapidly expanding at different stages of species segregation during primate evolution. Numbers of highly conserved LTR7 loci of each subfamily were determined for each primate species and reported as the percentage of corresponding loci residing in genomes of Modern Humans. Relative gains of corresponding LTR7 subfamilies were calculated as ratio of highly conserved loci in designated species. Red colored bars denote the most rapidly expanding LTR7 subfamilies at indicated evolutionary stages.

Notably, despite the large difference in numbers of LTR7 loci present in genomes of different NHP species, the overall balance amongst eleven different LR7 subfamilies appears substantially similar (**Figure 3**). A great degree of resemblance is particularly evident for evolutionary closely-related species (**Figure 3** and data not shown), which is exemplified by high correlation coefficient values of LTR7 subfamilies’ abundance profiles estimated in pair-wise comparisons between Humans and Chimpanzee (r = 0.970), Bonobo (r = 0.968), and Gorilla (r = 0.958). Analyses of degrees of resemblance of LTR7 subfamilies abundance profiles amongst genomes of Great Apes revealed nearly identical arrangements of LTR7 subfamilies composition: Chimpanzee and Bonobo comparison yielded a pair-wise correlation coefficient of 0.999, while Chimpanzee and Gorilla comparison resulted in a pair-wise correlation coefficient of 0.998.

**Figure 3.**
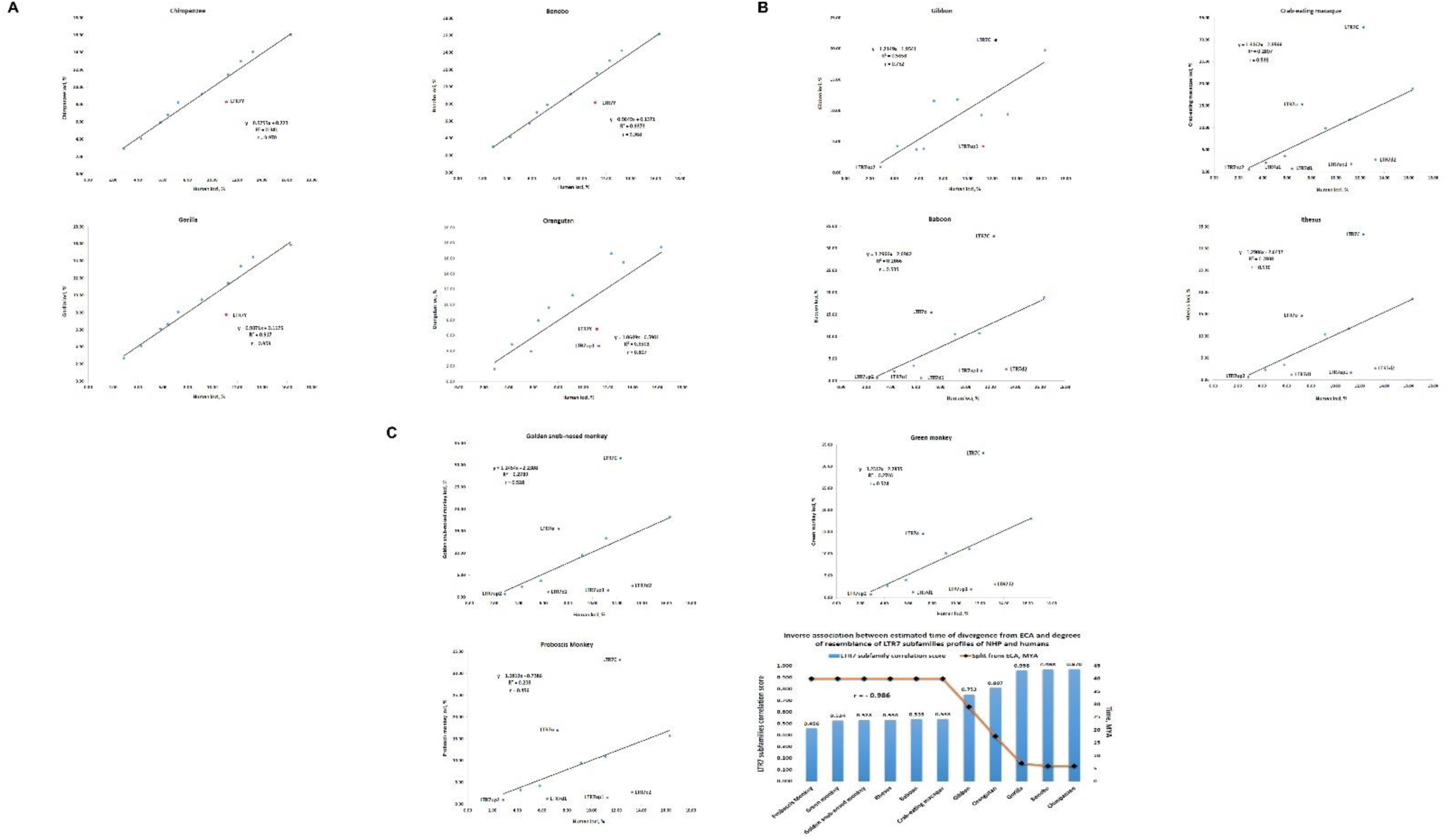
Correlation analyses of the granular evolutionary conservation patterns of regulatory LTR7 subfamilies. Granular evolutionary conservation patterns are presented as correlation plots of abundance profiles of LTR7 subfamilies in genomes of each non-human primate species vis-a-vis Modern Humans. Abundance profiles of LTR7 subfamilies were independently determined for each species and reported as percentage of loci of a given subfamily in a species’ genome. Note strikingly similar correlation coefficients for closely-related primate species which is gradually decreasing with increasing distance of species segregation. The inverse association between the estimated times of divergence from ECA and degrees of resemblance of LTR7 subfamilies abundance profiles of NHP species and Modern Humans (C; bottom right panel).

Similarly, inter-species correlation coefficients for pair-wise comparisons between different Old World Monkeys species consistently exceeded values of 0.99. When the abundance profile of LTR7 subfamilies in genomes of Modern Humans was utilized as a reference, a gradual decline of correlation coefficient values of LTR7 subfamilies’ abundance profiles estimated in pair-wise comparisons between Humans and more distant NHP species has been observed (**Figure 3**). A graphical summary of these findings reported in the **Figure 3C** illustrates the inverse association pattern between estimated times of divergence from ECA and degrees of resemblance of LTR7 subfamilies abundance profiles in genomes of NHP and Modern Humans (**Figure 3C**).

### Evolutionary conservation and divergence patterns of human-specific insertions of HERVH promoter LTR7 and HERVK promoter LTR5_Hs

Results of LTR loci sequence conservation analyses indicate that hundreds of LTRs in genomes of our closest evolutionary relatives, Chimpanzee and Bonobo, have DNA sequences divergent by more than 5% from orthologous sequences in genomes of Modern Humans. It was of interest to determine how many of fixed non-polymorphic LTR loci identified in human genome could be defined as human-specific compared to both Chimpanzee and Bonobo genomes. Setting a selection threshold at <10% of sequence identity, there are 175 LTR7 sequences and 176 LTR5_Hs sequences that could be classified as human-specific loci (**Figure 4**). Notably, a majority of candidate human-specific LTR7 (149/175; 85%) and LTR5_Hs (139/176; 80%) loci could be classified as bona fide human-specific insertions because they did not intersect any chains in genomes of both Chimpanzee and Bonobo.

**Figure 4.**
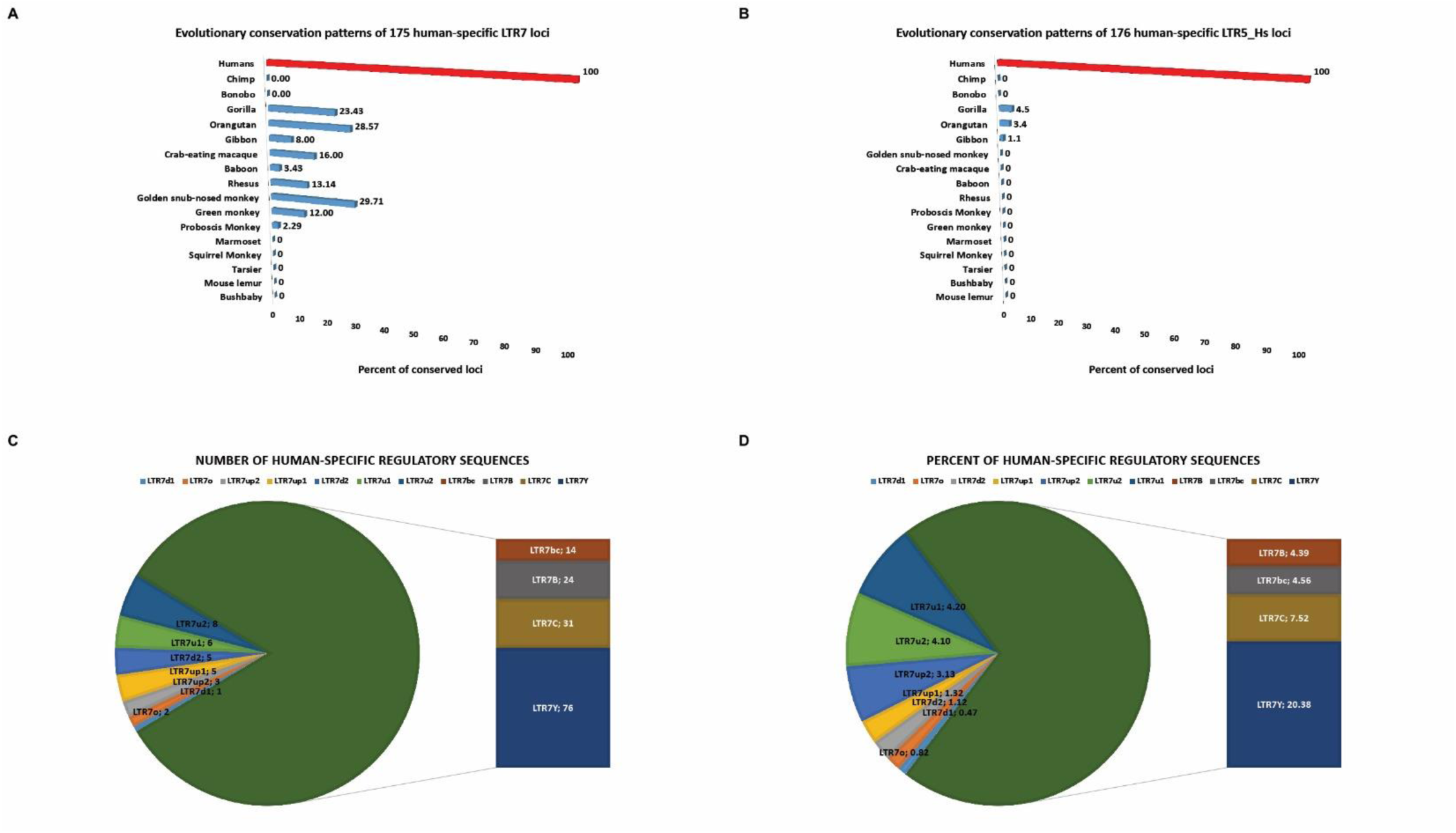
Evolutionary conservation patterns of human-specific LTR7 (A) and LTR5_Hs (B) loci in primates’ genomes identify highly-conserved LTR sequences in genomes of NHP representing remnants of past retroviral expansion events during primate evolution. Granular analyses of numbers (C) and percentages (D) of human-specific LTR7 elements’ distribution amongst eleven monophyletic LTR7 subfamilies identify LTR7Y subfamily as a dominant source of human-specific insertions in genomes of Modern Humans. In (D), percentages of human-specific loci for each LTR7 subfamily are reported calculated as fractions of all LTR7 sequences of corresponding subfamily.

However, evolutionary conservation analyses revealed that nearly half (84/175; 48%) of human-specific LTR7 loci could be mapped as highly conserved sequences present in genomes of Old World Monkeys, Gibbon, Orangutan, and Gorilla (**Figure 4A**), suggesting that these LTR7 loci were not retained in genomes of Chimpanzee and Bonobo but preserved in genomes of Modern Humans. In contrast, a much smaller fraction (8/176; 4.5%) of human-specific LTR5_Hs loci could be mapped as highly conserved sequences in genomes of Gibbon (2 loci), Orangutan (5 loci), and Gorilla (7 loci) (**Figure 4B**).

Granular analyses of human-specific regulatory sequences amongst different LTR7 subfamilies revealed that the LTR7Y subfamily harbors nearly half (76/175; 43%) of all human-specific LTR7 loci (**Figure 4C**). Overall, 20.4% of all LTRY sequences present in human genome could be classified as human-specific LTR7 loci (**Figure 4D**), while other LTR7 subfamilies harbor much smaller fractions of sequences defined as human-specific loci (**Figure 4D**).

### Inference of potential phenotypic impacts of human-specific insertions of HERVH promoter LTR7 and HERVK promoter LTR5_Hs

To infer potential biological functions of human-specific LTRs operating as distal regulatory loci, the GREAT algorithm was employed (Methods) to define the genome-wide connectivity maps of human-specific LTRs and their putative target genes. Concurrently with identification of putative regulatory target genes of human-specific LTRs, the GREAT algorithm performs stringent statistical enrichment analyses of functional annotations of identified genes, thus enabling the inference of potential biological significance of interrogated GRNs. Concurrently, a comprehensive panel of GSEA was executed employing the Enrichr bioinformatics platform (Methods) implemented on ∼30 genomics and proteomics databases (Methods) by imputing candidate down-stream regulatory target genes of human-specific LTRs (**Figure 5; Supplementary Figures S1 & S2**).

**Figure 5.**
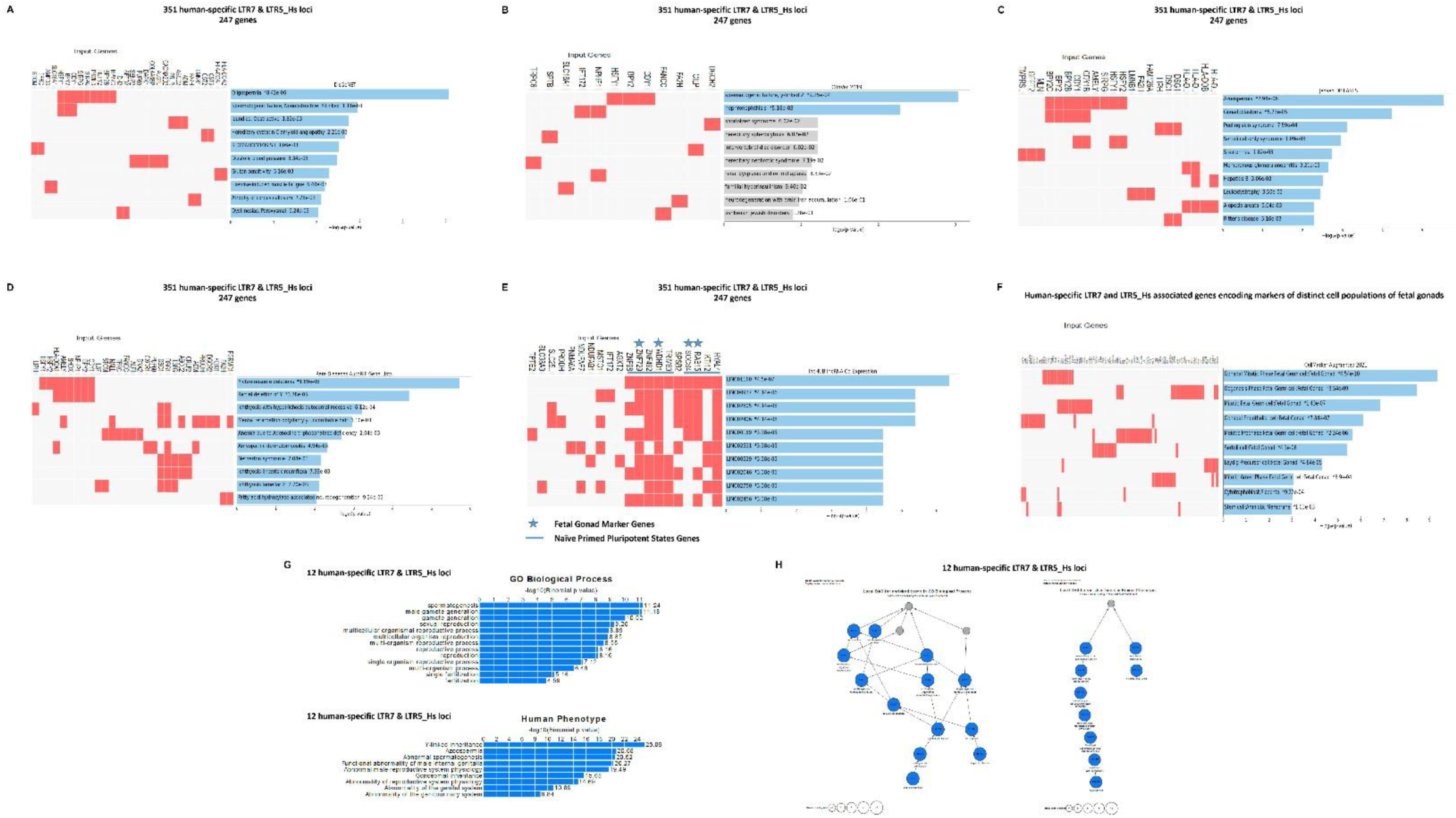
Inference of putative phenotypic impacts of human-specific LTR7 and LTR5_Hs loci based on GREAT algorithm-guided identification and functional annotations of proximity placement-linked genes. GSEA of 247 genes linked by GREAT with 351 human-specific LTR7 and LTR5_Hs using the Enrichr bioinformatics platform (A – F) and the GREAT algorithm-guided identification of 12 human-specific LTR7 elements (6 loci) and LTR5_Hs elements (6 loci) most significantly enriched in multiple GO Biological Process and Human Phenotype Ontology categories (G – H).

Consistent with documented biological roles of LTR7 and LTR5_Hs regulatory loci in establishment and maintenance of stemness and pluripotency phenotypes (Introduction), GSEA of 247 genes linked by GREAT with 351 human-specific LTR7 and LTR5_Hs revealed that a majority of significantly enriched records (70% of top 10 enriched records) represents genes associated with naïve and primed pluripotent states (GSEA of the database of human RNAseq GEO signatures; **Supplementary Figure S1**). Notable enrichment patterns of potential biological interest and translational significance were observed during the GSEA of several genomics databases of human common and rare diseases (**Figure 5**).

Top significantly enriched records identified by GSEA were Oligospermia (p = 8.42E-06; DisGeNET database); Spermatogenic Failure, Y-linked (p = 9.25E-04; ClinVar 2019 database), Azoospermia (p = 2.94E-06; Jensen Diseases database), Y chromosome deletions (p = 1.89E-06; Rare Diseases AutoRIF Gene Lists database) (**Figure 5**), suggesting that regulation of human spermatogenesis might be one of biologically important functions of genes under putative regulatory control of human-specific LTR7 and LTR5_Hs loci. Consistent with this idea, the GREAT algorithm analysis of 351 human-specific LTR7 and LTR5_Hs loci identified seven human genes (*BPY2; CDY1; DAZ2; HSFY1; RPS4Y2; SRY; LMNB1*) associated by the Human Phenotype Ontology database analysis (**Supplementary Table S1**) with phenotypes of Y-linked inheritance (HP:0001450; p = 3.92E-07); Abnormal male reproductive system physiology (HP:0012874; p = 1.65E-05); Azoospermia (HP:0000027; p = 3.87E-05); Abnormal spermatogenesis (HP:0008669; p = 5.22E-05); Functional abnormality of male internal genitalia (HP:0000025; p = 8.37E-05). Follow-up analytical experiments focused on human-specific LTR loci and genes linked with the listed above human phenotypes confirmed these observations. The GREAT algorithm-defined connectivity map of regulatory loci and target genes identified 12 human-specific sequences of LTR7 (6 loci) and LTR5_Hs (6 loci) that manifested statistically significant enrichments in GO Biological Process (13 significantly enriched records) and Human Phenotype Ontology (9 significantly enriched records) categories (**Figure 5; Supplementary Figure S1**).

GSEA of lncHUB lncRNA Co-Expression database defined 30 significantly enriched records of human long non-coding RNA molecules (lncRNAs) that are co-expressed in human tissues with a sub-sets of genes representing putative regulatory targets of human-specific LTRs (**Figure 5; Supplementary Figure S2**). Detailed investigation of co-expressed loci demonstrated that 8 of 10 genes co-expressed with top-scoring LNC01160 lncRNA (p = 4.5E-07; **Figure 5**) represent genes differential expression of which distinguishes Naïve and Primed pluripotent states, while four genes constitute genetic markers of fetal gonads (**Figure 5; Supplementary Figure S2**). Next, follow-up analyses were carried-out employing the CellMarker Augmented 2021 database of single-cell genomics-guided genetic markers of human and mouse cells comprising gene sets from the CellMarker database augmented with co-expression RNA-seq data from ARCHS4 (Enrichr). These analyses identified 88 genes representing putative regulatory targets of human-specific LTR7 and LTR5_Hs loci and comprising genetic markers of 12 distinct cell populations of fetal gonads (**Figure 5; Supplementary Figure S2**). Collectively, these observations suggest that genes implicated in development of human fetal gonads may represent regulatory targets of human-specific LTR7 and LTR5_Hs loci. Consistent with this hypothesis, expression of nearly three quarters of identified herein fetal gonad marker genes (64 of 88 genes; 73%) is significantly altered in human cells subjected to genetic and/or epigenetic targeting of LTR7/HERVH or LTR5_Hs loci (see below).

### GSEA of retroviral LTRs-linked genes revealed dominant enrichment patterns of physiological and pathological phenotypic traits affected by mammalian offspring survival (OS) genes associated with LTR7 and LTR5_Hs regulatory loci

DNA sequences derived from LTR7/HERVH and LTR5_Hs/HERVK retroviral insertions have been identified as one of significant sources of the evolutionary origin of human-specific regulatory sequences (HSRS), including transcription factor-binding sites (TFBS) for stemness state master regulators NANOG, OCT4, and SOX2 (Glinsky, 2015 – 2021). Since mammalian offspring survival (OS) genes have been implicated as one of the putative genomic regulatory targets of HSRS (Glinsky, 2020), it was of interest to determine whether mammalian OS genes are enriched amongst candidate regulatory targets of LTR7 and LTR5_Hs loci. Amongst 18,777 human genes comprising the GREAT database gene set (hg38 release of the human reference database), there are 2,413 mammalian OS genes defined as genes mutations of which have been associated by the MGI database search with premature death, embryonic lethality, as well as pre-, peri-, neo-, and post-natal lethality phenotypes of both complete and incomplete penetrance. Of these, a total of 562 mammalian OS genes has been identified as putative regulatory targets of LTR7 loci (**Table 1**), which represents a significant enrichment compared to expected by chance number of genes (p = 1.131E-10; 2-tail Fisher exact test). In contrast, the number of mammalian OS genes identified as putative regulatory targets of LTR5_Hs loci (126 genes) did not exceed the enrichment significance threshold level (**Table 1**). Detailed analyses of the enrichment levels’ distribution of mammalian OS genes amongst different LTR7 subfamilies demonstrate that proportions of OS genes appear enriched amongst putative LTR7 regulatory targets of various LTR7 subfamilies (**Table 2**). These observations suggest that mammalian OS genes seem to remain one of favorite regulatory targets throughout ∼30 MYA of the divergent evolution of LTR7 loci.

**Table 1.**
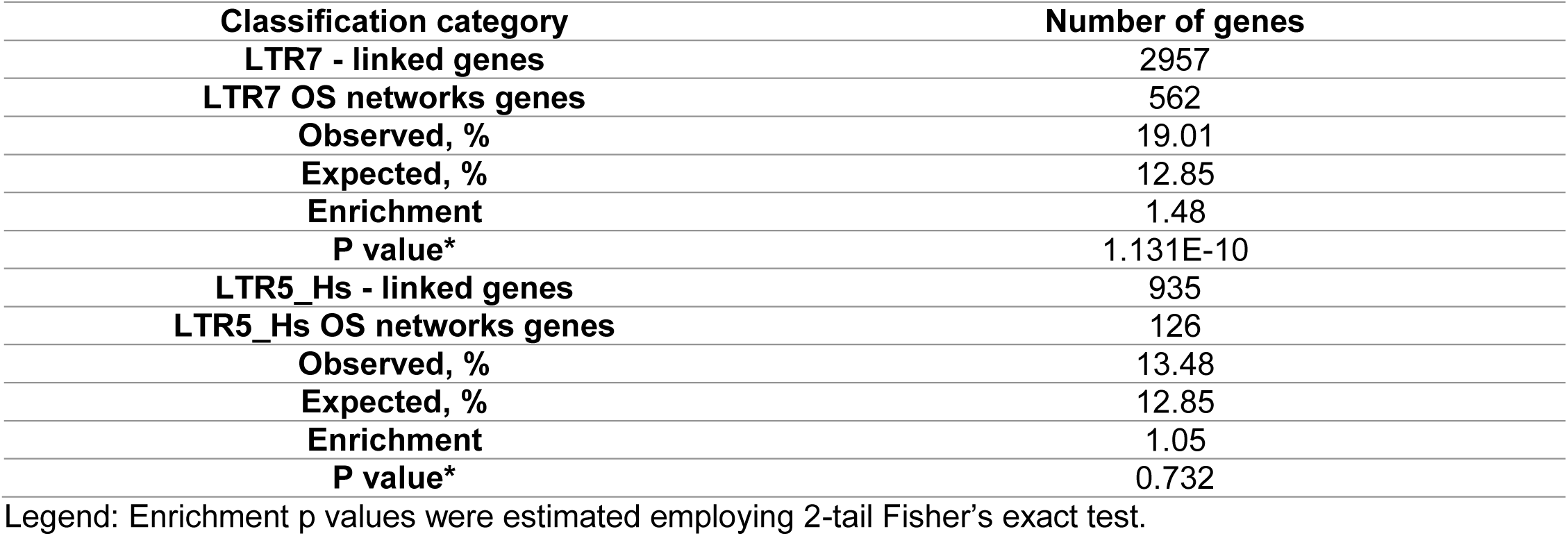
Enrichment patterns of mammalian offspring survival (OS) genes linked with LTR7 and LTR5_Hs loci.

**Table 2.**
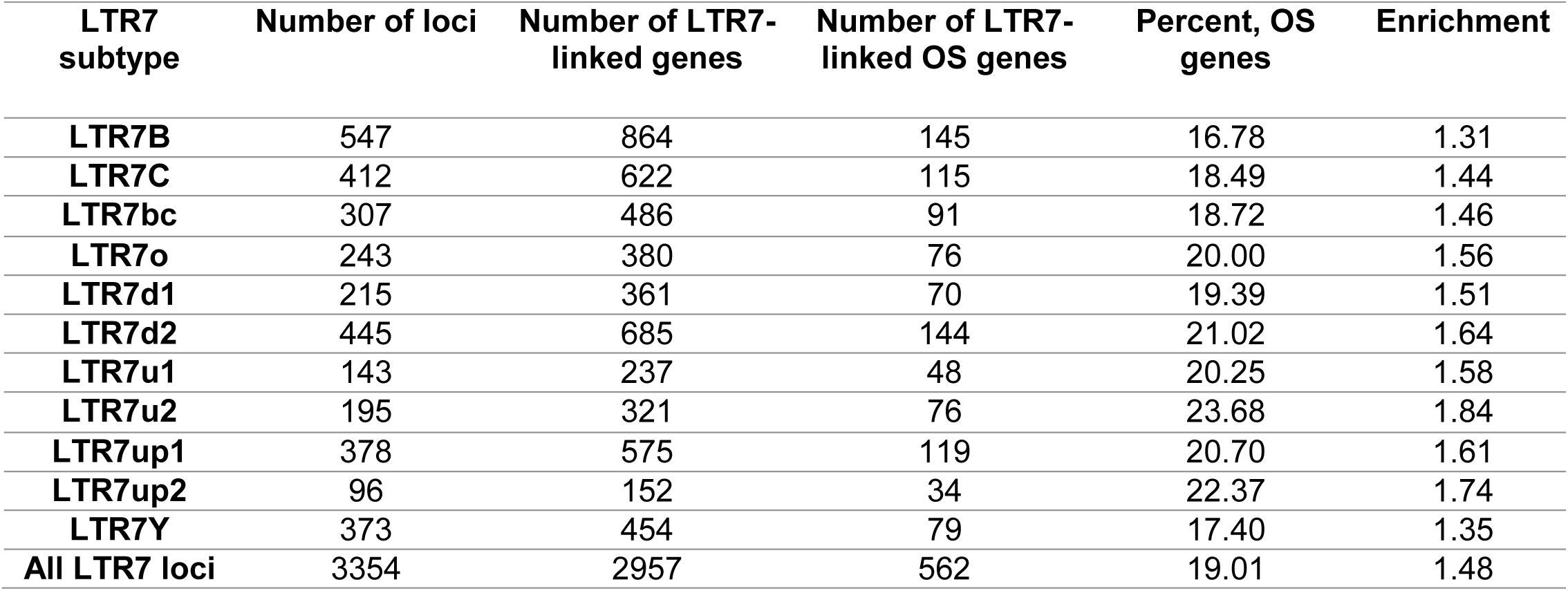
Association patterns of distinct subfamilies of regulatory LTR7 loci with offspring survival (OS) genes.

It was of interest to determine whether mammalian OS genes may have the broader impacts on pathophysiology of Modern Humans extending beyond offspring survival phenotypes. To this end, independent GSEA were carried out on all genes defined as putative regulatory targets of LTR7 and LTR5_Hs loci and subsets of regulatory targets comprising mammalian OS genes and non-OS genes (**Table 3**). These analyses revealed clearly discernable dominant enrichment patterns of phenotypic traits affected by mammalian OS genes linked with LTR7 and LTR5_Hs regulatory loci across the large panel of genomics and proteomics databases reflecting a broad spectrum of human pathophysiology (**Table 3**). Enrichment patterns’ differences were particularly notable for GSEA of databases of human common and rare diseases as well as Human Phenotype Ontology database, suggesting that mammalian OS genes may have significant impacts on development of phenotypic traits and pathological conditions of Modern Humans. Significantly, clearly-defined heritability features seem apparent for a majority of LTR7-linked mammalian OS genes associated with 466 phenotypic traits reported in the Human Phenotype Ontology database because they are represented by either Autosomal dominant inheritance genes (HP: 0000006; p = 1.84E-20) or Autosomal recessive inheritance (HP: 0000007; p = 5.79E-10) genes (**Figure 6**). Three top-ranked phenotypic traits captured by GSEA of 562 LTR7-linked mammalian OS genes employing the DisGeNET database of human diseases (Carcinogenesis; p = 1.45E-42; Intellectual disability; p = 2.66E-34; Neoplasm metastasis; p = 1.32E-32) appear associated with overlapping networks of genes (**Figure 6**), perhaps, reflecting yet poorly understood common mechanistic features for malignant neoplasms and brain disorders. Consistent with this hypothesis, a qualitatively similar genotype-phenotype association patterns were documented by GSEA of 126 LTR5_Hs-linked mammalian OS genes (**Figure 6**).

**Figure 6.**
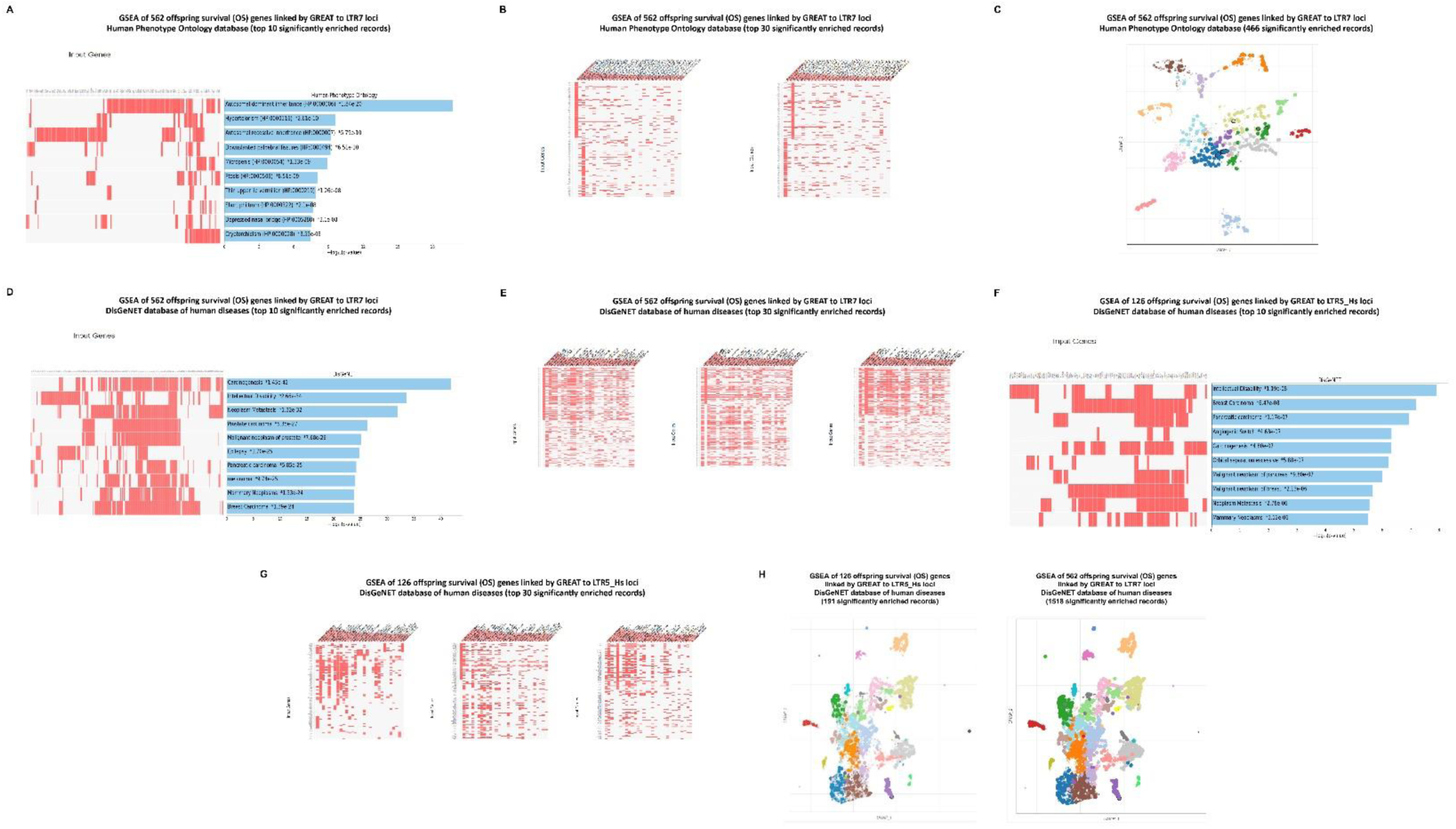
Analysis of phenotypic impacts of LTR7 - and LTR5_Hs - linked mammalian offspring survival (OS) genes revealed by GSEA of the Human Phenotype Ontology database (A – C) and the DisGeNET database of human diseases (D – H). The scatterplots in (C) and (H) are organized so that similar gene sets are clustered together. Larger, darker, black-outlined points represent significantly enriched terms. Clusters are computed using the Leiden algorithm. Points are plotted on the first two UMAP dimensions. In web-based software settings, hovering over points will display the associated gene set name and the p-value. Reader may have to zoom in using the toolbar next to the plot in order to see details in densely-populated clusters. Plots can also be downloaded as a svg using the save function on the toolbar.

**Table 3.** Dominant enrichment patterns of phenotypic traits affected by offspring survival (OS) genes linked with LTR7 and LTR5_Hs regulatory loci.

| **Classification category** | **LTR7** | **LTR7**<br>**OS**<br>**genes** | **LTR7**<br>**non-OS**<br>**genes** | **LTR5_Hs** | **LTR5_Hs**<br>**OS**<br>**genes** | **LTR5_Hs**<br>**non-OS**<br>**genes** |
| --- | --- | --- | --- | --- | --- | --- |
| **Number of associated genes** | 2957 | 562 | 2395 | 935 | 126 | 809 |
| **Significantly enriched records of phenotypic traits identified by the GSEA of genomic databases** |  |  |  |  |  |  |
| **DisGeNET database** | 72 | 1518 | 0 | 1 | 191 | 1 |
| **PanglaoDB Augmented 2021** | 70 | 67 | 33 | 3 | 4 | 0 |
| **CellMarker Augmented 2021** | 203 | 181 | 83 | 3 | 0 | 2 |
| **Azimuth Cell Types 2021** | 89 | 17 | 47 | 0 | 0 | 0 |
| **ARCHS4 Human Tissues** | 36 | 37 | 24 | 2 | 11 | 0 |
| **Human Gene Atlas** | 2 | 10 | 0 | 0 | 1 | 0 |
| **Allen Brain Atlas database of up-regulated genes** | 420 | 506 | 43 | 0 | 0 | 0 |
| **Allen Brain Atlas database of down-regulated genes** | 101 | 230 | 5 | 0 | 1 | 0 |
| **GTEx tissues' expression database of up-regulated genes** | 334 | 583 | 129 | 0 | 48 | 0 |
| **GTEx tissues' expression database of down-regulated genes** | 98 | 329 | 56 | 0 | 80 | 0 |
| **Reactome 2016** | 3 | 50 | 0 | 2 | 77 | 1 |
| **WikiPathway 2021 Human** | 1 | 68 | 0 | 0 | 20 | 0 |
| **KEGG 2021 Human** | 2 | 38 | 1 | 1 | 15 | 1 |
| **BioPlanet 2019** | 6 | 111 | 1 | 3 | 4 | 1 |
| **Jensen DISEASES database** | 40 | 75 | 15 | 4 | 6 | 3 |
| **MGI Mammalian Phenotype Level 4 2021** | 25 | 1253 | 0 | 0 | 207 | 0 |
| **Human Phenotype Ontology** | 3 | 466 | 0 | 0 | 26 | 0 |
| **GTEx Aging Signatures 2021** | 14 | 0 | 11 | 0 | 0 | 0 |
| **Disease Perturbations from GEO up-regulated genes** | 63 | 160 | 14 | 0 | 0 | 0 |
| **Disease Perturbations from GEO down-regulated genes** | 88 | 153 | 23 | 0 | 0 | 0 |
| **Significantly enriched records of phenotypic traits identified by the GSEA of human rare diseases databases** |  |  |  |  |  |  |
| **Rare Diseases GeneRIF Gene Lists** | 1 | 497 | 0 | 0 | 5 | 0 |
| **Rare Diseases AutoRIF Gene Lists** | 15 | 738 | 0 | 0 | 2 | 0 |
| **Rare Diseases AutoRIF ARCHS4 Predictions** | 57 | 161 | 0 | 0 | 0 | 0 |
| **Rare Diseases GeneRIF ARCHS4 Predictions** | 47 | 137 | 0 | 0 | 11 | 0 |
| **Significantly enriched records of phenotypic traits identified by the GSEA of targeted TF perturbations, TF PPI, and PPI Hub Proteins genomic databases** |  |  |  |  |  |  |
| **TF Perturbations Followed by Expression** | 368 | 704 | 76 | 4 | 26 | 1 |
| **Transcription Factor PPIs** | 0 | 86 | 0 | 0 | 0 | 0 |
| **PPI Hub Proteins** | 0 | 105 | 0 | 0 | 9 | 0 |

Legend: Gene Set Enrichment Analyses (GSEA) were carried out for each gene set and numbers of significantly enriched phenotypic traits defined at the adjusted p values < 0.05 were recorded. TF, transcription factors; PPI, protein-protein interactions; GEO, Gene Ontology Omnibus. Complete lists of phenotypic traits, associated genes, and corresponding statistical metrics for analyzed genomic databases are reported in the Supplement.

Exceptions from quantitatively dominant enrichment patterns of phenotypic associations of mammalian OS genes were noted for a database reporting genetic markers of human tissues (ARCHS4 Human Tissues database; **Table 3**) and several databases of cell type-specific markers (PanglaoDB Augmented 2021; CellMarker Augmented 2021; and Azimuth Cell Types 2021 databases; **Table 3**). These findings demonstrate consistent tissue- and cell-type specific expression profiles of both mammalian OS genes and non-OS genes comprising putative regulatory targets of LTR7 loci, perhaps, reflecting their contributions to functions of a broad spectrum of differentiated cells in human body (**Table 3**). In contrast, GSEA of LTR7-linked genes focused on the GTEx Aging Signatures 2021 database revealed that significantly-associated traits of numerous human aging tissues (nerve; small intestine; uterus; beast; brain; lung; blood vessels) appear linked with non-OS genes (**Table 3; Supplementary Table S2**).

Genetic targeting of 704 transcription factor (TF) - coding genes affected expression of large numbers of mammalian OS genes comprising putative regulatory targets of LTR7 loci, which significantly exceeded numbers of targeted genes expected by chance (**Table 3; Supplementary Table S2**).

These observations indicate that reported herein LTR7-linked genes may function in human cells as down-stream targets of genomic regulatory networks governed by hundreds of TFs. Intriguingly, protein products of many mammalian OS genes comprising putative regulatory targets of LTR7 loci were identified as partners of protein-protein interactions (PPI) of at least 86 TFs and 105 PPI Hub proteins, which are proteins that interact with protein products encoded by at least 50 other genes. These findings support the hypothesis that engagements in PPI of the multimolecular complexes operating in human cells may represent an important mechanistic vector of biological activities of mammalian OS genes comprising putative regulatory targets of LTR7 loci.

### Identification and characterization of retroviral LTR7 and LTR5_Hs loci associated with genes regulating synaptic transmission and protein-protein interactions at synapses

GSEA of genomic databases focused on gene expression signatures of human tissues and cell types revealed a clear prevalence of enrichment records related to brain and CNS functions amongst significantly enriched phenotypic traits affected by genes comprising putative regulatory targets of LTR7 loci (**Supplementary Table S2**). For example, records of cell types and tissues related to brain and CNS functions constitute 55% and 75% of top 20 significantly enriched records identified by GSEA of the single-cell sequencing PanglaoDB Augmented 2021 and ARCHS4 Human Tissues databases, respectively. Strikingly, all top 20 significantly enriched records (**Supplementary Table S2**) identified by GSEA of the GTEx human tissues’ expression database of up-regulated genes and single-cell genomics-guided Azimuth Cell Types 2021 database reference either brain samples (GTEx database) or different highly specialized types of GABAergic and Glutamatergic neurons (Azimuth database). Overall, 82 of 89 (92%) of all significantly enriched records identified by GSEA of the Azimuth Cell Types 2021 database represent different highly specialized types of neurons (**Supplementary Table S2**). Significantly enriched records of cell types identified by GSEA of the single-cell sequencing PanglaoDB Augmented 2021 database represent cells of distinsct neurodevelopmental stages and morphologically diverse cell types residing and functioning in human brain, which include Neural Stem/Precursor cells, Radial Glia cells, Bergman Glia cells, Pyramidal cells, Tanycytes, Immature neurons, Interneurons, Trigeminal neurons, GABAergic neurons, and Glutamatergic neurons (**Supplementary Table S2**). Collectively, these observations indicate that one of the important biological functions of genes comprising putative regulatory targets of LTR7 loci is contribution to development and functions of the human brain. Consistent with this hypothesis, GSEA of the Allen Brain Atlas database identified 521 significantly enriched records of different human brain regions harboring expression signatures of both up-regulated (420 brain regions) and down-regulated (101 brain regions) genes comprising putative LTR7 regulatory targets (**Supplementary Table S2**).

Based on these findings, it was of interest to determine whether genes that play the essential biological role in brain functions are enriched amongst putative LTR7-taget genes. To test this hypothesis, records of 355 genes defined by the Reactome database as genes whose products regulate the synaptic transmission and are engaged in protein-protein interactions at synapses (collectively designated here synaptic transmission networks’ genes) were retrieved and analyzed. It has been determined that there are 87 synaptic transmission networks’ genes comprising putative regulatory targets of LTR7 loci (**Table 4**), which represents the significant enrichment compared to the expected by chance value (p = 0.01; 2-tail Fisher’s exact test). In contrast, there were only 19 synaptic networks’ genes among genes comprising putative regulatory targets of LTR5_Hs loci (**Table 4**), which corresponds to the expected by chance value.

**Table 4.** Enrichment patterns of synaptic networks genes linked with LTR7 and LTR5_Hs loci.

| **Classification category** | **Number of genes** |
| --- | --- |
| **LTR7 - linked genes** | 2957 |
| **LTR7 Synaptic networks genes** | 87 |
| **Observed, %** | 2.94 |
| **Expected, %** | 1.89 |
| **Enrichment** | 1.56 |
| **P value\*** | 0.0109 |
| **LTR5_Hs - linked genes** | 935 |
| **LTR5_Hs Synaptic networks genes** | 19 |
| **Observed, %** | 2.03 |
| **Expected, %** | 1.89 |
| **Enrichment** | 1.07 |
| **P value\*** | 1 |

Legend: Enrichment p values were estimated employing 2-tail Fisher's exact test.

Next, the assessments were made of regulatory loci/target genes connectivity patterns with respect to synaptic transmission networks’ genes for individual LTR7 subfamilies. To this end, 220 LTR7 loci linked to 87 synaptic transmission networks’ genes by the GREAT algorithm (**Table 5; Supplementary Table S3**) were retrieved, all genes comprising their putative regulatory targets were identified, and numbers (fractions) of associated synaptic transmission networks’ genes were determined for each LTR7 subfamily (**Table 5**). It has been determined that all LTR7 subfamilies appear to manifest putative regulatory links to synaptic transmission networks’ genes (**Table 5**), suggesting that observed associations between LTR7 loci and synaptic transmission networks’ genes remain relatively constant during primate evolution. Notably, when a hypothetical genomic regulatory network connecting LTR7 loci and synaptic transmission networks’ genes was interrogated using the GREAT algorithm, Gene Ontology analyses identified numerous highly significantly enriched phenotypic traits affected by synaptic transmission networks’ genes linked with LTR7 loci (**Figure 7**). For example, analysis of GO Cellular Component database revealed 46 significantly enriched terms, while GO Molecular Function database identified 64 significantly enriched terms and GO Biological Process database defined 146 significantly enriched records (**Figure 7; Supplementary Table S3**). In contrast, the GREAT algorithm identified a single significantly enriched employing Human Phenotype ontology database, namely Autism (Binominal FDR q value = 1.313E-15).

**Figure 7.**
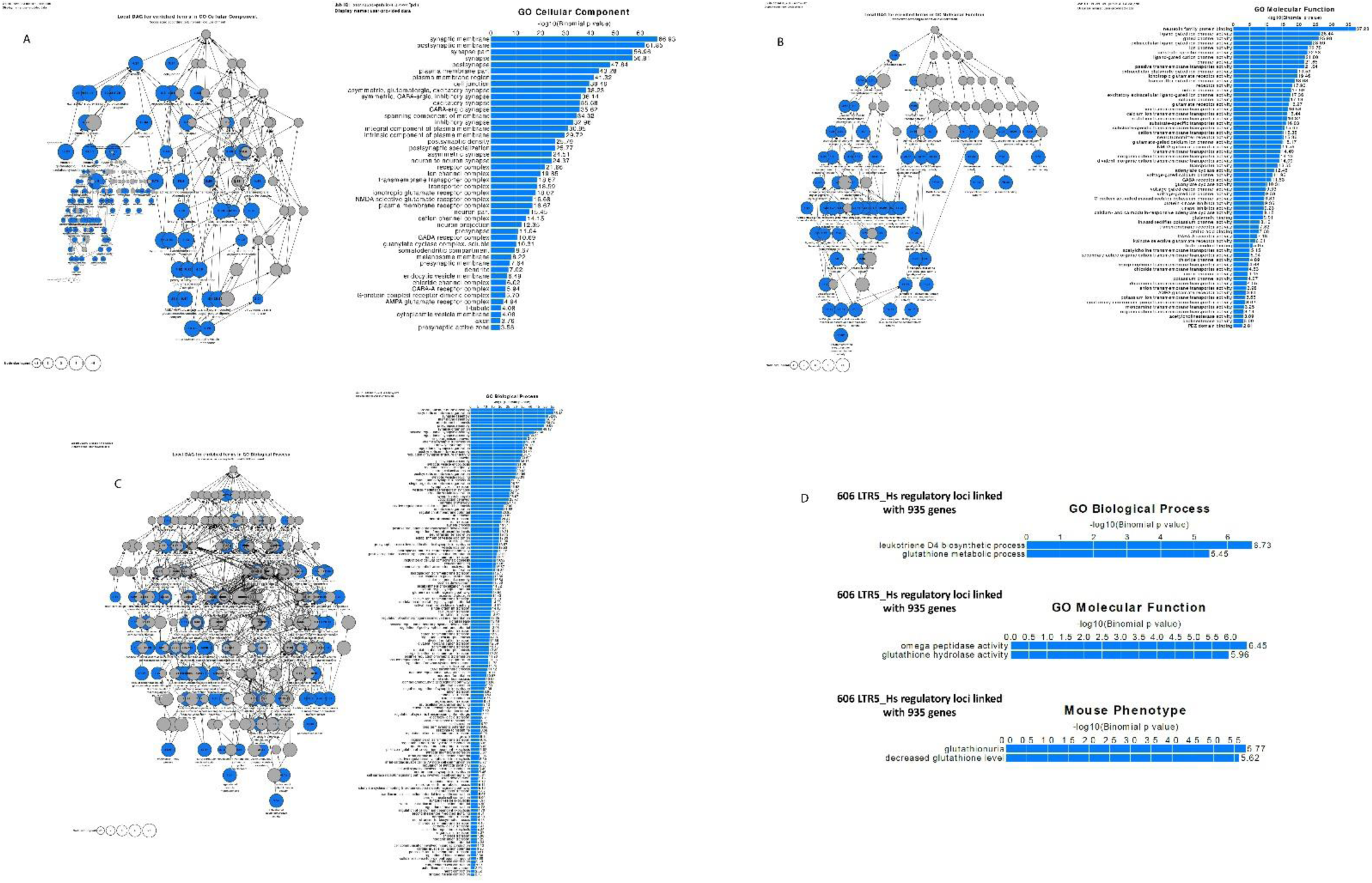
Identification and characterization LTR7 regulatory loci linked with genes whose products affect transmission across synapses and protein-protein interactions at synapses. Potential phenotypic impacts of 235 human regulatory LTR7 loci linked by the GREAT algorithm with 170 down-stream target genes revealed by interrogation of GO Cellular Component database (panel A; 46 significant terms), GO Molecular Function database (panel B; 64 significant terms), and GO Biological Process database (panel C; 146 significant terms). In contrast, analyses of 606 LTR5_Hs regulatory loci linked by GREAT with 935 gene identify 2 significantly enriched terms in GO Biological Process, GO Molecular Function, and Mouse Phenotype databases. Analyses were carried out using the GREAT algorithm designed to predict functions of cis-regulatory regions. Genomic coordinates of interrogated LTRs were based on the hg38 release of the human reference genome database.

**Table 5.** LTR7 regulatory loci linked with genes mediating transmission across synapses and protein-protein interactions at synapses.

| **LTR7 subfamily** | **Number of loci\*** | **Number of linked genes** | **Number of synaptic transmission network genes\*\*** | **Percent of synaptic transmission network genes** |
| --- | --- | --- | --- | --- |
| **LTR7B** | 28 | 37 | 22 | 59.5 |
| **LTR7C** | 24 | 38 | 19 | 50.0 |
| **LTR7bc** | 29 | 46 | 26 | 56.5 |
| **LTR7o** | 23 | 34 | 18 | 52.9 |
| **LTR7d1** | 15 | 26 | 15 | 57.7 |
| **LTR7d2** | 39 | 60 | 31 | 51.7 |
| **LTR7u1** | 10 | 14 | 9 | 64.3 |
| **LTR7u2** | 13 | 21 | 12 | 57.1 |
| **LTR7up1** | 23 | 36 | 18 | 50.0 |
| **LTR7up2** | 5 | 8 | 4 | 50.0 |
| **LTR7Y** | 11 | 18 | 10 | 55.6 |
| **All LTR7 loci** | 220 | 170 | 87 | 51.2 |

Legend: \* number of individual non-redundant LTR7 regulatory loci; in total, there are 235 regulatory loci linked with 170 human genes by the GREAT algorithm, including synaptic transmission networks' genes. \*\*Reported results are based on the Reactome database analysis of 355 synaptic transmission network genes.

Numerous significantly enriched records were identified by GSEA of Mouse Phenotype (110 significant terms) and Mouse Phenotype Single KO (97 significant terms) databases (**Figure 8**). These findings together with reported herein regulatory connectivity maps of HERV’s LTRs and their putative target genes provide readily available well-characterized mouse models for experimental interrogations of postulated causal regulatory effects of HERVs LTRs on specific target genes and respective phenotypes. For example, visualization of significantly enriched records identified by GO Molecular Function analysis at different stages of primate evolution depicts activities of the kainate selective glutamate receptor and the AMPA glutamate receptor as potentially important biological functions during the evolutionary transition period from Great Apes to Humans (**Figure 8**).

**Figure 8.**
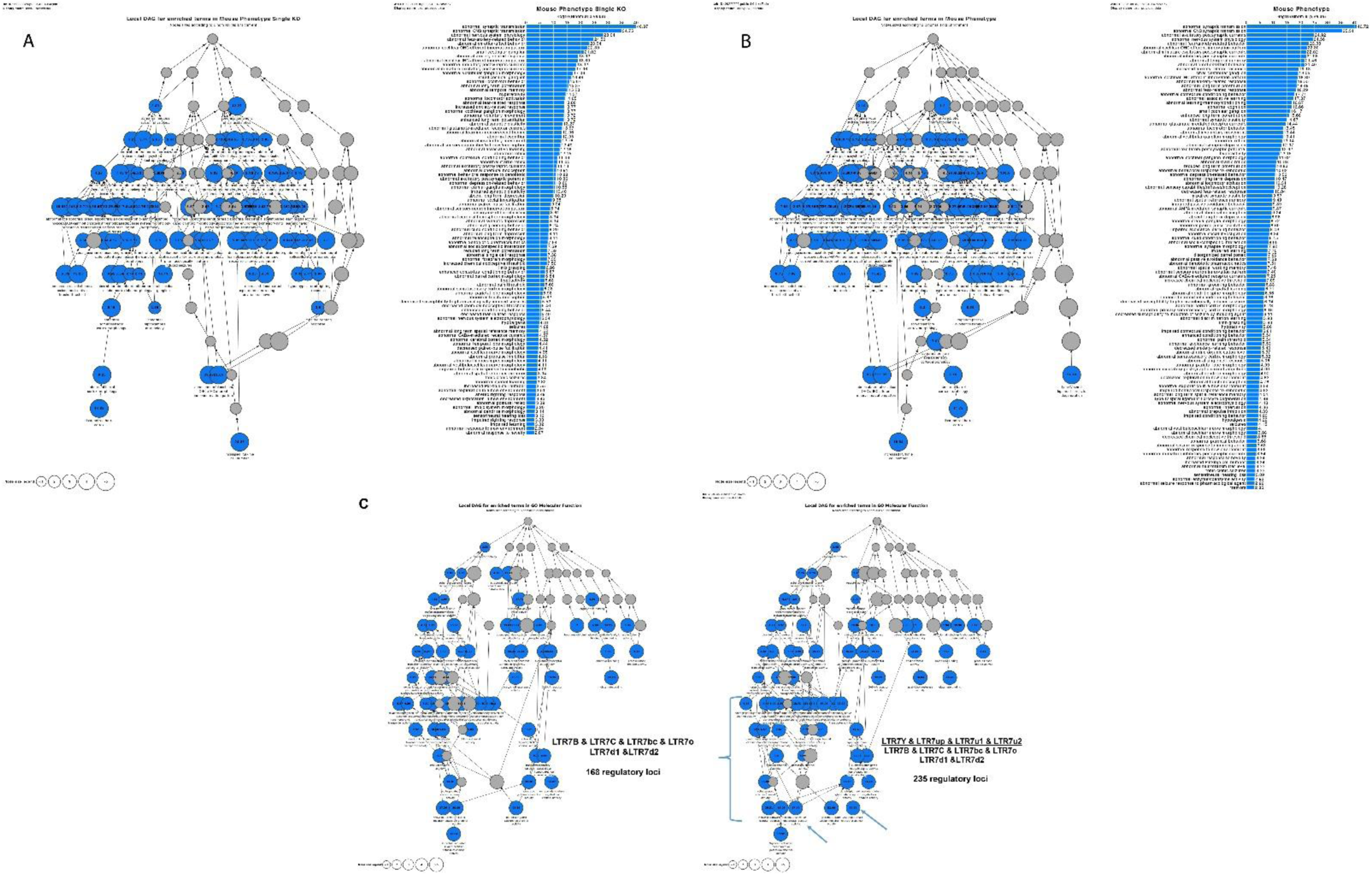
Potential phenotypic impacts of 235 regulatory LTR7 loci revealed by interrogation of Mouse Phenotype Single KO database (panel A; 97 significant terms) and Mouse Phenotype database (panel B; 110 significant terms). Differential GO Molecular Function analyses of genetically and evolutionary distinct subfamilies of human regulatory LTR loci (panel C) comprising 168 and 235 LTR7 loci designed to highlight LTR7-linked phenotypic traits enriched in humans after separation of chimpanzee and human lineages (depicted by arrows are the kainate selective glutamate receptor activity and AMPA glutamate receptor activity). Analyses were carried out using the GREAT algorithm designed to predict functions of cis-regulatory regions. Genomics coordinates of LTR7 loci were based on the hg38 release of the human reference genome database.

Analyses of evolutionary dynamics of connectivity patterns between candidate regulatory LTRs and down-stream target genes amongst specific functional and/or morphological categories suggest that dominant LTR expansion patterns during evolution of Great Ape consist of linking newly emerged regulatory LTR loci to either new down-stream target gene(s) or genes already integrated into LTR regulatory networks, which are contributing to and/or engaged within a specific phenotypic trait that already being targeted for evolutionary innovations at the earlier stages of primate evolution.

### Identification and characterization of high-fidelity genetic regulatory targets of LTR7 and LTR5_Hs loci

Inferences of potential phenotypic impacts of LT7 and LTR5_Hs loci on physiological traits and pathological conditions of Modern Humans were based on GSEA-guided assessments of documented biological functions and morphological features of genes comprising putative regulatory of LTR loci identified by the GREAT algorithm. To extend this line of inquiry further, it was of interest to determine what fraction of candidate LTR regulatory target genes constitutes bona fide transcriptional targets of LTRs in human cells defined as genes expression of which is altered following genetic targeting of LTRs. It has been determined that expression of a majority of genes (1570 of 2957; 53%) comprising candidate regulatory targets of LTR7 loci is significantly affected in hESC subjected to either targeted genetic interference with or epigenetic silencing of LTR7/HERVH loci (**Table 6**). Similarly, expression of a majority of genes (486 of 935 genes; 52%) identified as putative regulatory targets of LTR5_Hs loci is significantly altered in human teratocarcinoma cells following targeted epigenetic silencing or activation of LTR5_Hs loci (**Table 7**). Overall, expression of 1944 of 3515 genes (55%) comprising LTRs’ candidate regulatory targets was significantly affected following genetic or epigenetic manipulations of LTR7 and/or LTR5_Hs loci (**Table 8**). Based on these observations, it has been concluded that expression of a majority of genes identified herein as putative regulatory targets of LTR7 and/or LTR5_Hs loci appears altered in human cells following targeted genetic and/or epigenetic manipulations of LTR sequences, which is consistent with definition of these sets of genes as high-fidelity down-stream regulatory targets of LTR7 and LTR5_Hs loci. Interestingly, genes defined as high-fidelity down-stream regulatory targets of both LTR7 and LTR5_Hs loci represent 66.6% of all candidate regulatory targets of both LTR7 and LTR5_Hs loci (**Table 8**), which is significantly higher than corresponding metrics recorded for genes linked with either LTR7 loci (p = 1.53E-08; 2-tail Fisher’s exact test), LTR5_Hs loci (p = 1.70E-13; 2-tail Fisher’s exact test), or genes associated with at least two LTR7 loci (p = 9.958E-05; 2-tail Fisher’s exact test).

**Table 6.** A catalogue of human genes identified as regulatory targets of LTR5_Hs elements.

| **Classification category** | **Number of records** |
| --- | --- |
| **LTR5_Hs loci (hg38)** | 606 |
| **GREAT linked genes** | 935 |
| **CRISPRi target genes\*** | 4251 |
| **CRISPRi/GREAT target genes** | 447 |
| **KRABa target genes\*** | 621 |
| **KRABa/GREAT target genes** | 191 |
| **CRISPRi/KRABa/GREAT target genes** | **486** |

Legend: \*number of LTR5\_Hs target genes in the GREAT hg38 set of 18777 human genes.

**Table 7.**
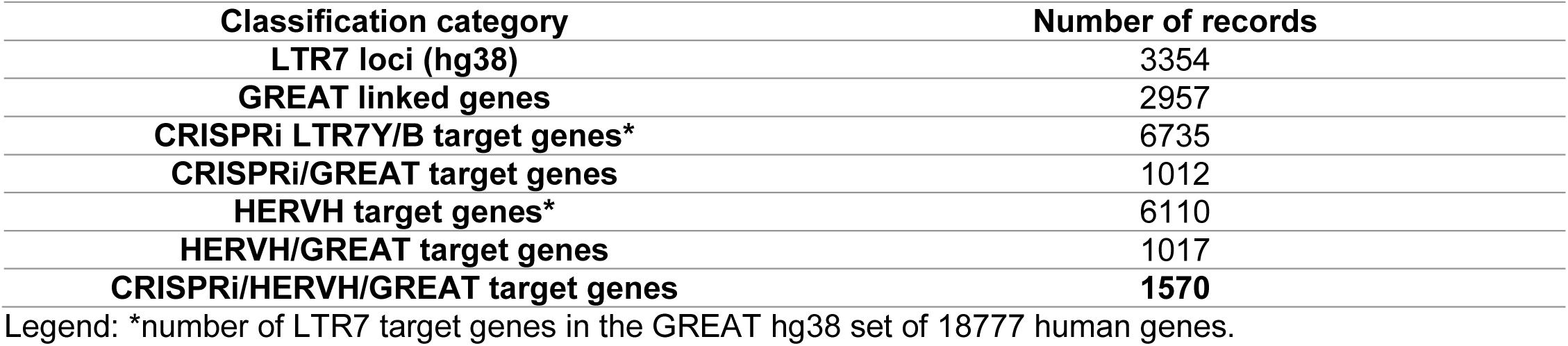
A catalogue of human genes identified as regulatory targets of LTR7 elements.

**Table 8.** A significant majority of LTR-linked mammalian OS genes and fetal gonad’s cells marker genes are high fidelity LTR-regulated genes.

| Classification category | Number of genes | LTR-regulated | Percent | P value* |
| --- | --- | --- | --- | --- |
| LTR7 target genes | 2957 | 1570 | 53.09 |  |
| LTR5_Hs target genes | 935 | 486 | 51.98 |  |
| LTR7 and/or LTR5_Hs target genes | 3515 | 1944 | 55.31 |  |
| LTR7 target genes (at least 2 LTR7 loci per gene) | 1202 | 664 | 55.24 | 9.958E-05 |
| LTR7 and LTR5_Hs target genes | 377 | 251 | **66.58** | 9.958E-05 |
| LTR7 only target genes | 2580 | 1319 | 51.12 | 1.53E-08 |
| LTR5_Hs only target genes | 558 | 235 | 42.11 | 1.70E-13 |
| LTR7 mammalian OS target genes | 562 | 392 | **69.75** | 5.96E-19 |
| LTR7 mammalian non-OS target genes | 2395 | 1178 | 49.19 | 5.96E-19 |
| LTR5_Hs mammalian OS target genes | 126 | 92 | **73.02** | 2.85E-07 |
| LTR5_Hs mammalian non-OS target genes | 809 | 394 | 48.70 | 2.85E-07 |
| LTR7 and/or LTR5_Hs mammalian OS target genes | 630 | 432 | **68.57** | 8.34E-14 |
| LTR7 and/or LTR5_Hs mammalian non-OS target genes | 2885 | 1512 | 52.41 | 8.34E-14 |
| LTR7 and LTR5_Hs mammalian OS target genes | 58 | 50 | **86.21** | 0.0004271 |
| LTR7 and LTR5_Hs mammalian non-OS target genes | 319 | 201 | 63.01 | 0.0004271 |
| Human-specific LTR7 and/or LTR5_Hs target genes | 391 | 215 | 54.99 |  |
| Fetal gonad's cells markers hsLTR7 and/or hsLTR5_Hs target genes | 88 | 64 | **72.73** | 0.0001478 |
| Human-specific LTR7 and/or LTR5_Hs target genes (excluding fetal gonad's cell marker genes) | 303 | 151 | 49.83 | 0.0001478 |

Legend: \*, P values were estimated using the 2-tail Fisher's exact test.

Fractions of genes representing high-fidelity down-stream regulatory targets of LTR7 and LTR5_Hs loci appear significantly higher among mammalian OS genes compared to non-OS genes (**Table 8**), reaching 86.2% (p = 0.0004; 2-tail Fisher’s exact test) for mammalian OS genes defined as targets of both LTR7 and LTR5_Hs loci (**Table 8**). Similarly, 64 of 88 (72.7%) genes encoding Fetal Gonad’s cells markers associated with human-specific LTR7 and/or LTR5_Hs loci could be defined as their high-fidelity down-stream regulatory targets (**Table 8**). Therefore, these functional categories and corresponding genes linked by present analyses to LTR7 and LTR5_Hs genomic regulatory networks should be considered as high-priority aims for experimental validation of bona fide transcriptional regulatory targets of LTR7 and LTR5_Hs loci.

A majority of genes identified in this contribution as putative regulatory targets of LTR7 and/or LTR5_Hs loci manifests experimentally validated significant gene expression changes in response to targeted genetic or epigenetic manipulations (interference; silencing; or activation) of candidate up-stream regulatory LTRs (**Tables 6 – 8**). Therefore, these genes could be defined as high-fidelity down-stream regulatory targets of LTR7 and/or LTR5_Hs loci. Taking into account these observations, it was of interest to evaluate potential biological impacts of LTR7 and LTR5_Hs loci on physiology and pathology of Modern Humans employing GSEA focused only on high-fidelity down-stream regulatory target genes of LTR7 and LTR5_Hs elements. Results of these analytical experiments reinforce and extend reported above observations on biological roles and plausible pathophysiological effects of LTR7- and LTR5_Hs-linked genes (**Figures 9** **- 12; Supplementary Figures S3, S4; Supplementary Table S4**).

**Figure 9.**
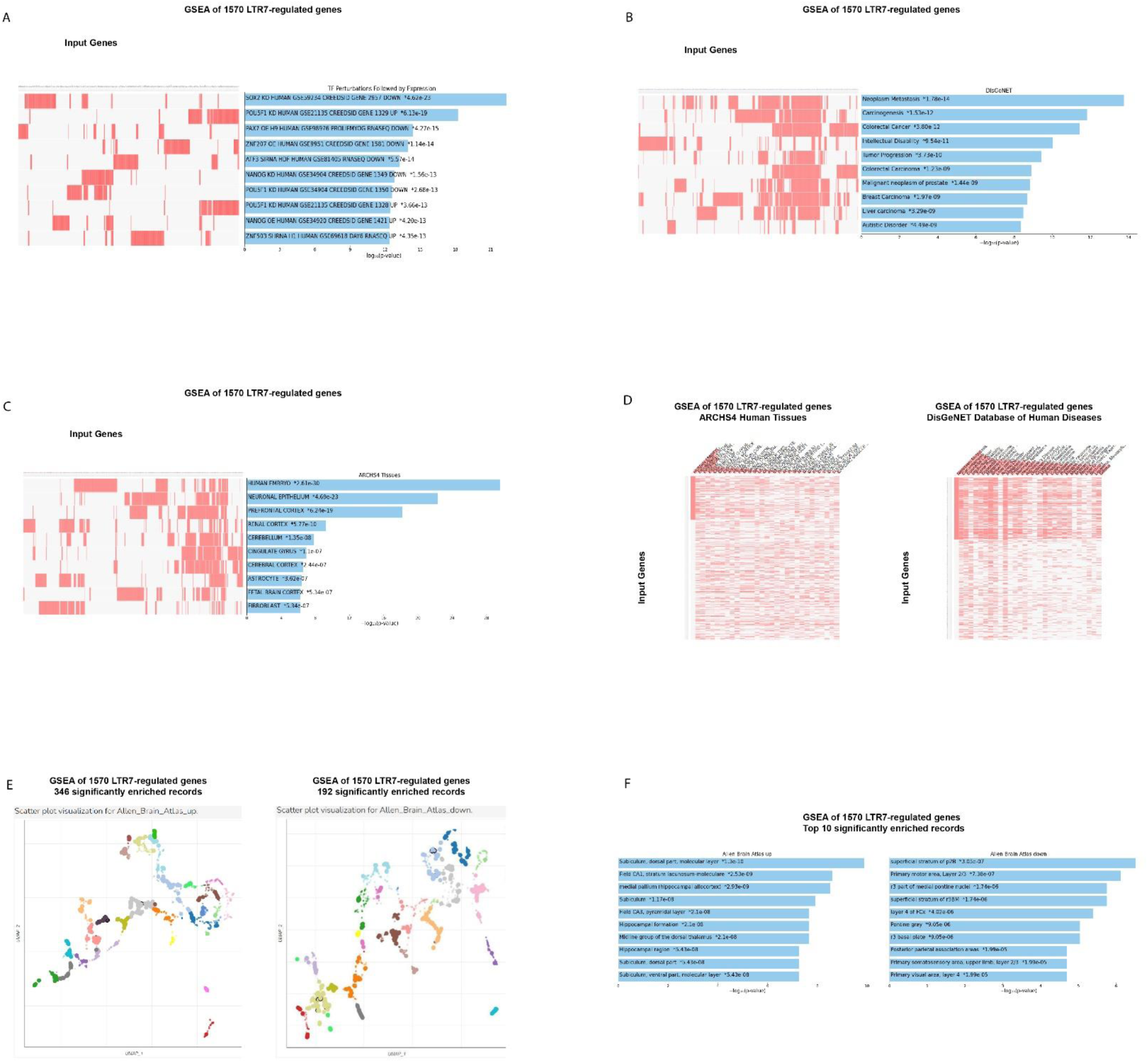
Potential phenotypic impacts of LTR7 elements revealed by GSEA of 1570 high-fidelity down-steam target genes employing the Transcription Factors (TF) perturbations followed by expression database (A), the DisGeNET database of human diseases (B; D), the ARCHS4 Human Tissues database (C; D), the Allan Brain Atlas databases of up-regulated genes (E; F; left panels) and down-regulated genes (E; F; right panels). In (D) top 30 significantly enriched records of gene sets were sorted by genes comprising the expression signature of the Human Embryo (left panel) and genes comprising the expression signature of the Neoplasm Metastasis (right panel). In panels (A), (B), (C), and (F) results illustrating the top 10 significantly enriched records are reported. All reported significantly enriched records were identified at the significance threshold of adjusted p value < 0.05 by the GSEA of 1570 LTR7-regulated genes employing the corresponding genomic databases (Methods).

Underscoring the important roles of LTR7 and LTR5_Hs loci in regulation of stemness and pluripotency state-related phenotypes, GSEA of 377 genes linked with both LTR7 and LTR5_Hs regulatory elements employing Human RNAseq Automatic GEO Signatures database identified amongst top 10 significantly enriched traits gene expression signatures of Naïve and Primed pluripotent states (p = 2.41E-09); chromatin-associated Sin3B protein-regulated quiescence (p = 1.27E-08); and pluripotent state 3D chromosome landscape (p = 6.40E-08) (**Supplementary Figure S3A**). Recapitulating, in part, phenotype-enrichment findings attributed to human-specific regulatory LTR elements (**Figure 5**), GSEA of the DisGeNET database of human disorders identified Non-obstructive Y-linked Spermatogenic Failure (p = 1.32E-08); Male sterility due to Y-chromosome deletions (p = 1.45E-05); Partial chromosome Y deletion (p = 1.45E-05); Oligospermia (p = 4.09E-05); and Chromic Alcoholic Intoxication (p = 6.07E-05) amongst top significantly enriched records (**Supplementary Figures S3B; 3C**). Schizophrenia (p = 2.40E-05) and Autism Spectrum Disorder (p = 3.55E-05) were scored as top 2 significantly enriched traits based on GSEA of the Disease Perturbations from GEO database focused on down-regulated genes (**Supplementary Figure S3D**).

These observations indicate that despite distinct evolutionary histories separated by millions’ years of primates’ germlines colonization and expansion, genes representing down-stream regulatory targets of LTR7 and LTR5_Hs loci seem to exert the apparently cooperative phenotypic effects ascertained from the significantly enriched traits recorded by GSEA.

The apparent cooperative effects on phenotypic traits could be seeing when significantly enriched records attributed to 1570 high-fidelity target genes of LTR7 loci (**Figure 9**) and 486 high-fidelity target genes of LTR5_Hs (**Figure 10**) were compared for GSEA of the Transcription Factors’ (TF) perturbations followed by expression analyses database. These observations indicate that master transcriptional regulators of the pluripotency phenotype, namely SOX2, POU5F1, and NANOG, represent common up-stream TFs regulating expression of high-fidelity regulatory target genes of both LTR7 and LTR5_Hs loci. This conclusion is supported by results of GSEA of 1944 genes comprising high-fidelity down-steam regulatory targets of LTR7 and/or LTR5_Hs elements (**Figure 11**) as evidenced by higher numbers of implicated signature genes and lower enrichment p values for down-stream targets of the SOX2 and POU5F1 TFs recorded by GSEA of a cumulative set of 1944 down-stream target genes. Increased numbers of enriched signature genes and lower enrichment p values for a cumulative set of 1944 genes consisting of high-fidelity down-stream regulatory targets of LTR7 and/or LTR5_Hs elements (**Figures 11; 12**) compared to gene sets linked with either LTR7 (**Figure 9**) or LTR5_Hs (**Figure 10**) loci were recorded for numerous significantly enriched phenotypic traits, including human embryo development stages, morphological components of the central nervous system (prefrontal cortex, cerebellum, fetal brain cortex, subiculum, hippocampal formation), and different cell types of fetal gonads. Exceptions from this trend were noted for GSEA of the Human RNAseq Automatic signatures database (**Figure 10C** and **Figure 11A**), indicating that down-stream target genes comprising expression signatures related to Naïve and Primed pluripotent states appear biased toward LTR5_Hs regulatory elements.

**Figure 10.**
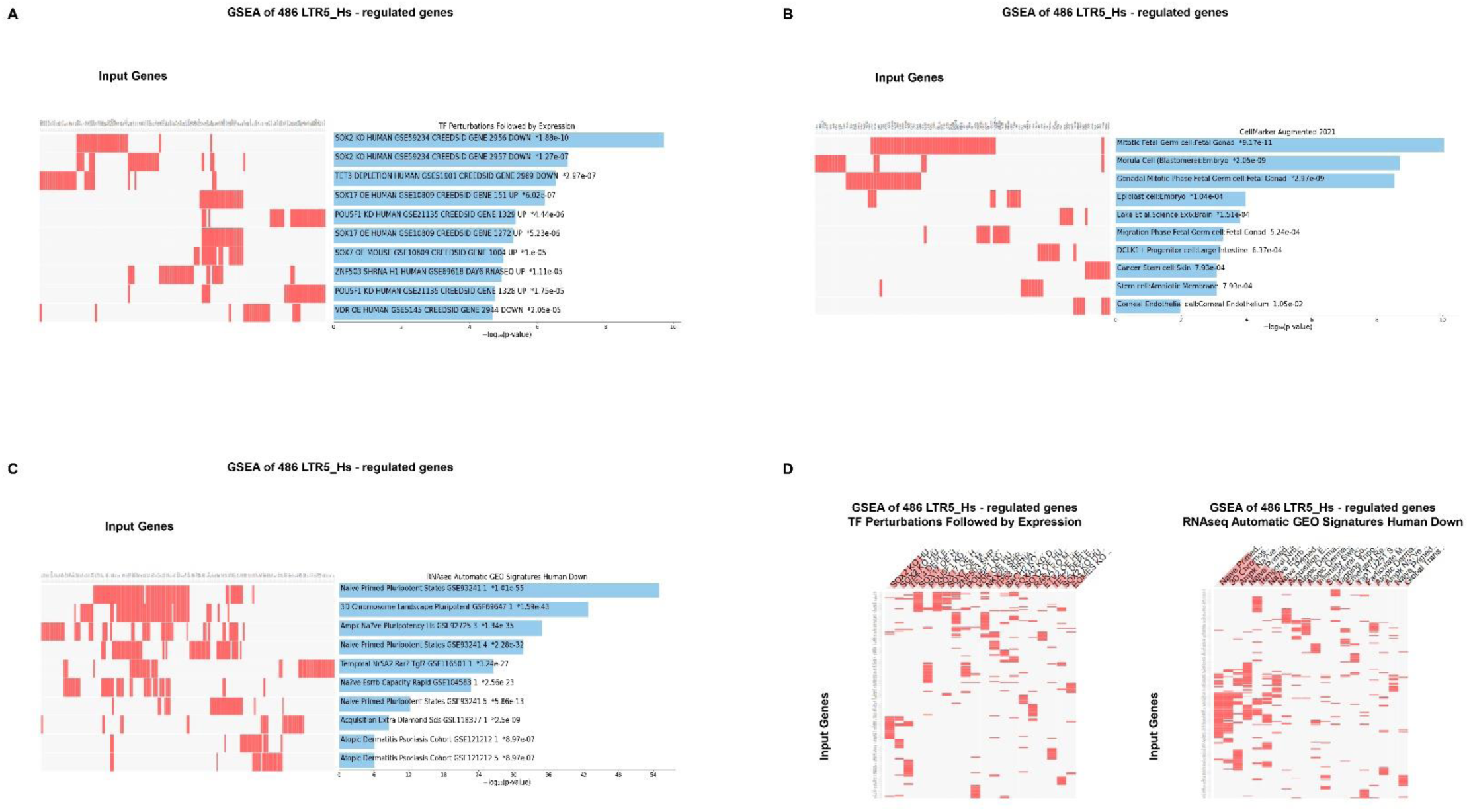
Assessments of potential phenotypic impacts of LTR5_Hs elements based on GSEA of 486 high-fidelity down-steam target genes employing the Transcription Factors (TF) perturbations followed by expression database (A; D, left panel), the Cell Marker Augmented 2021 database (B), the Human RNAseq Automatic GEO Signatures database (C; D, right panel). In panels (A), (B), and (C) results illustrating the top 10 significantly enriched records are reported. In (D) top 30 significantly enriched records of gene sets are reported. All reported significantly enriched records were identified at the significance threshold of adjusted p value < 0.05 by the GSEA of 486 LTR5_Hs-regulated genes employing the corresponding genomic databases (Methods).

**Figure 11.**
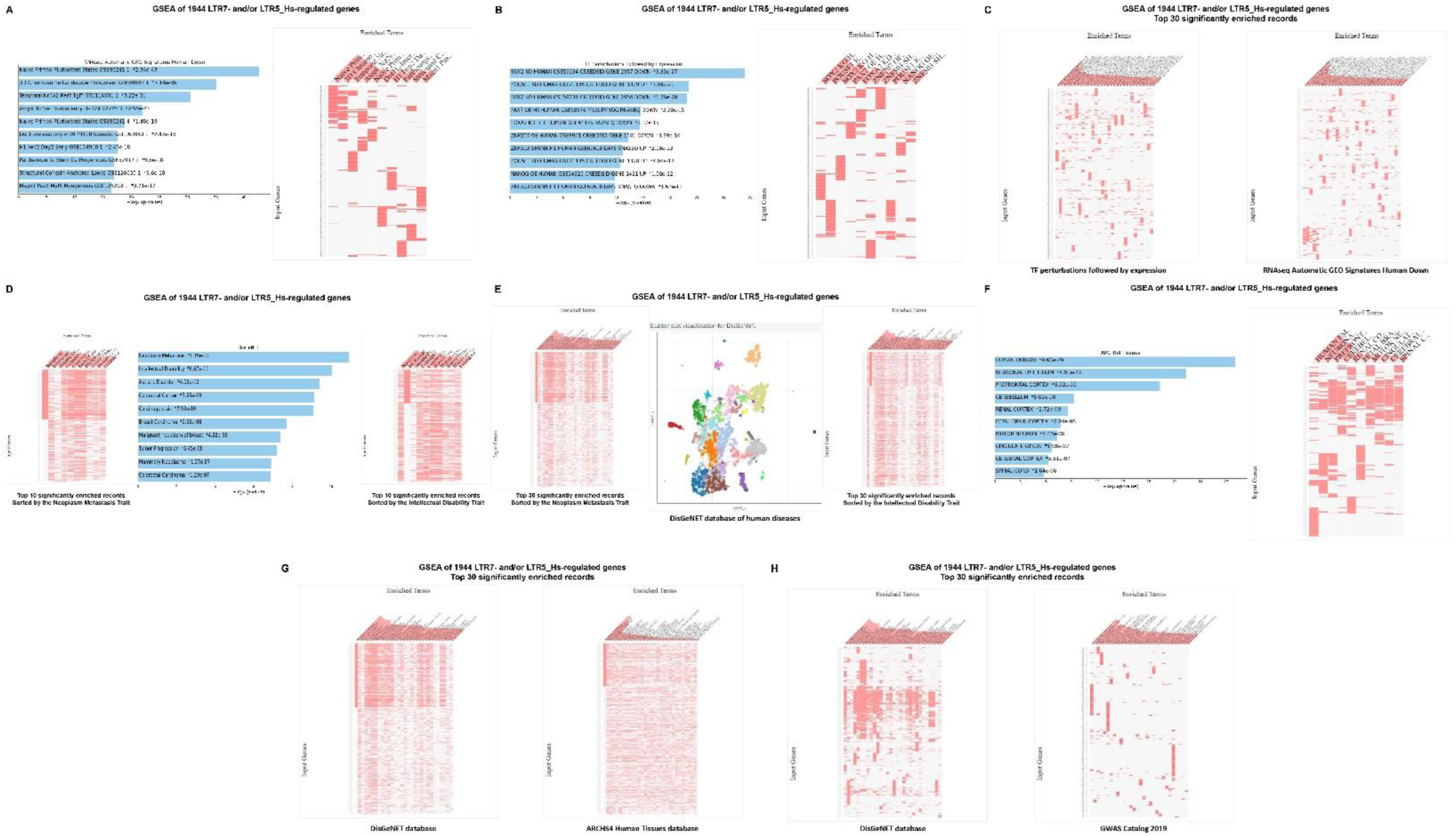
Potential phenotypic impacts of LTR7 and LTR5_Hs elements revealed by GSEA of 1944 high-fidelity down-steam target genes employing the Human RNAseq Automatic GEO Signatures database (A; C, right panel), the Transcription Factors (TF) perturbations followed by expression database (B; C, left panel), the DisGeNET database of human diseases (D; E; G and H, left panels), the ARCHS4 Human Tissues database (F; G, right panels), and the GWAS Catalog 2019 database (H, right panel). In (D), top 10 significantly enriched records of gene sets were sorted by genes comprising the expression signature of the Intellectual Disability trait (right panel) and genes comprising the expression signature of the Neoplasm Metastasis trait (left panel). In (E), top 30 significantly enriched records of gene sets were sorted by genes comprising the expression signature of the Intellectual Disability trait (right panel) and genes comprising the expression signature of the Neoplasm Metastasis trait (left panel). Middle panel in (E) shows scatterplot visualization of GSEA results of the DusGeNET database of human diseases. In panels (A), (B), (D), and (F) results illustrating the top 10 significantly enriched records are reported. In panels (C), (E), (G), and (H) results illustrating the top 30 significantly enriched records are reported. All reported significantly enriched records were identified at the significance threshold of adjusted p value < 0.05 by the GSEA of 1944 genes regulated by LTR7- and/or LTR5_Hs loci employing the corresponding genomic databases (Methods).

**Figure 12.**
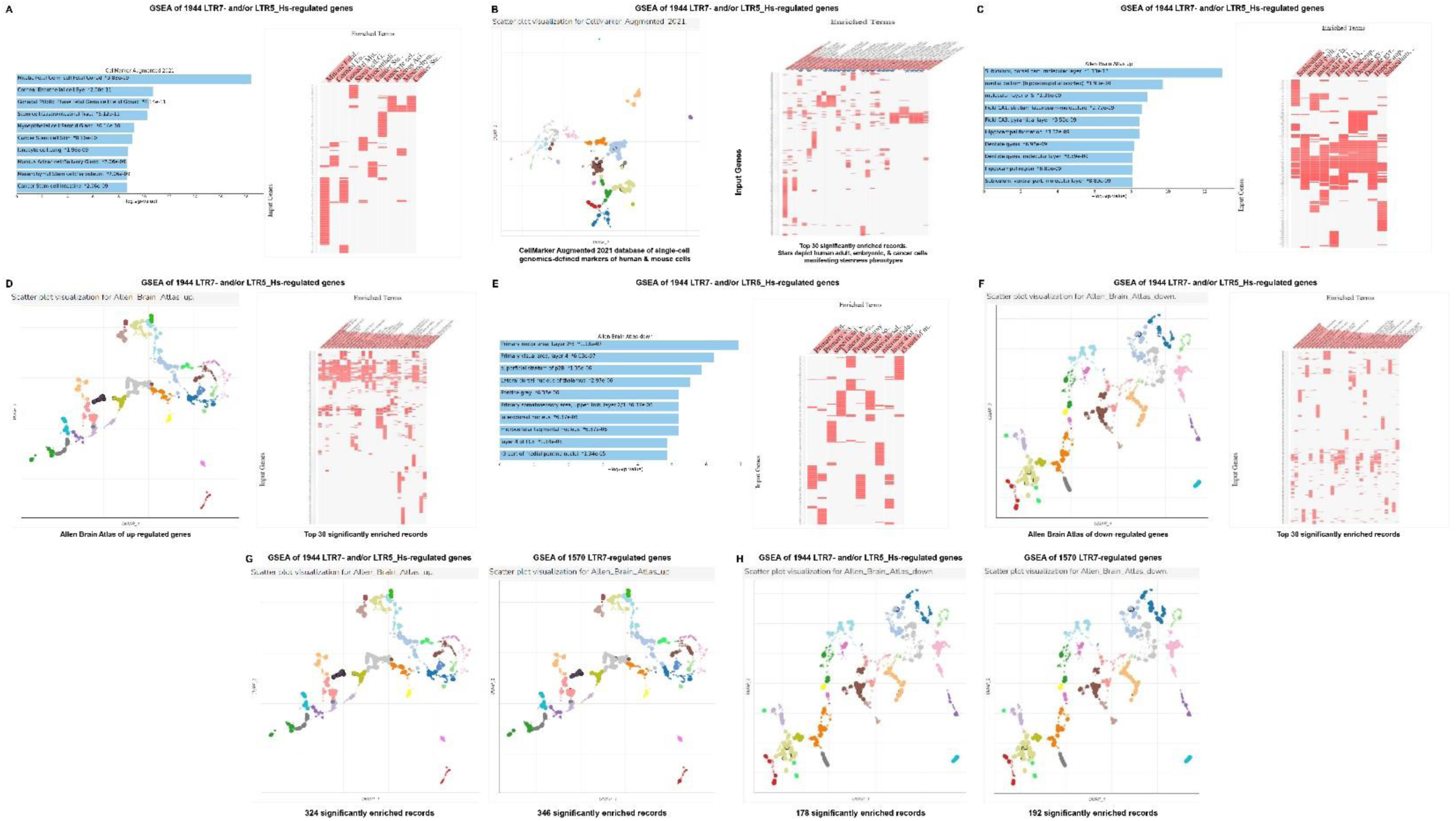
Potential phenotypic impacts of LTR7 and LTR5_Hs elements revealed by GSEA of 1944 high-fidelity down-steam target genes employing the Cell Marker Augmented 2021 database (A; B), the Allan Brain Atlas databases of up-regulated genes (C; D; G) and down-regulated genes (E; F; H). Left panel in (B) shows scatterplot visualization of GSEA results of the Cell Marker Augmented 2021 database of single-cell genomics-defined markers of human cells. Right panel in (B) reports top 30 significantly enriched records of gene sets identified by the GSEA of the Cell Marker Augmented 2021 database highlighting by stars gene expression signatures of human adult, embryonic, and cancer cells manifesting stemness phenotypes. Left panel in (D) shows scatterplot visualization of GSEA results of the Allan Brain Atlas database of up-regulated genes. Right panel in (D) reports top 30 significantly enriched records of gene sets identified by the Allan Brain Atlas database of up-regulated genes. Left panel in (F) shows scatterplot visualization of GSEA results of the Allan Brain Atlas database of down-regulated genes. Right panel in (F) reports top 30 significantly enriched records of gene sets identified by the Allan Brain Atlas database of down-regulated genes. Figure (G) shows side-by-side aligned scatterplots’ visualization of GSEA results of the 1944 LTR7- and/or LTR5_Hs-regulated genes (left panel; 324 significantly enriched records) and 1570 LTR7-regulated genes (right panel; 346 significantly enriched records) employing the Allan Brain Atlas database of up-regulated genes. Figure (H) shows side-by-side aligned scatterplots’ visualization of GSEA results of the 1944 LTR7- and/or LTR5_Hs-regulated genes (left panel; 178 significantly enriched records) and 1570 LTR7-regulated genes (right panel; 192 significantly enriched records) employing the Allan Brain Atlas database of down-regulated genes. In panels (A), (C), and (E) results illustrating the top 10 significantly enriched records are reported. All reported significantly enriched records were identified at the significance threshold of adjusted p value < 0.05 by the GSEA of genes regulated by LTR7- and/or LTR5_Hs loci employing the corresponding genomic databases (Methods).

In several instances GSEA of a cumulative set of 1944 genes consisting of high-fidelity down-stream regulatory targets of LTR7 and/or LTR5_Hs elements (**Figures 11; 12**) recorded the apparent qualitative differences compared to gene sets linked with either LTR7 (**Figure 9**) or LTR5_Hs (**Figure 10**) loci. For example, GSEA of the Cell Marker Augmented 2021 database revealed that human cells manifesting stemness phenotypes, including tissue-specific adult stem cells, embryonic stem cells, and cancer stem cells residing in various organs, consist a large majority (67%) of top 30 significantly enriched records (**Figure 12B**).

Similarly, increased numbers of up-regulated genes of dentate gyrus signatures and lower enrichment p values were recorded by GSEA of the Allen Brain Atlas database for a cumulative set of 1944 genes consisting of high-fidelity down-stream regulatory targets of LTR7 and/or LTR5_Hs elements (**Figure 12C**). However, the overall enrichment patterns of LTR high-fidelity target genes identified by GSEA of the Allen Brain Atlas databases for both up-regulated (**Figure 12G**) and down-regulated (**Figure 12H**) genes appear driven by genes expression of which is regulated by LTR7 elements.

GSEA of 1944 genes comprising a cumulative set of high-fidelity down-steam regulatory targets of LTR7 and/or LTR5_Hs loci revealed marked enrichment for genes implicated in neoplasm metastasis, intellectual disability, multiple cancer types, autism, Alzheimer’s, schizophrenia, and other brain disorders (**Figures 11, 12**). Similar to findings recorded for 562 LTR7-linked mammalian OS genes (**Figure 6**), three most significantly enriched records of human diseases (Neoplasm metastasis; p = 1.15E-11; Intellectual disability; p = 6.67E-11; and Autistic disorder; p = 4.06E-10) appear associated with partially overlapping networks of genes (**Figure 11E; 11D**), suggesting that transcriptional misregulation of high-fidelity down-stream target genes of LTR7 and/or LTR5_Hs elements may contribute to pathogenesis of multiple types of human cancers and brain disorders.

## Discussion

### Insights from LTR family-specific and granular analyses of evolutionary origin, expansion, conservation, and divergence of LTR7 and LTR5_Hs elements during primate evolution

In this contribution, targeted DNA sequences conservation analyses of 17 primate species were employed to identify genomes manifesting the consistent presence of highly-conserved (HC) retroviral LTR sequences at the earliest time points during primate evolution. Results of these analyses were utilized to infer the timelines of initial colonization and subsequent expansion of human endogenous retroviruses (HERV) LTR7/HERVH and LTR5_Hs/HERVK amongst primate species. LTR7/HERVH and LTR5_Hs/HERVK retroviruses appear to have distinct evolutionary histories of successful colonization of primates’ genomes charted by evidence of the earliest consistent presence and expansion of HC LTR sequences. In contrast to genomes of New World Monkeys, genomes of the Old World Monkey lineage consistently harbor ∼18% of HC-LTR7 loci residing in genomes of Modern Humans. These observations suggest that LTR7/HERVH have entered germlines of primate species in Africa after the separation of the New World Monkey lineage. The earliest presence of HC-LTR5_Hs loci has been identified in the Gibbon’s genome (24% of HC-LTR5_Hs loci residing in human genomes), suggesting that LTR5_Hs/HERVK successfully colonized primates’ germlines after the segregation of Gibbons’ species. Subsequently, both LTR7 and LTR5_Hs underwent a marked ∼4-5-fold expansion in genomes of Great Apes. Intriguingly, timelines of quantitative expansion of both LTR7 and LTR5_Hs loci during evolution of Great Apes appear to replicate the consensus evolutionary sequence of increasing cognitive and behavioral complexities of non-human primates, which seems particularly striking for LTR7 loci and 8 distinct LTR7 subfamilies.

Discovery of a complex polyphyletic composition of LTR7 elements comprising of at least eight monophyletic subfamilies (Carter et al., 2022) prompted sequence conservation analyses of each individual monophyletic subfamilies of LTR7 loci in genomes of sixteen NHP. Results of these analyses suggest that diversification of LTR7 loci into genetically and regulatory distinct subfamilies may have occurred early during primate evolution and subsequent cycles of LTR7 expansion appear to faithfully maintain this diversity. This conclusion is supported by observations that highly conserved sequences of all 8 monophyletic LTR7 subfamilies are present in genomes of all analyzed in this study Old World Monkeys’ species as well as in genomes of Gibbon, Orangutan, Gorilla, Bonobo, and Chimpanzee (**Figure 2**). Despite large differences in the numbers of highly conserved LTR7 loci amongst different primate species, the overall quantitative balance of 8 distinct monophyletic LTR7 subfamilies appears maintained very tightly during millions’ years of primate evolution, which is reflected by nearly perfect correlations of LTR7 subfamilies abundance profiles between evolutionary closely-related primate species (**Figure 3**). On a 40 MYA scale of primate evolution from Old World Monkeys to Modern Humans, the phenomenon of conservation of LTR7 subfamilies abundance profiles is illustrated by the strong inverse correlation (r = - 0.986) of a degree of resemblance of NHP species’ and Modern Humans LTR7 subfamilies abundance profiles and estimated divergence time from ECA (**Figure 3C**).

One of the notable features of LTR7 sequences conservations analyses in genomes of five Old World Monkeys’ species was strikingly similar numbers of gains and losses of LTR7 loci independently estimated for each species (**Figure 1**). Using these data as a baseline for estimates of numbers of LTR7 loci gains per MYA during primate evolution (**Figure 1D**), a model of a putative species-specific expansion of regulatory LTR7 loci during primate evolution was built for genomes of eleven NHP (**Figure 1C**). For each species, relative gains (a number of highly conserved LTR7 loci identified in a genome) and losses (a deficit of highly conserved LT7 loci compared to genomes of Modern Humans) of LTR7 loci were calculated vis-a-vis Modern Human’s genome housing 3354 highly-conserved LTR7 elements (**Figure 1C**). Based on these estimates tailored to a presumed timeline of Old World Monkeys’ segregation from ECA, a hypothetical model defining primate species segregation timelines from ECA was developed, which implies putative direct associations of LTR7 loci acquisitions in primates’ genomes with timelines of species segregation processes during primate evolution (**Figure 1D**). It will be of interest to determine whether this apparent association might reflect the causative impacts of LTR7/HERVH expansions on the emergence and segregation of primate species.

### Insights from analyses of potential phenotypic impacts of LTR7 and LTR5_Hs elements on physiology and pathology of Modern Humans

Extensive investigations of LTR7 elements conclusively documented their regulatory functions and locus-specific differential expression in human preimplantation embryogenesis as well as in human embryonic and pluripotent stem cells (Fort et al., 2014; Gemmell et al., 2015; Glinsky et al., 2018; Göke et al.,2015; Izsvák et al., 2016; Kelley and Rinn, 2012; Loewer et al., 2010; Lu et al., 2014; Ohnuki et al., 2014; Pontis et al., 2019; Römer et al., 2017; Santoni et al., 2012; Takahashi et al., 2021 Theunissen et al., 2016; Wang et al., 2014; Zhang et al., 2019). Similarly, LTR5_Hs/HERVK-derived loci manifest transcriptional and biological activities in human preimplantation embryos and in naïve hESCs (Grow et al., 2015). Many facets of activities of LTR5_Hs elements were attributed to acquisition of enhancer-like chromatin state signatures concomitantly with transcriptional reactivation of HERVK sequences (Grow et al., 2015), consistent with LTR5_Hs elements acting as distal enhancers exerting global long-range effects on expression of thousands human genes (Fuentes et al., 2018). Thus, our understanding of LTR7 and LTR5_Hs functions was restricted to a large degree to preimplantation embryogenesis, hESC, and pluripotent stem cells. Results of analytical experiments carried out in this contribution strongly argue that LTR7 and LTR5_Hs elements may affect many previously underappreciated aspects of physiological functions and pathological conditions of Modern Humans.

Inferences of potential phenotypic effects of LTR7 and LTR5_Hs elements were based on assessments of experimentally validated biological functions and cell type-specific differential expression profiles of genes identified as down-stream regulatory targets of LTR7 and LTR5_Hs loci. To ensure the high-stringency definition of candidate down-stream regulatory targets and reduce the likelihood of spurious associations, only significantly enriched records of gene signatures and linked phenotypic traits identified by GSEA at the significance threshold of adjusted p value < 0.05 and/or FDR q value < 0.05 were considered. Importantly, all observations that have been considered as strong evidence of implied biological effects of LTR7 and LTR5_Hs loci were validated by GSEA of genes experimentally defined as down-stream regulatory targets of LTR7 and LTR5_Hs elements. Confirming the validity of this analytical approach, the important roles of LTR7 and LTR5_Hs loci in regulation of preimplantation embryogenesis, stemness, and pluripotency state-related phenotypes have been documented for both LTR7 and LTR5_Hs regulatory elements.

One of the most intriguing findings reported herein is the postulated regulatory effect of human-specific LTR7 and LTR5_Hs loci on genes encoding markers of 12 distinct cells’ populations of fetal gonads, as well as genes implicated in physiology and pathology of human spermatogenesis, including Y-linked spermatogenic failure, oligo- and azoospermia. Identified in this contribution readily available well-characterized mouse models conclusively linking genes and phenotypes of interest should facilitate the experimental testing of the validity of this hypothesis.

Mammalian offspring survival (MOS) genes have been identified as one of consistent regulatory targets throughout ∼30 MYA of the divergent evolution of LTR7 loci. Significantly, differential GSEA of LTR-linked MOS versus non-MOS genes identified dominant enrichment patterns of phenotypic traits affected by 562 LTR7-regulated and 126 LTR5_Hs-regulated MOS genes. Specifically, GSEA of LTR7-linked MOS genes identified more than 2200 significantly enriched records of human common and rare diseases and 466 significantly enriched records of Human Phenotype Ontology traits, including 92 genes of Autosomal Dominant Inheritance and 93 genes of Autosomal Recessive Inheritance. It will be of interest to test experimentally whether regulatory effects on MOS genes could be one of contributing genetic determinants driving species fitness and divergence during primate evolution.

One of the most consistent observations documented by interrogations of LTR7-linked down-stream target genes was a clear prevalence of enrichment records related to brain and CNS functions amongst significantly enriched phenotypic traits identified by GSEA of genomic databases focused on gene expression signatures of tissues and cell types across human body. For instance, GSEA of the single-cell sequencing PanglaoDB Augmented 2021 database identified significantly enriched records of gene signatures representative of cells of distinct neurodevelopmental stages and morphologically diverse cell types residing and functioning in human brain, including Neural Stem/Precursor cells, Radial Glia cells, Bergman Glia cells, Pyramidal cells, Tanycytes, Immature neurons, Interneurons, Trigeminal neurons, GABAergic neurons, and Glutamatergic neurons. GSEA of LTR7-linked downstream target genes employing the Allen Brain Atlas database identified 521 significantly enriched records of different human brain regions harboring expression signatures of both up-regulated (420 brain regions) and down-regulated (101 brain regions) genes. These observations indicate that LTR7-linked down-stream target genes may contribute to multiple facets of development and functions of human brain. In depth analyses of LTR7 and LTR5_Hs loci linked with down-stream target genes affecting synaptic transmission and protein-protein interactions at synapses provide further evidence supporting this hypothesis.

One of important conclusions that could be derived from present analyses is that despite clearly distinct evolutionary histories of LTR7/HERVH and LTR5_Hs/HERVK retroviruses separated in time by millions of years, genes representing down-stream regulatory targets of LTR7 and LTR5_Hs loci exert the apparently cooperative and exceedingly broad phenotypic impacts on physiology and pathology of Modern Humans. Considering random patterns of retroviral insertions during the initial stages of genome colonization and expansion, it would be of interest to determine how this cooperative phenotypic impacts have been attained and what the role of natural selection is in the alignment of phenotypic effects of distinct retroviral families. Collectively, observations reported in this contribution highlight LTR7 and LTR5_Hs regulatory elements as important genomic determinants of Modern Humans’ health and disease states, which exert their phenotypic impacts via effects on down-stream target genes from preimplantation embryogenesis throughout developmental stages to adulthood and aging.

## Supplementary Information is available online

Supplementary information includes Supplementary Tables S1 – S4 and Supplementary Figures S1 – S4.

## Author Contributions

This is a single author contribution. All elements of this work, including the conception of ideas, formulation, and development of concepts, execution of experiments, analysis of data, and writing of the paper, were performed by the author.

## Acknowledgements

This work was made possible by the open public access policies of major grant funding agencies and international genomic databases and the willingness of many investigators worldwide to share their primary research data. Author would like to thank you Victoria Glinskii for invaluable expert assistance with graphical presentation of the results of this study. This work was supported, in part, by OncoScar, LLC.

