## Supplementary for "Molecular diversity and phenotypic pleiotropy of genomic regulatory loci derived from human endogenous retrovirus type H (HERVH) promoter LTR7 and HERVK promoter LTR5_Hs and their impacts on pathophysiology of Modern Humans": Supplementary Figure S1,.pptx

### Slide 1
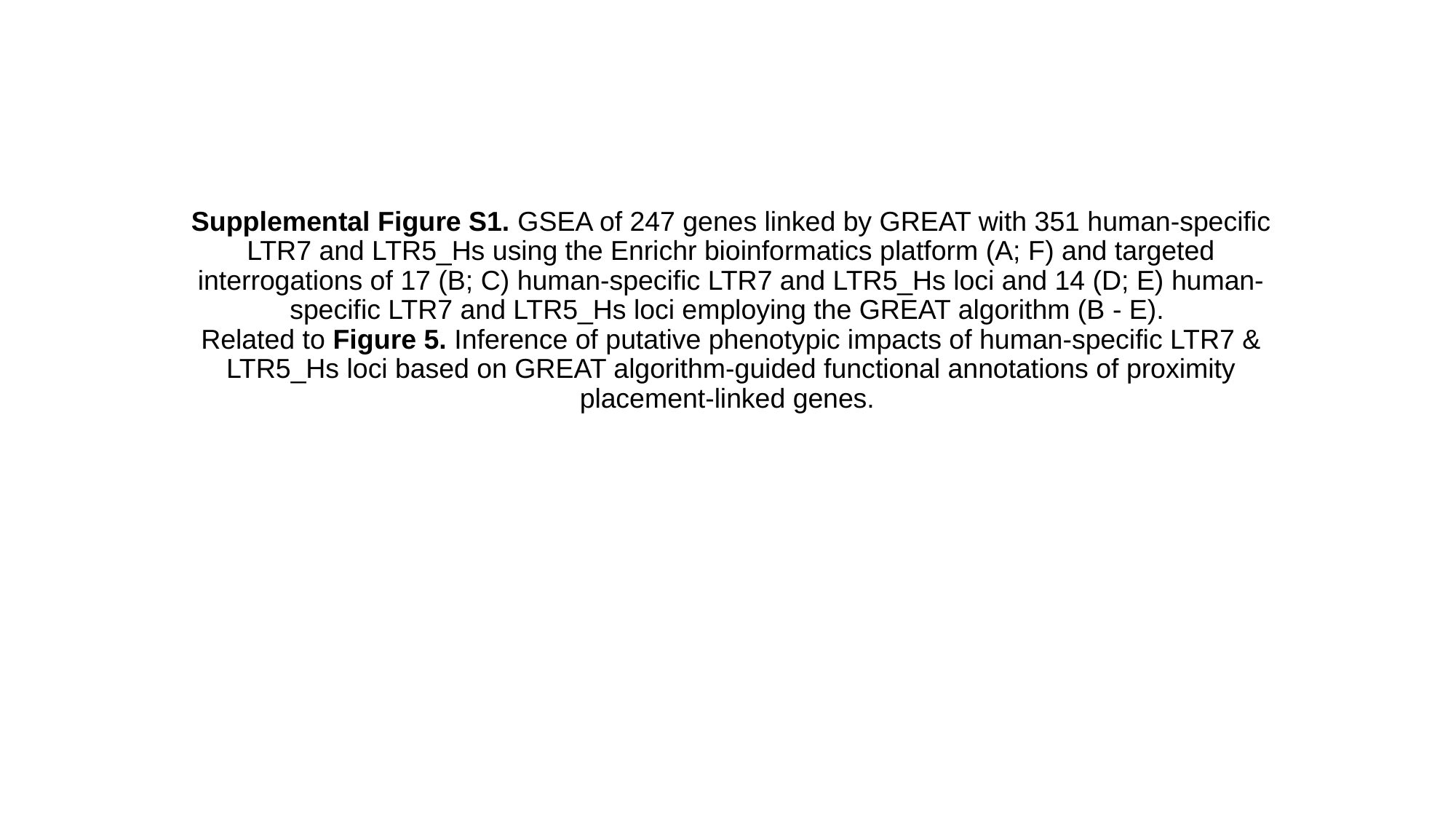

#### Supplemental Figure S1. GSEA of 247 genes linked by GREAT with 351 human-specific LTR7 and LTR5_Hs using the Enrichr bioinformatics platform (A; F) and targeted interrogations of 17 (B; C) human-specific LTR7 and LTR5_Hs loci and 14 (D; E) human-specific LTR7 and LTR5_Hs loci employing the GREAT algorithm (B - E). Related to Figure 5. Inference of putative phenotypic impacts of human-specific LTR7 & LTR5_Hs loci based on GREAT algorithm-guided functional annotations of proximity placement-linked genes.

### Slide 2
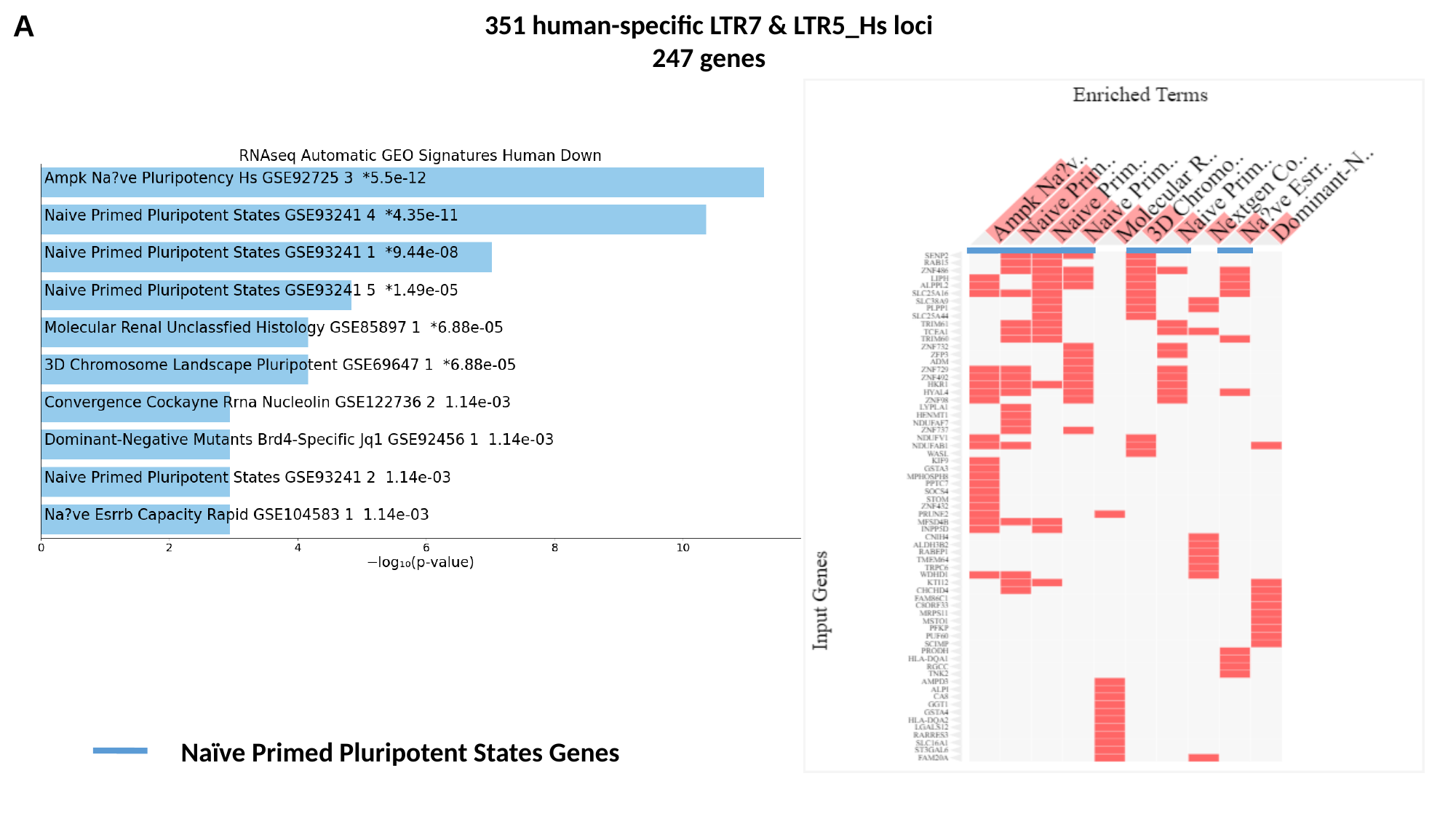

A
351 human-specific LTR7 & LTR5_Hs loci
247 genes
Naïve Primed Pluripotent States Genes

### Slide 3
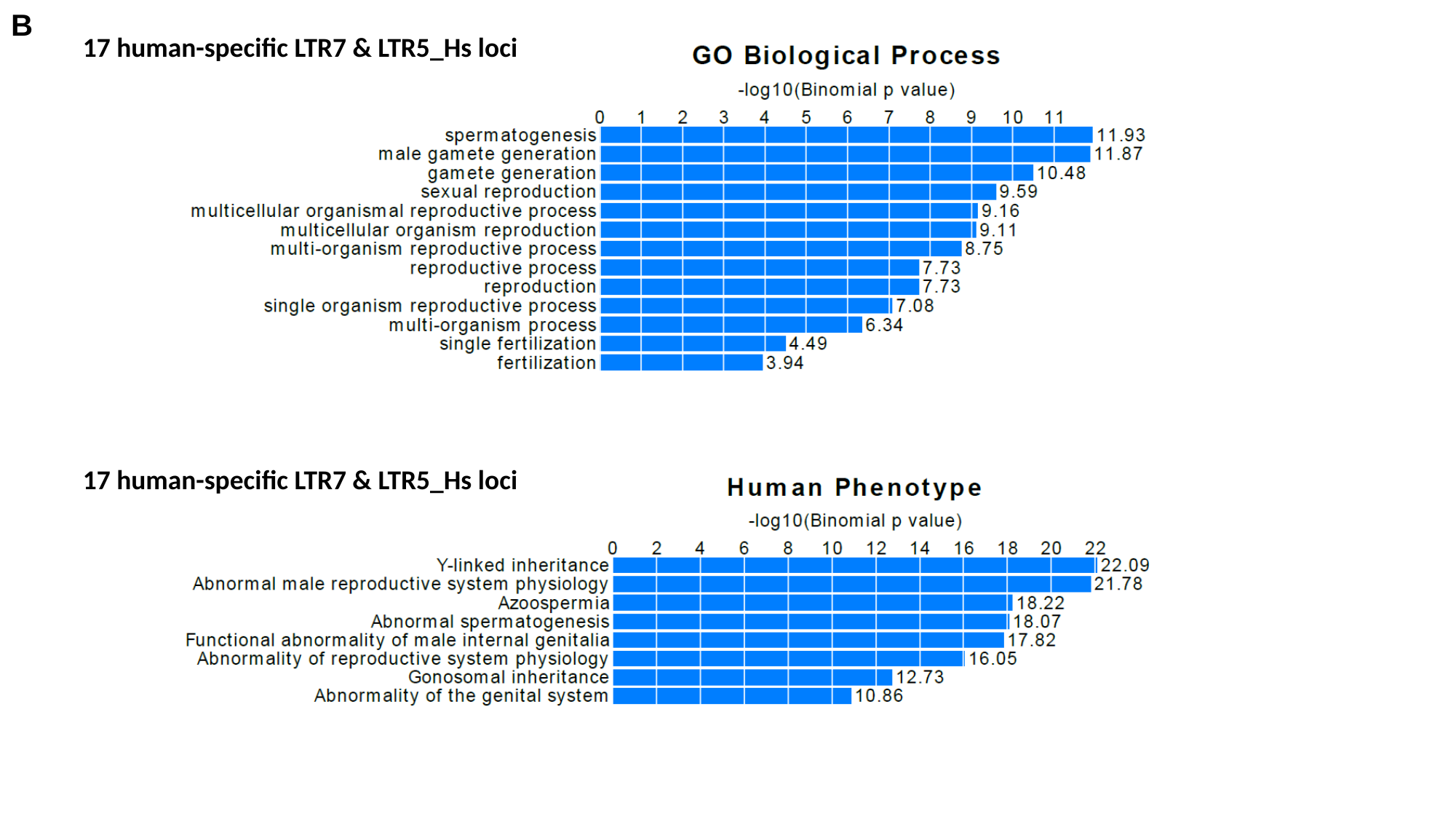

B
17 human-specific LTR7 & LTR5_Hs loci
17 human-specific LTR7 & LTR5_Hs loci

### Slide 4
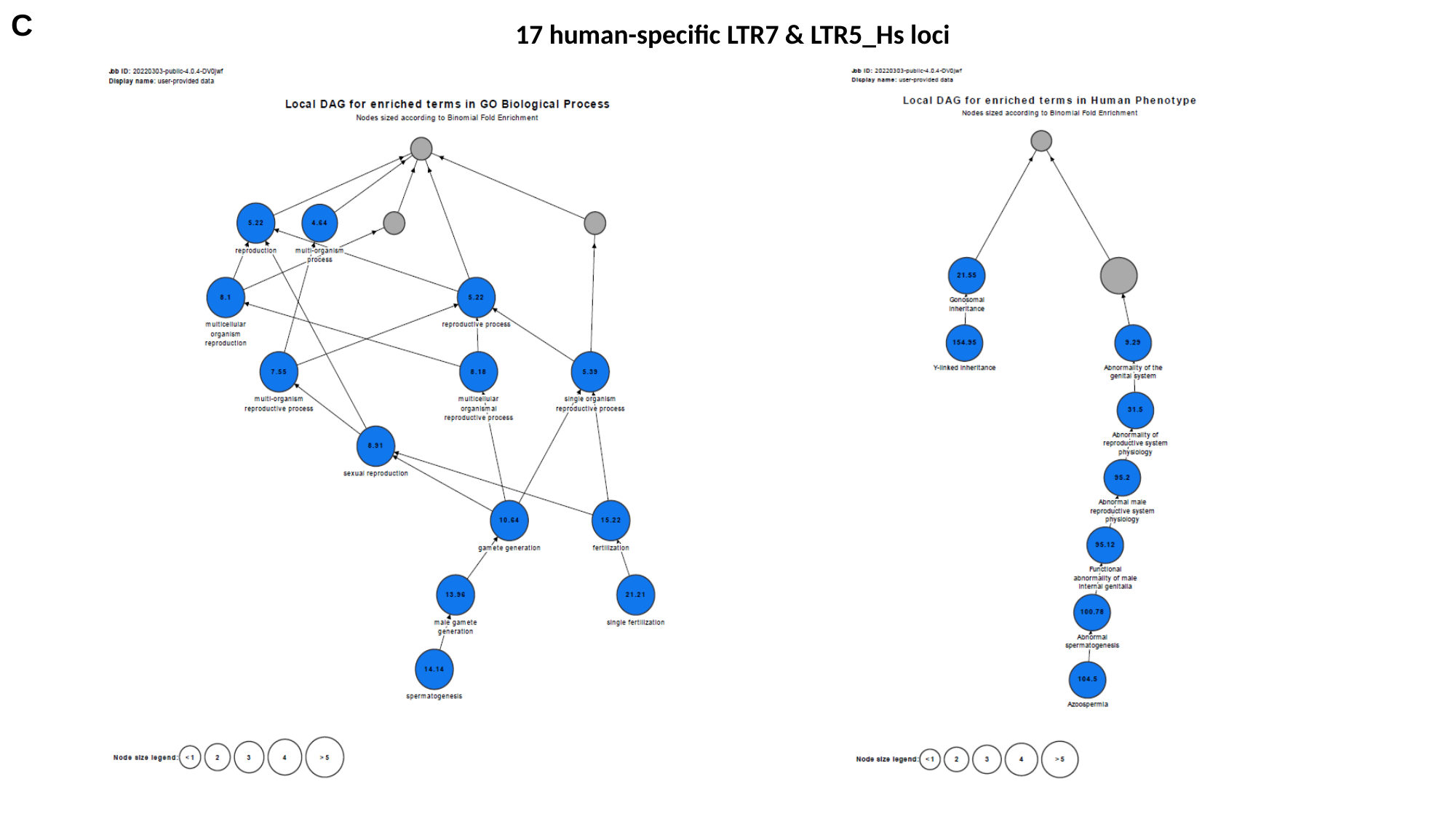

C
17 human-specific LTR7 & LTR5_Hs loci

### Slide 5
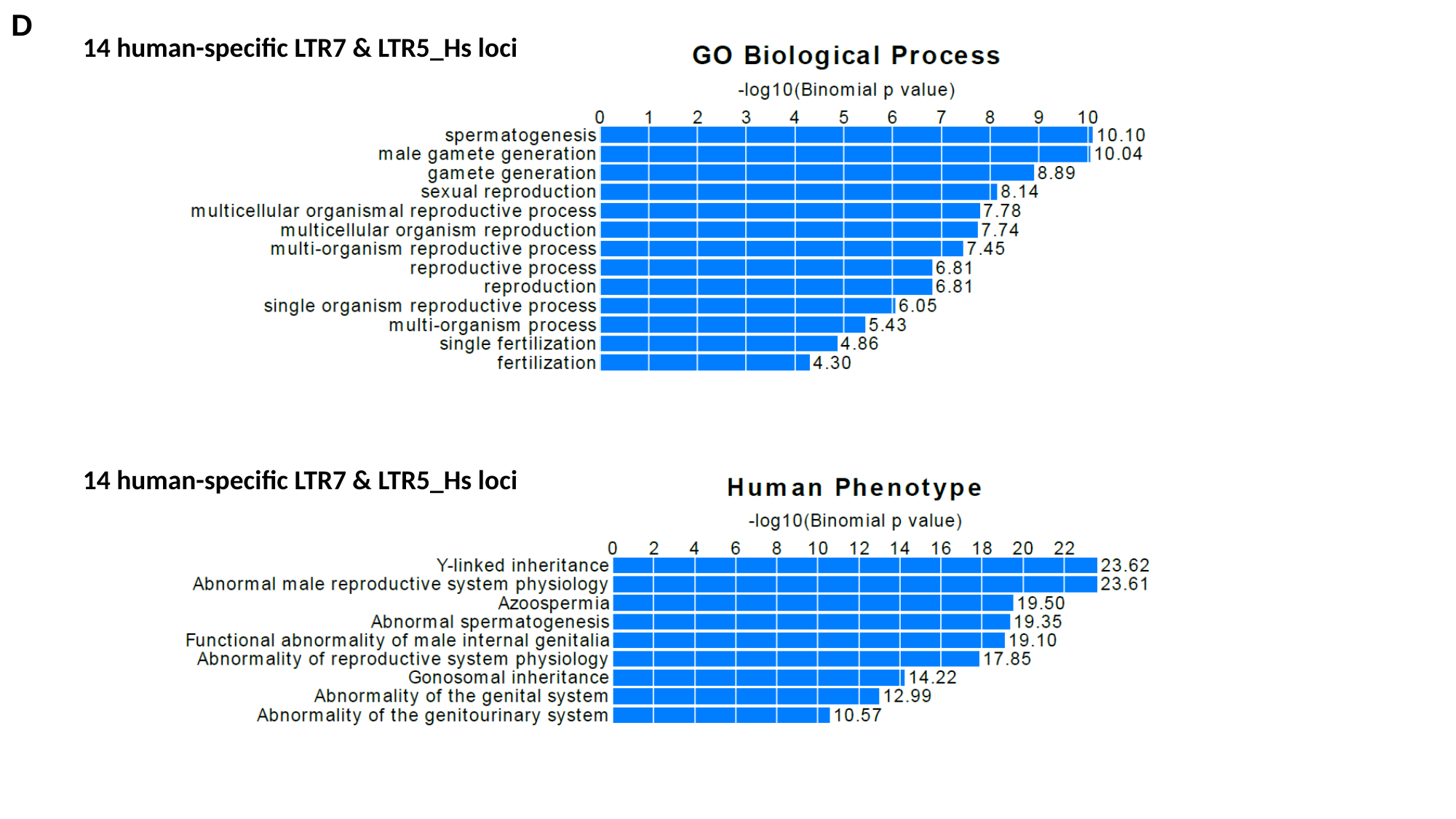

D
14 human-specific LTR7 & LTR5_Hs loci
14 human-specific LTR7 & LTR5_Hs loci

### Slide 6
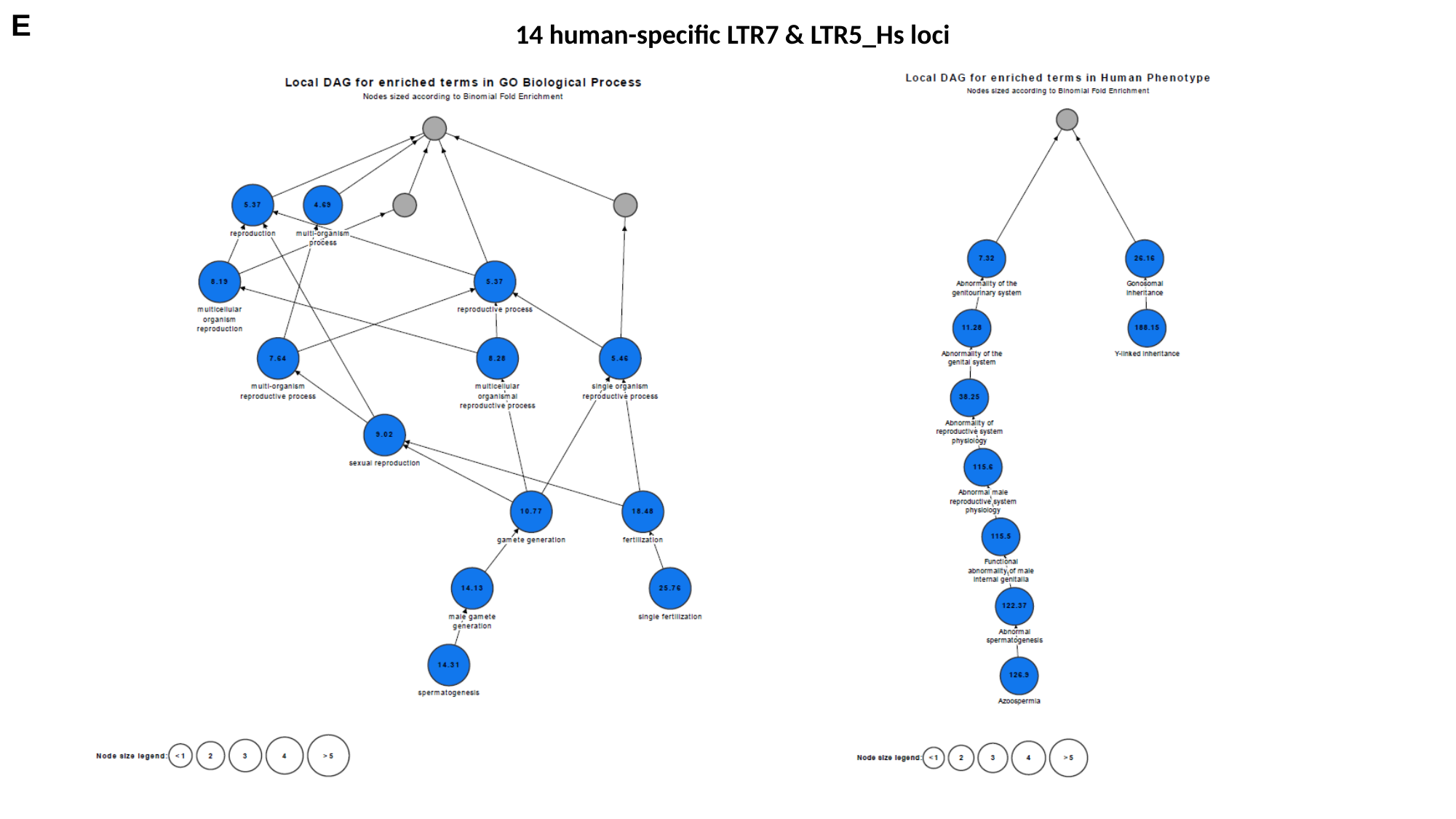

E
14 human-specific LTR7 & LTR5_Hs loci

### Slide 7
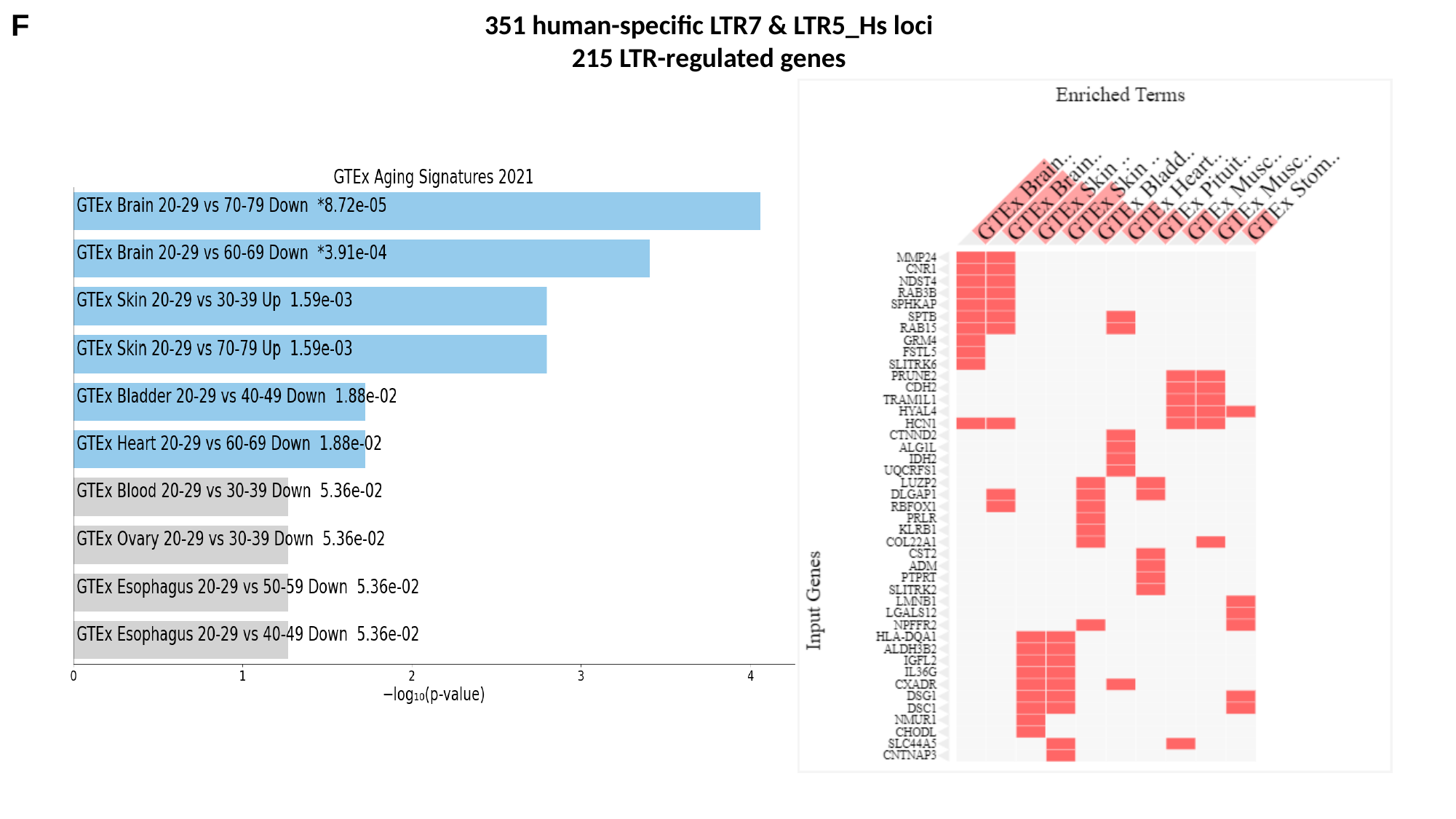

F
351 human-specific LTR7 & LTR5_Hs loci
215 LTR-regulated genes
