## Supplementary for "Molecular diversity and phenotypic pleiotropy of genomic regulatory loci derived from human endogenous retrovirus type H (HERVH) promoter LTR7 and HERVK promoter LTR5_Hs and their impacts on pathophysiology of Modern Humans": Supplementary Figure S2..pptx

### Slide 1
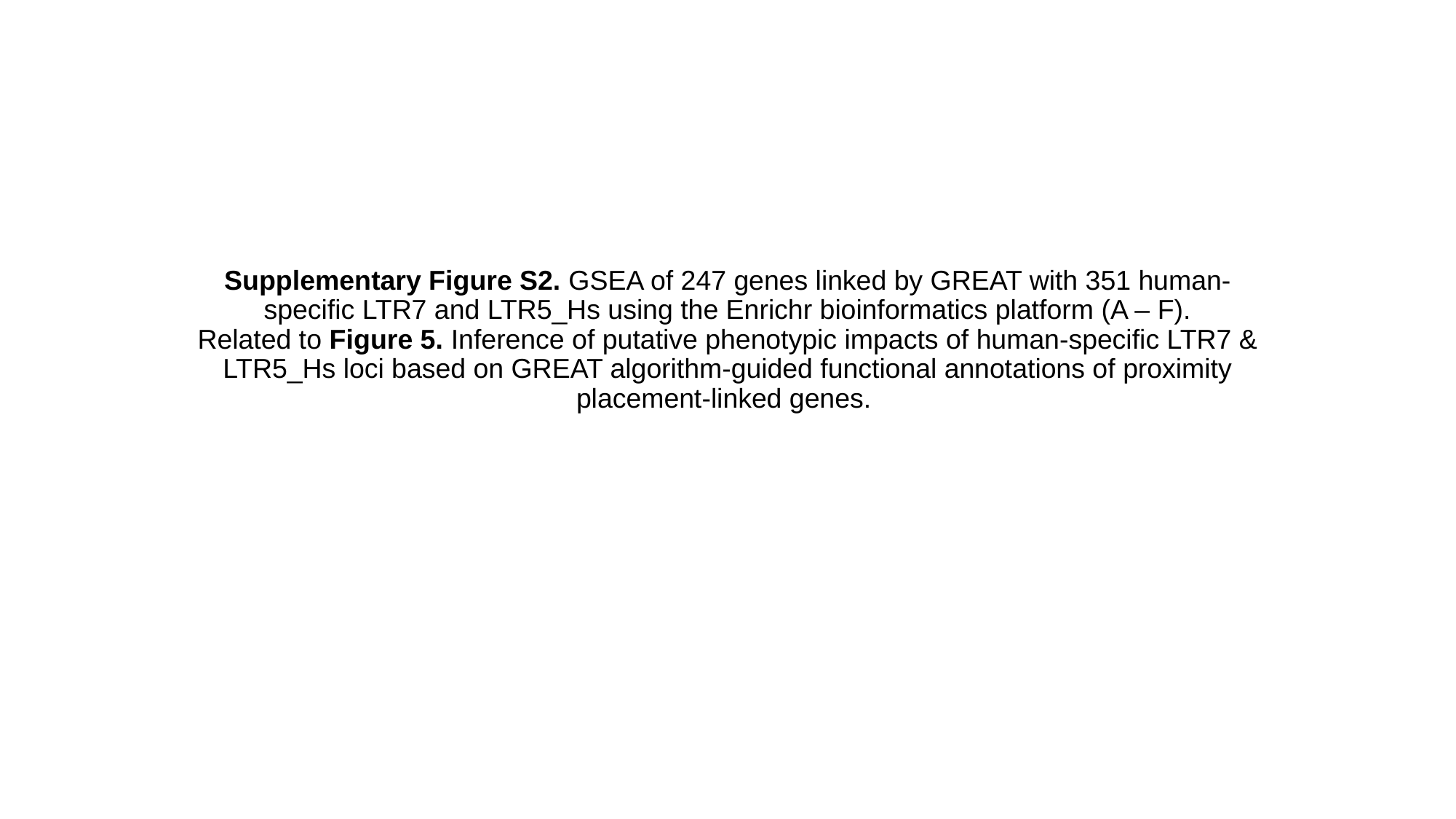

#### Supplementary Figure S2. GSEA of 247 genes linked by GREAT with 351 human-specific LTR7 and LTR5_Hs using the Enrichr bioinformatics platform (A – F). Related to Figure 5. Inference of putative phenotypic impacts of human-specific LTR7 & LTR5_Hs loci based on GREAT algorithm-guided functional annotations of proximity placement-linked genes.

### Slide 2
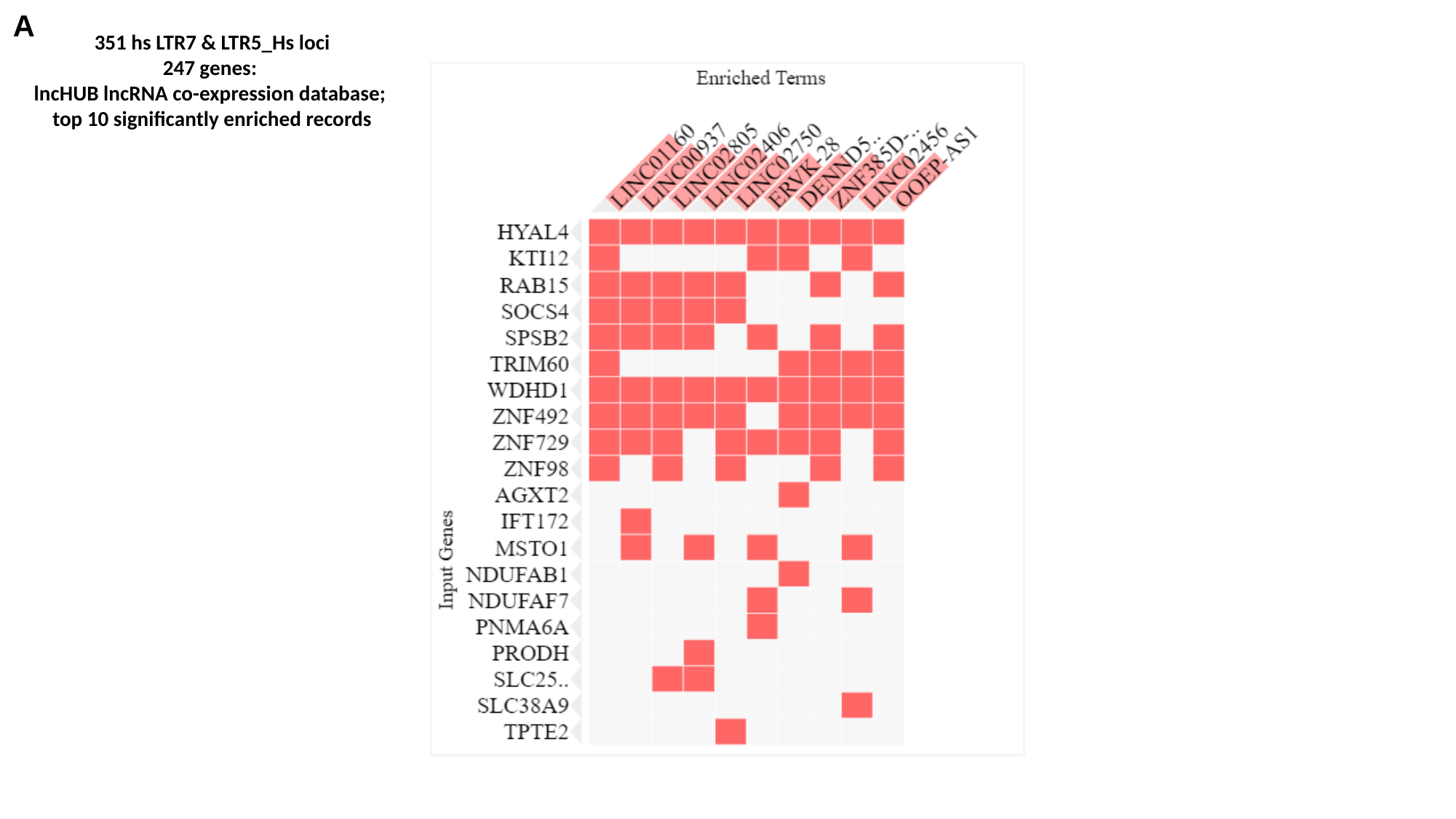

A
351 hs LTR7 & LTR5_Hs loci
247 genes:
lncHUB lncRNA co-expression database;
top 10 significantly enriched records

### Slide 3
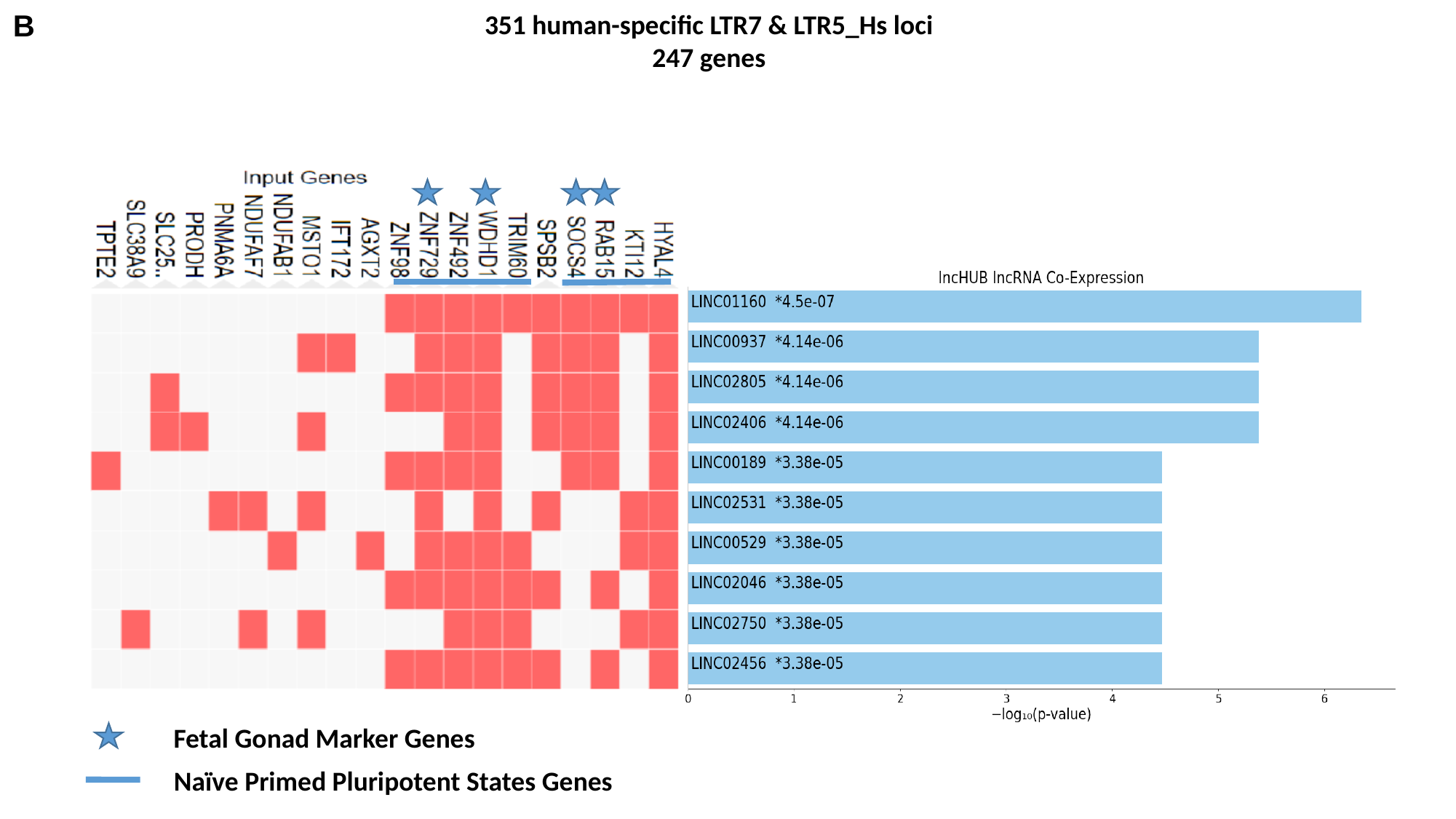

B
351 human-specific LTR7 & LTR5_Hs loci
247 genes
Fetal Gonad Marker Genes
Naïve Primed Pluripotent States Genes

### Slide 4
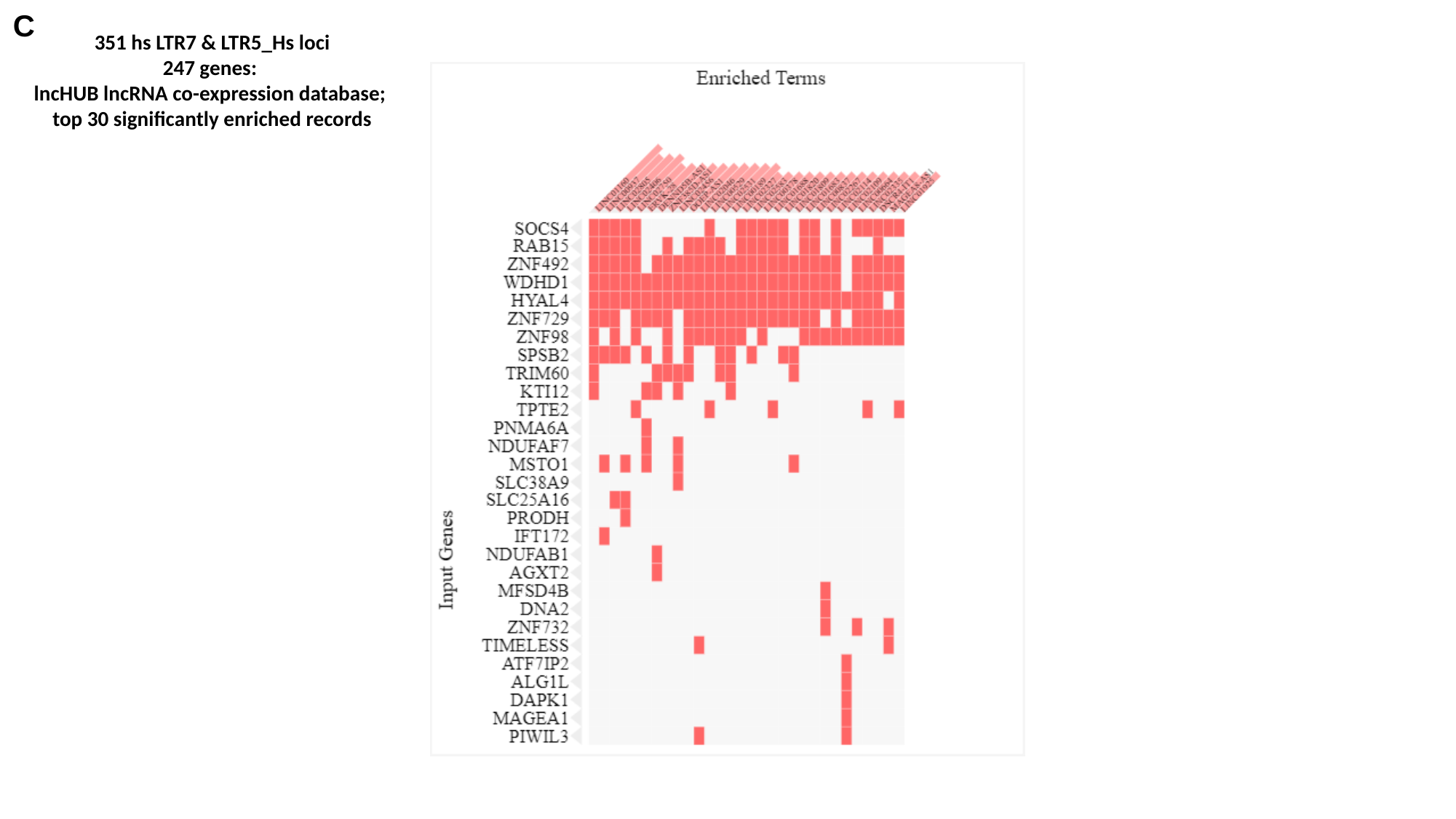

C
351 hs LTR7 & LTR5_Hs loci
247 genes:
lncHUB lncRNA co-expression database;
top 30 significantly enriched records

### Slide 5
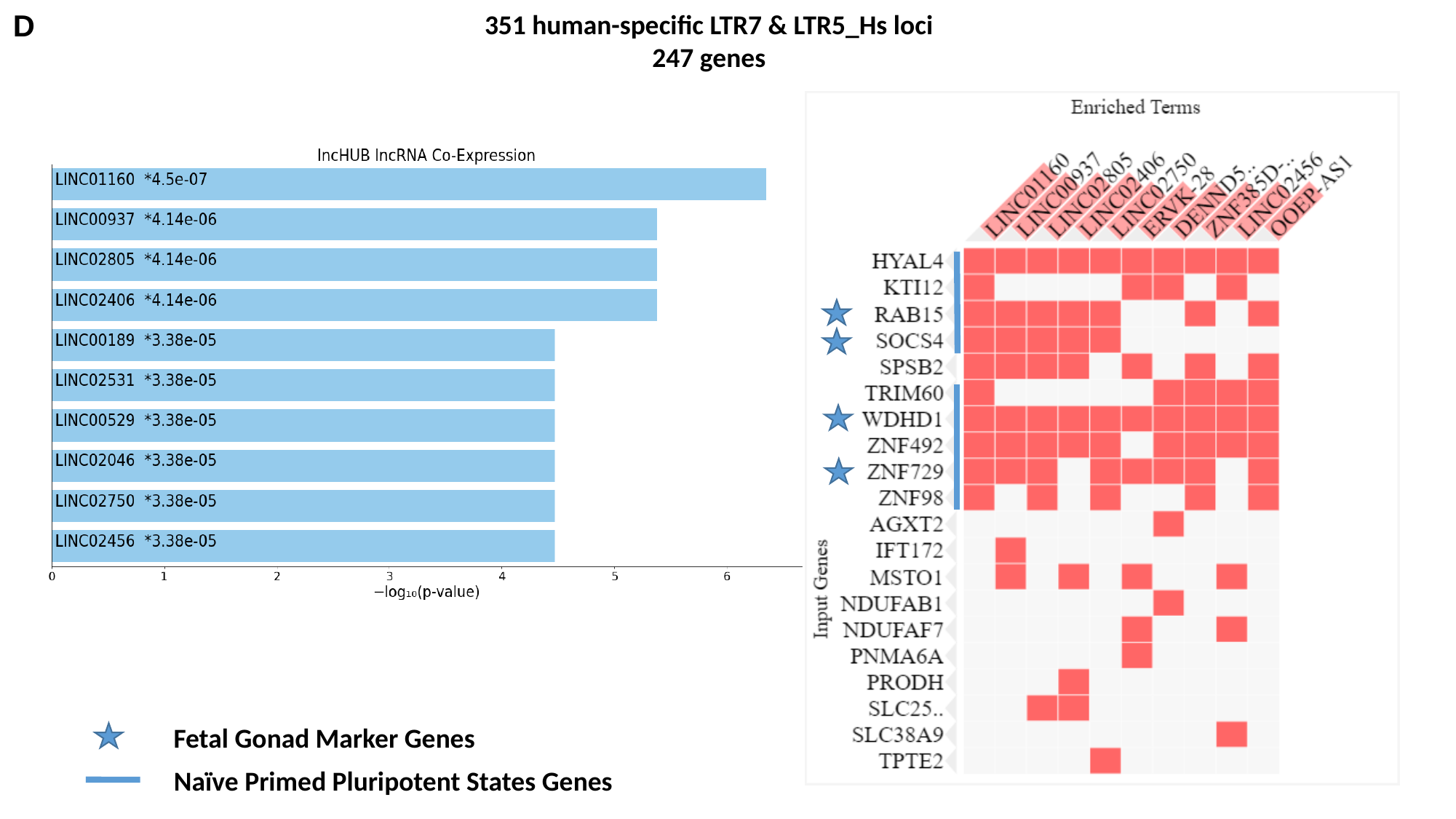

D
351 human-specific LTR7 & LTR5_Hs loci
247 genes
Fetal Gonad Marker Genes
Naïve Primed Pluripotent States Genes

### Slide 6
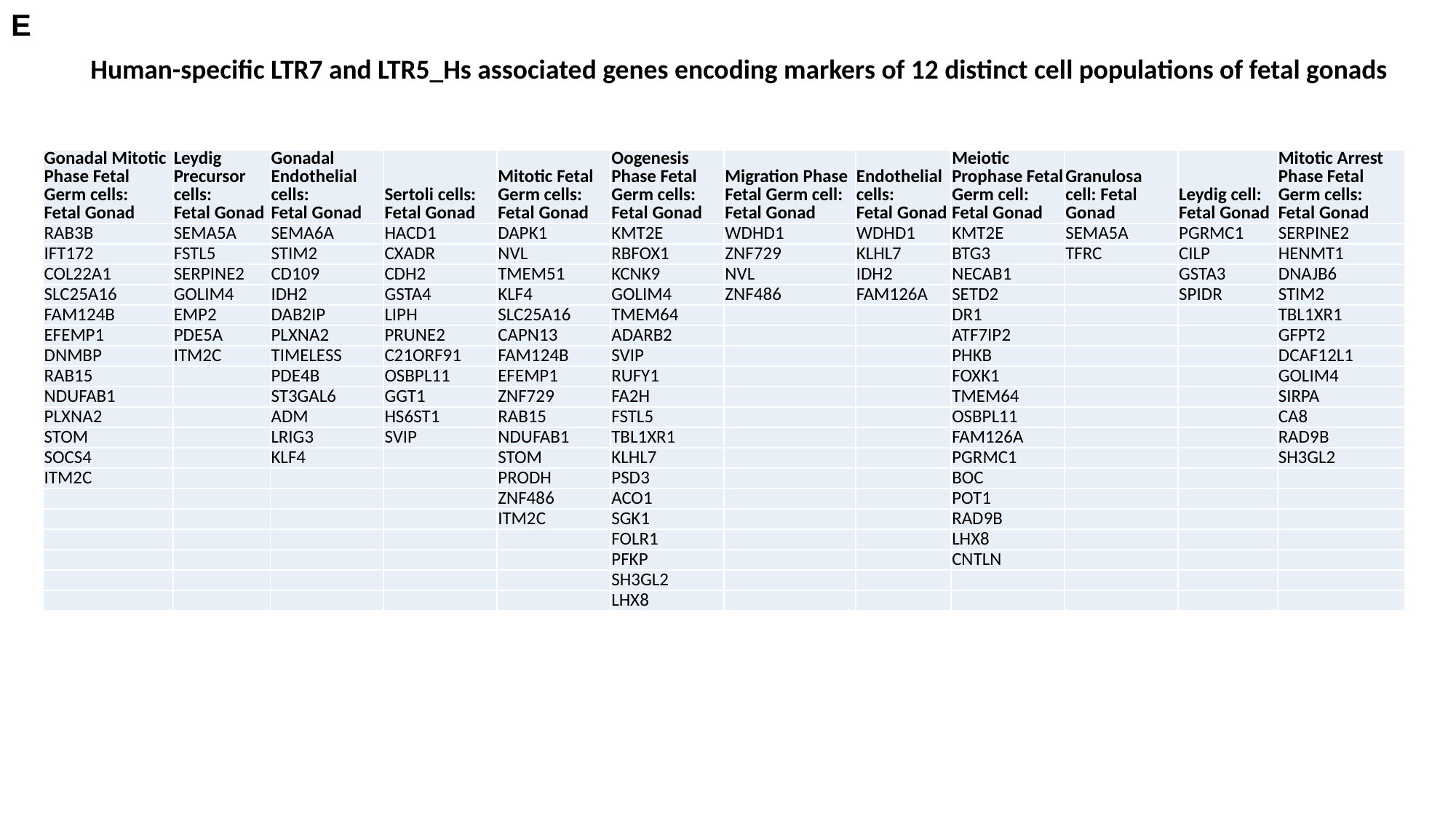

E
Human-specific LTR7 and LTR5_Hs associated genes encoding markers of 12 distinct cell populations of fetal gonads
| Gonadal Mitotic Phase Fetal Germ cells: Fetal Gonad | Leydig Precursor cells: Fetal Gonad | Gonadal Endothelial cells: Fetal Gonad | Sertoli cells: Fetal Gonad | Mitotic Fetal Germ cells: Fetal Gonad | Oogenesis Phase Fetal Germ cells: Fetal Gonad | Migration Phase Fetal Germ cell: Fetal Gonad | Endothelial cells: Fetal Gonad | Meiotic Prophase Fetal Germ cell: Fetal Gonad | Granulosa cell: Fetal Gonad | Leydig cell: Fetal Gonad | Mitotic Arrest Phase Fetal Germ cells: Fetal Gonad |
| --- | --- | --- | --- | --- | --- | --- | --- | --- | --- | --- | --- |
| RAB3B | SEMA5A | SEMA6A | HACD1 | DAPK1 | KMT2E | WDHD1 | WDHD1 | KMT2E | SEMA5A | PGRMC1 | SERPINE2 |
| IFT172 | FSTL5 | STIM2 | CXADR | NVL | RBFOX1 | ZNF729 | KLHL7 | BTG3 | TFRC | CILP | HENMT1 |
| COL22A1 | SERPINE2 | CD109 | CDH2 | TMEM51 | KCNK9 | NVL | IDH2 | NECAB1 | | GSTA3 | DNAJB6 |
| SLC25A16 | GOLIM4 | IDH2 | GSTA4 | KLF4 | GOLIM4 | ZNF486 | FAM126A | SETD2 | | SPIDR | STIM2 |
| FAM124B | EMP2 | DAB2IP | LIPH | SLC25A16 | TMEM64 | | | DR1 | | | TBL1XR1 |
| EFEMP1 | PDE5A | PLXNA2 | PRUNE2 | CAPN13 | ADARB2 | | | ATF7IP2 | | | GFPT2 |
| DNMBP | ITM2C | TIMELESS | C21ORF91 | FAM124B | SVIP | | | PHKB | | | DCAF12L1 |
| RAB15 | | PDE4B | OSBPL11 | EFEMP1 | RUFY1 | | | FOXK1 | | | GOLIM4 |
| NDUFAB1 | | ST3GAL6 | GGT1 | ZNF729 | FA2H | | | TMEM64 | | | SIRPA |
| PLXNA2 | | ADM | HS6ST1 | RAB15 | FSTL5 | | | OSBPL11 | | | CA8 |
| STOM | | LRIG3 | SVIP | NDUFAB1 | TBL1XR1 | | | FAM126A | | | RAD9B |
| SOCS4 | | KLF4 | | STOM | KLHL7 | | | PGRMC1 | | | SH3GL2 |
| ITM2C | | | | PRODH | PSD3 | | | BOC | | | |
| | | | | ZNF486 | ACO1 | | | POT1 | | | |
| | | | | ITM2C | SGK1 | | | RAD9B | | | |
| | | | | | FOLR1 | | | LHX8 | | | |
| | | | | | PFKP | | | CNTLN | | | |
| | | | | | SH3GL2 | | | | | | |
| | | | | | LHX8 | | | | | | |

### Slide 7
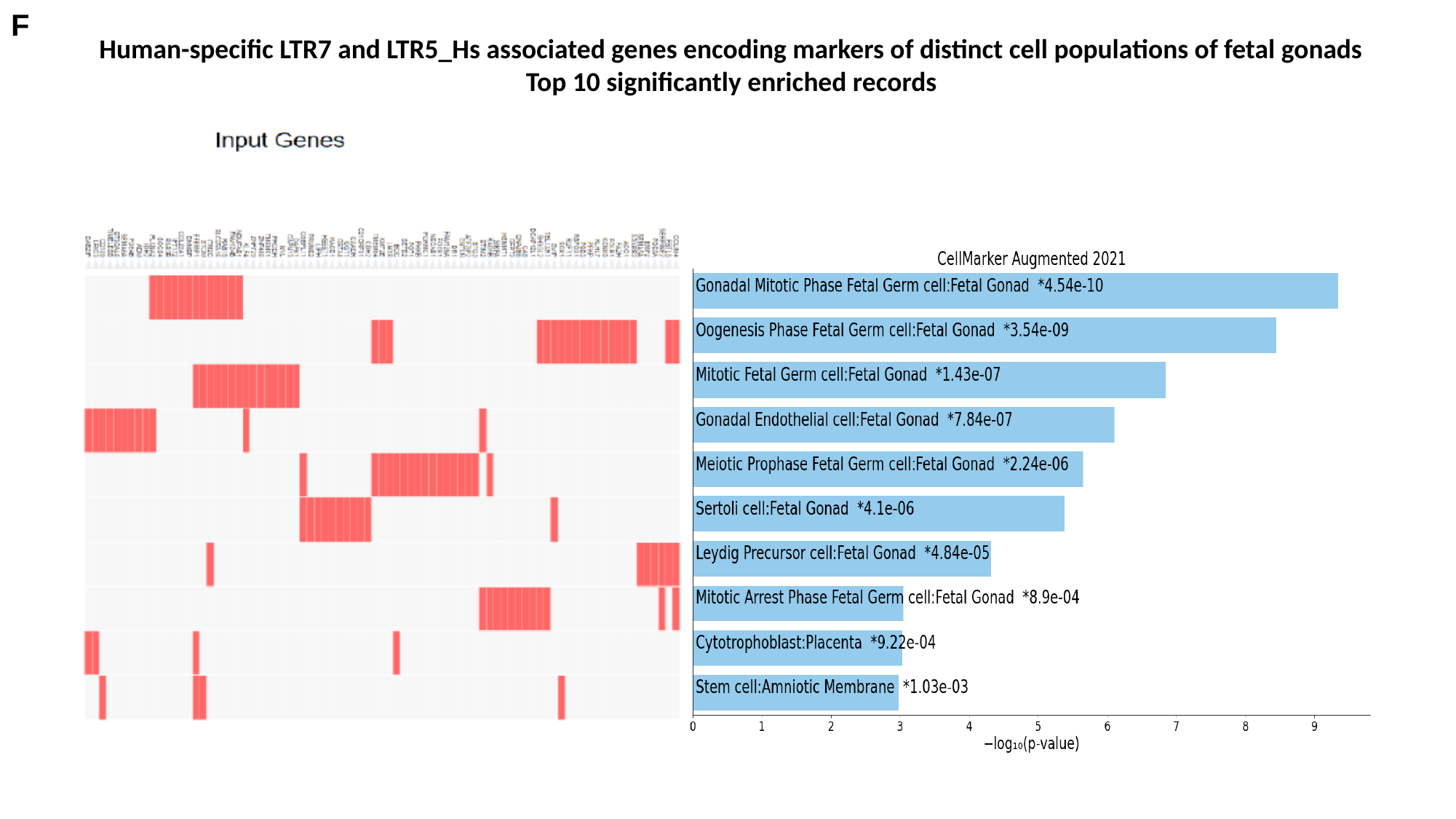

F
Human-specific LTR7 and LTR5_Hs associated genes encoding markers of distinct cell populations of fetal gonads
Top 10 significantly enriched records
