## Supplementary for "Molecular diversity and phenotypic pleiotropy of genomic regulatory loci derived from human endogenous retrovirus type H (HERVH) promoter LTR7 and HERVK promoter LTR5_Hs and their impacts on pathophysiology of Modern Humans": Supplementary Figure S3. 377 LTR7-linked genes among 935 LTR5_Hs linked genes.pptx

### Slide 1
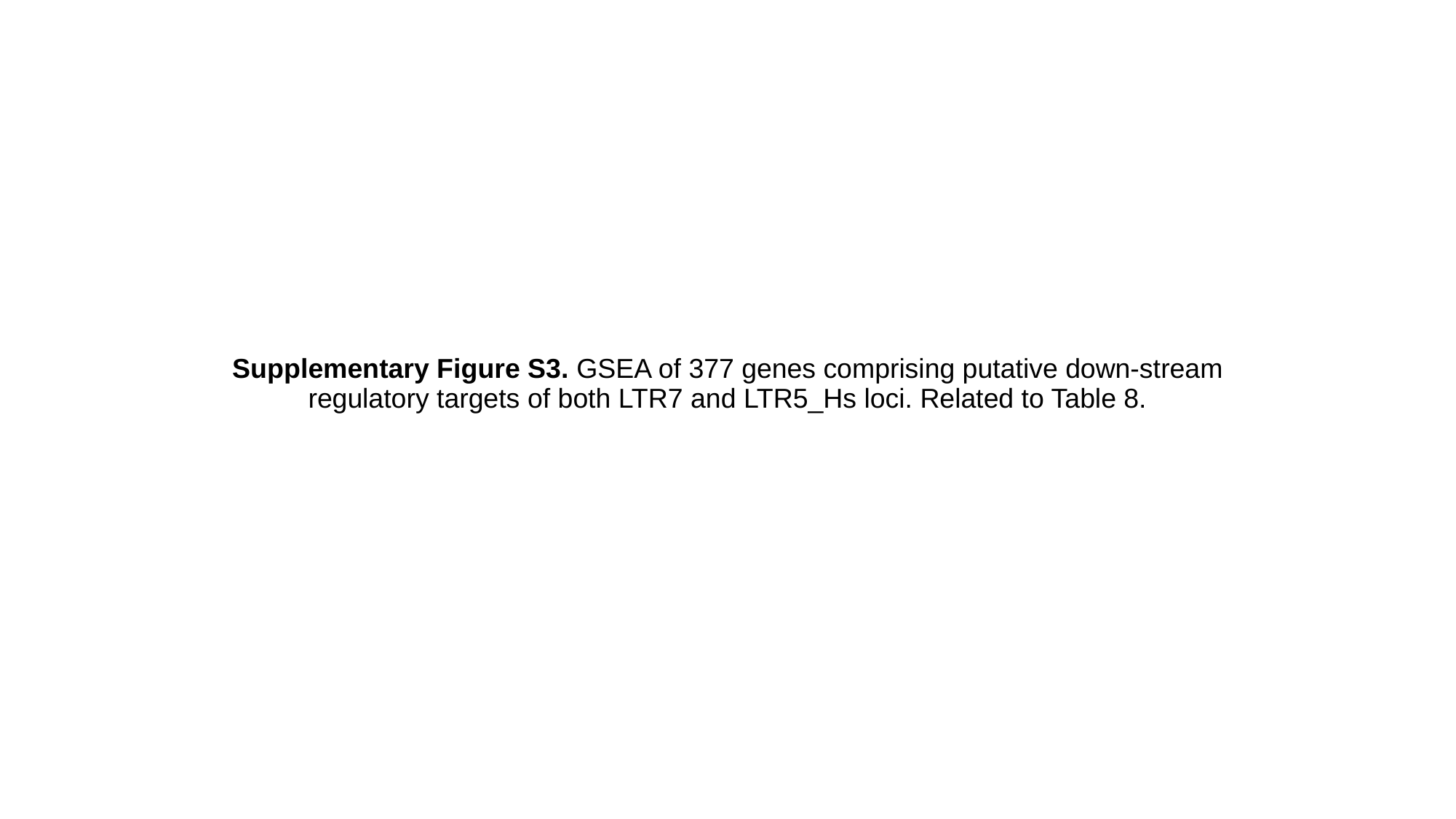

#### Supplementary Figure S3. GSEA of 377 genes comprising putative down-stream regulatory targets of both LTR7 and LTR5_Hs loci. Related to Table 8.

### Slide 2
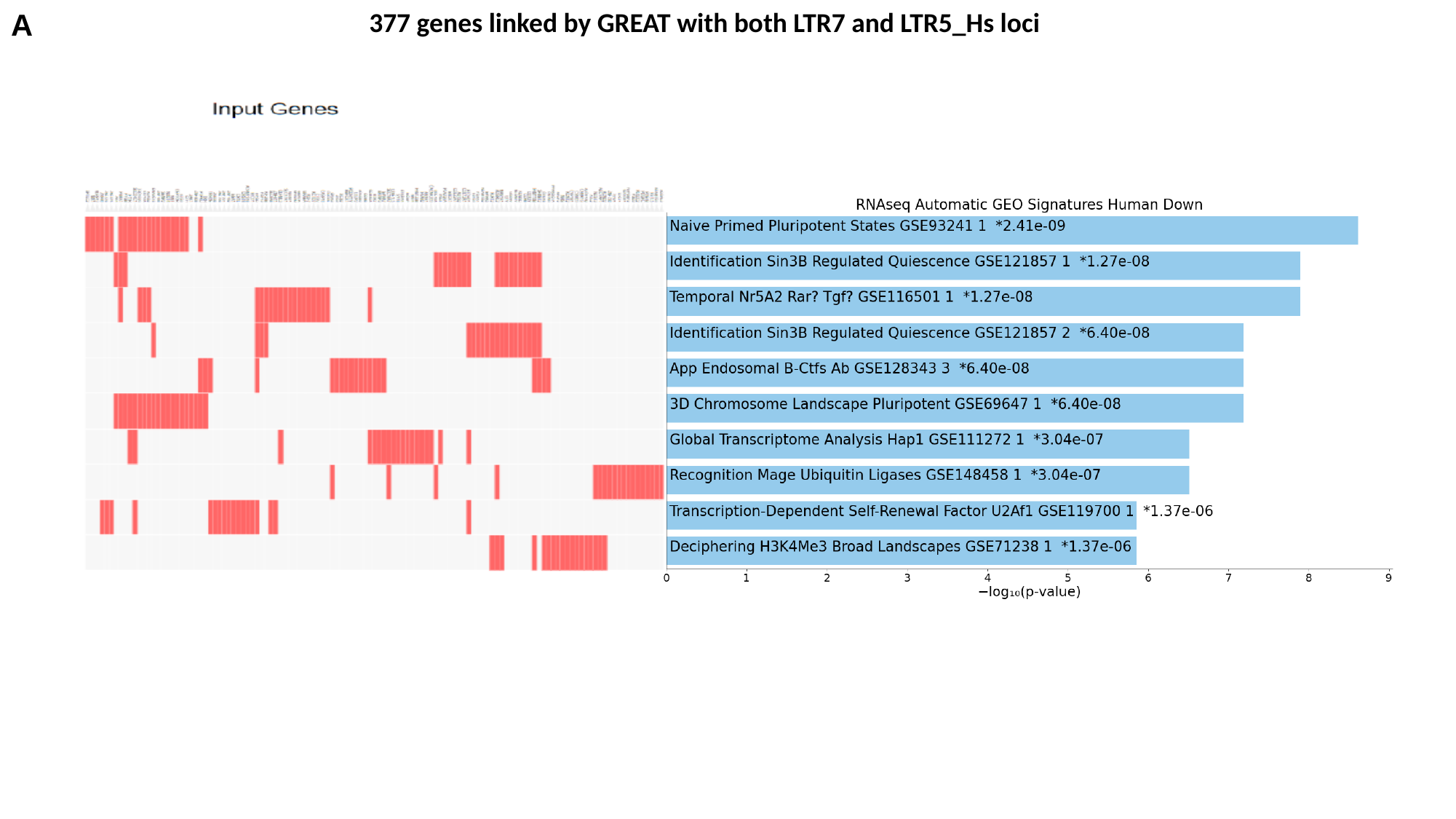

A
377 genes linked by GREAT with both LTR7 and LTR5_Hs loci

### Slide 3
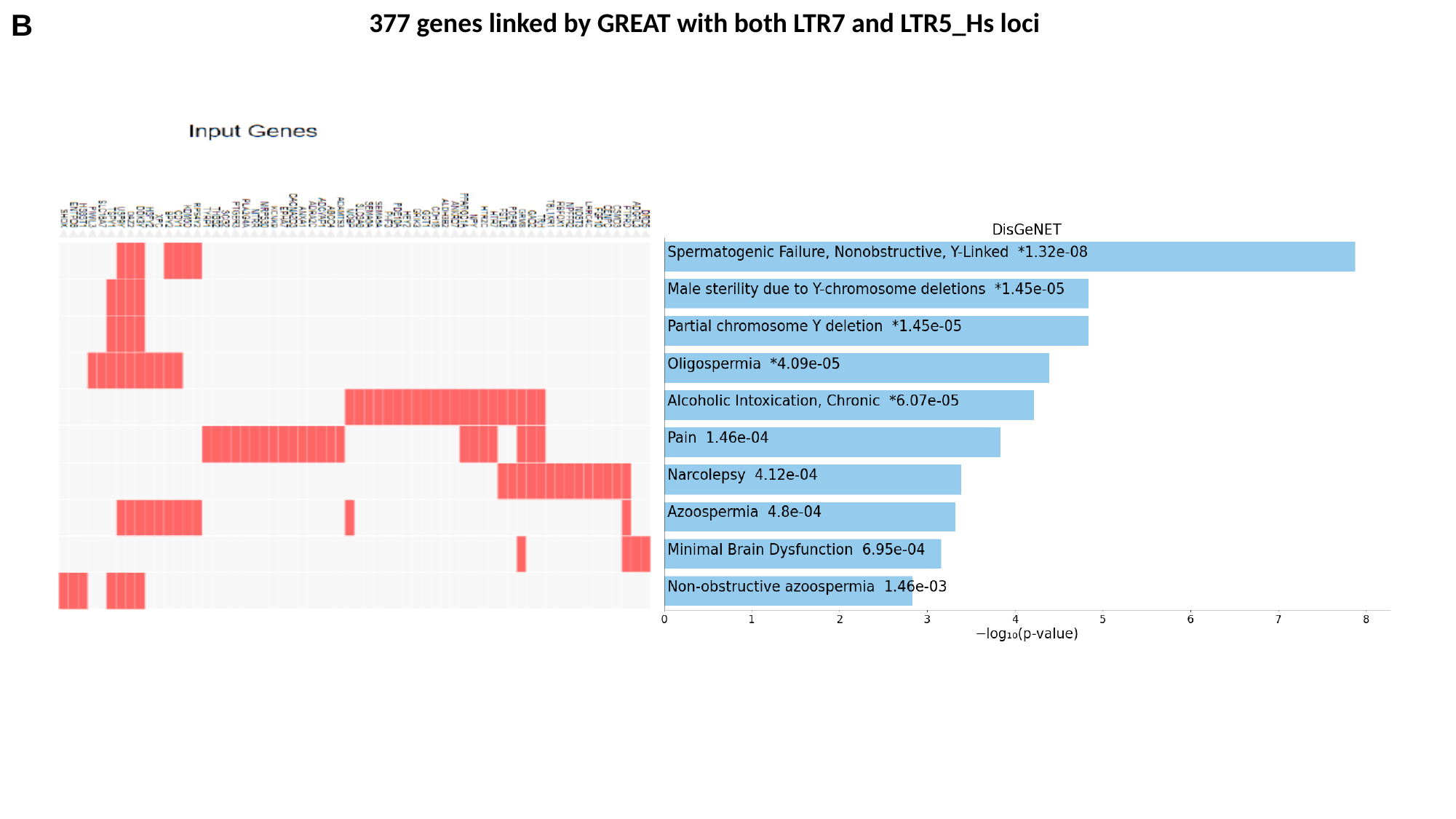

B
377 genes linked by GREAT with both LTR7 and LTR5_Hs loci

### Slide 4
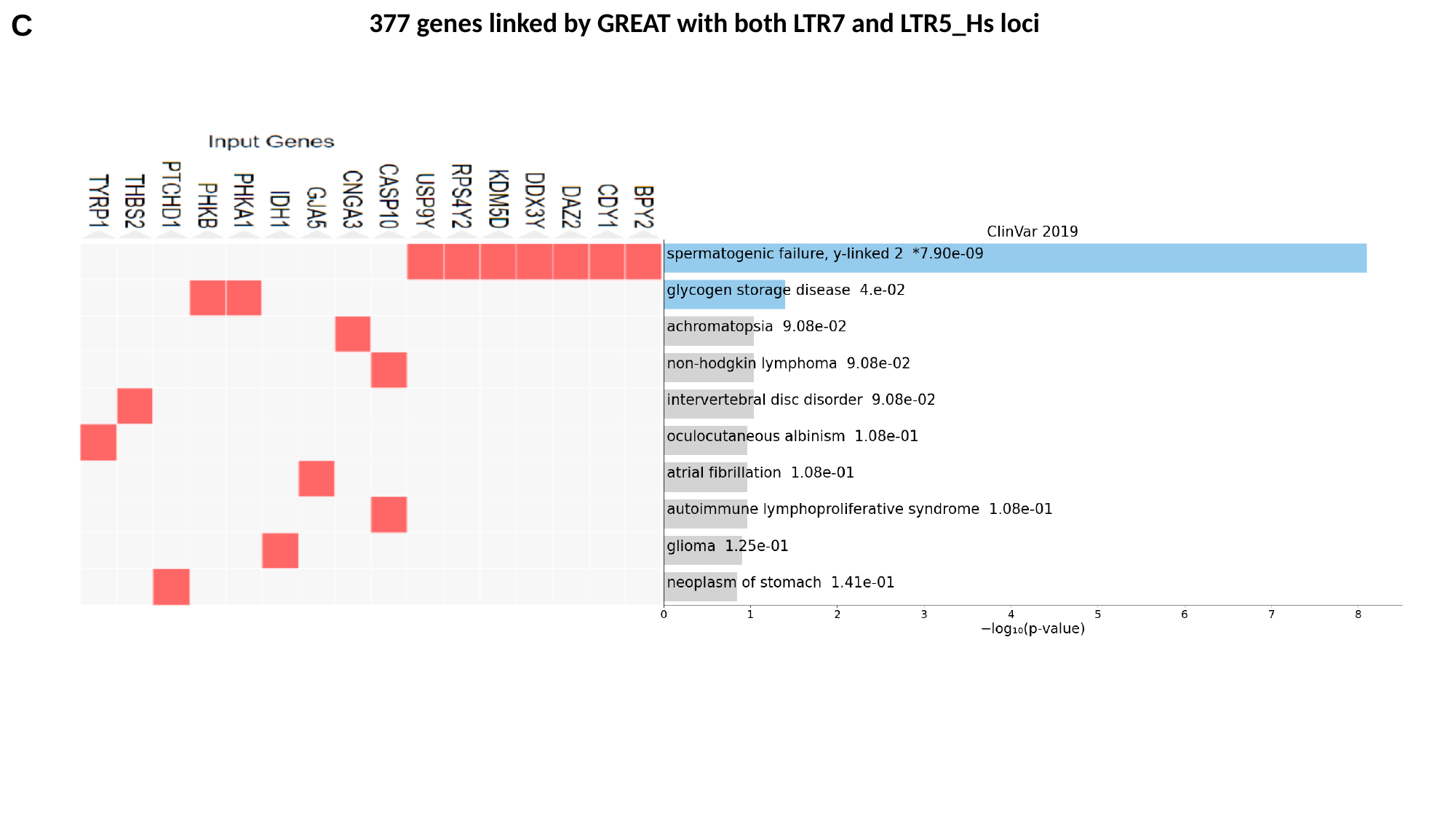

C
377 genes linked by GREAT with both LTR7 and LTR5_Hs loci

### Slide 5
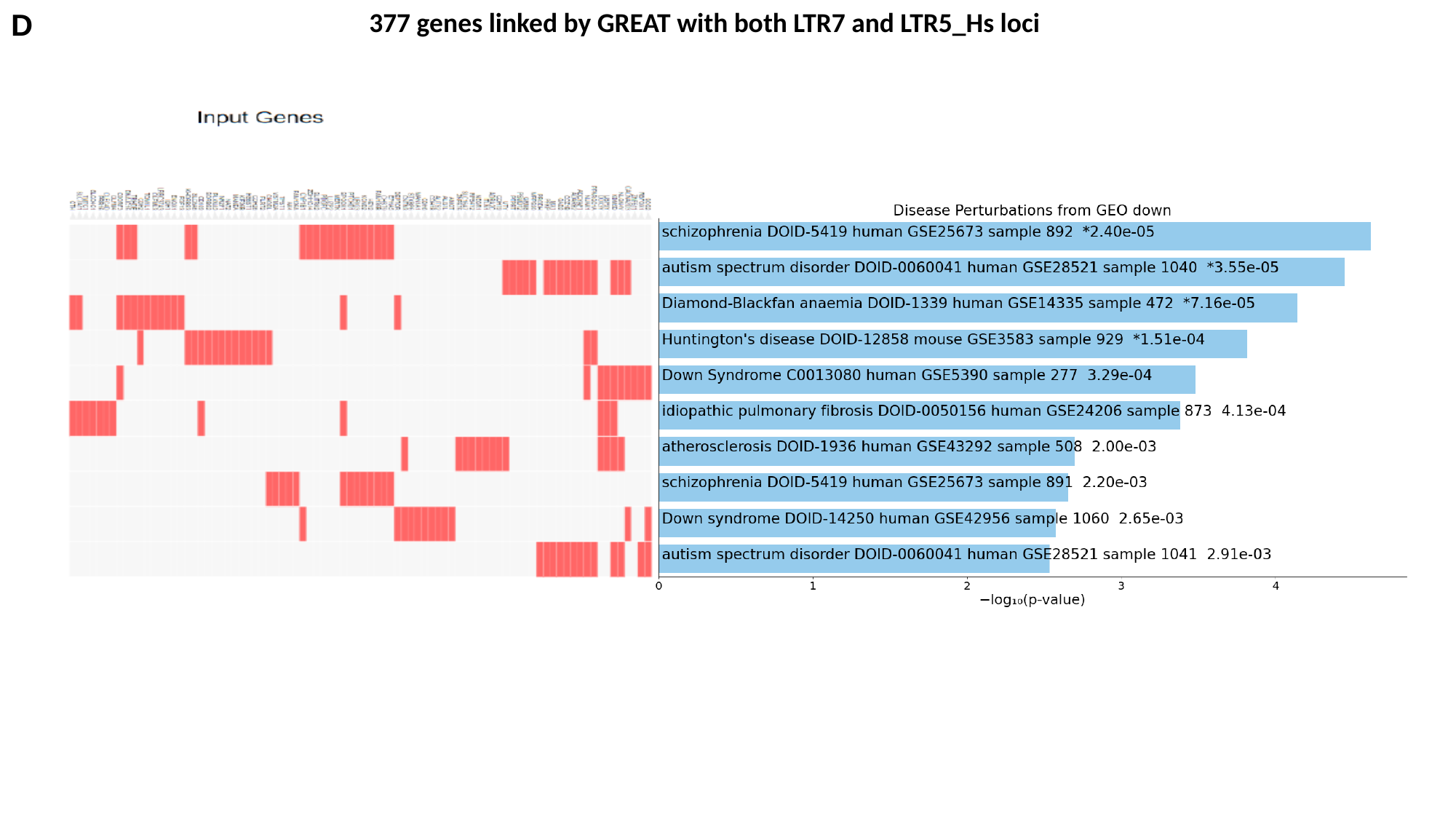

D
377 genes linked by GREAT with both LTR7 and LTR5_Hs loci
