## Supplementary for "Molecular diversity and phenotypic pleiotropy of genomic regulatory loci derived from human endogenous retrovirus type H (HERVH) promoter LTR7 and HERVK promoter LTR5_Hs and their impacts on pathophysiology of Modern Humans": Supplementary Figure S4. 16 TF genes 377 LTR7-linked genes among 935 LTR5_Hs linked genes.pptx

### Slide 1
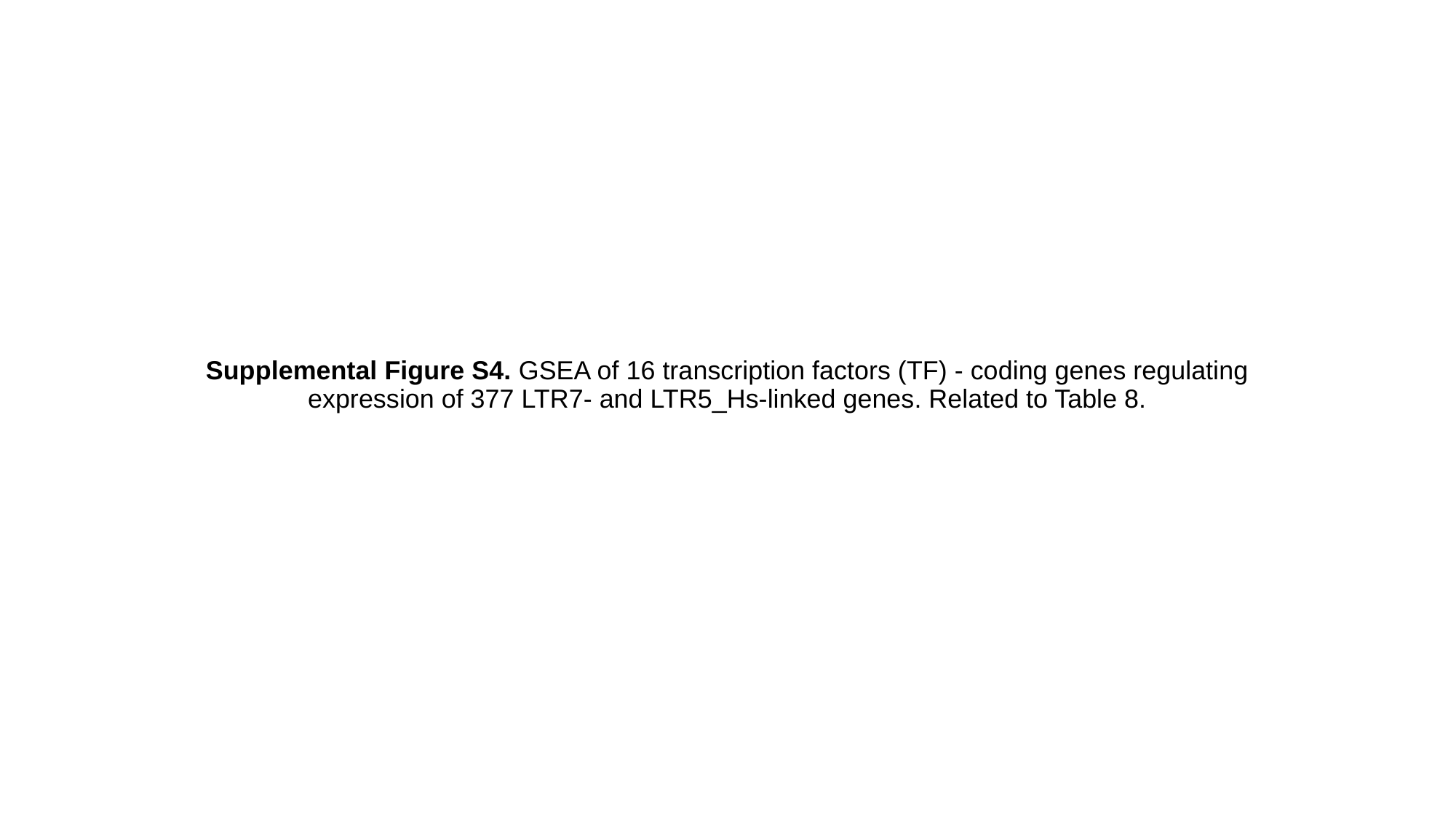

#### Supplemental Figure S4. GSEA of 16 transcription factors (TF) - coding genes regulating expression of 377 LTR7- and LTR5_Hs-linked genes. Related to Table 8.

### Slide 2
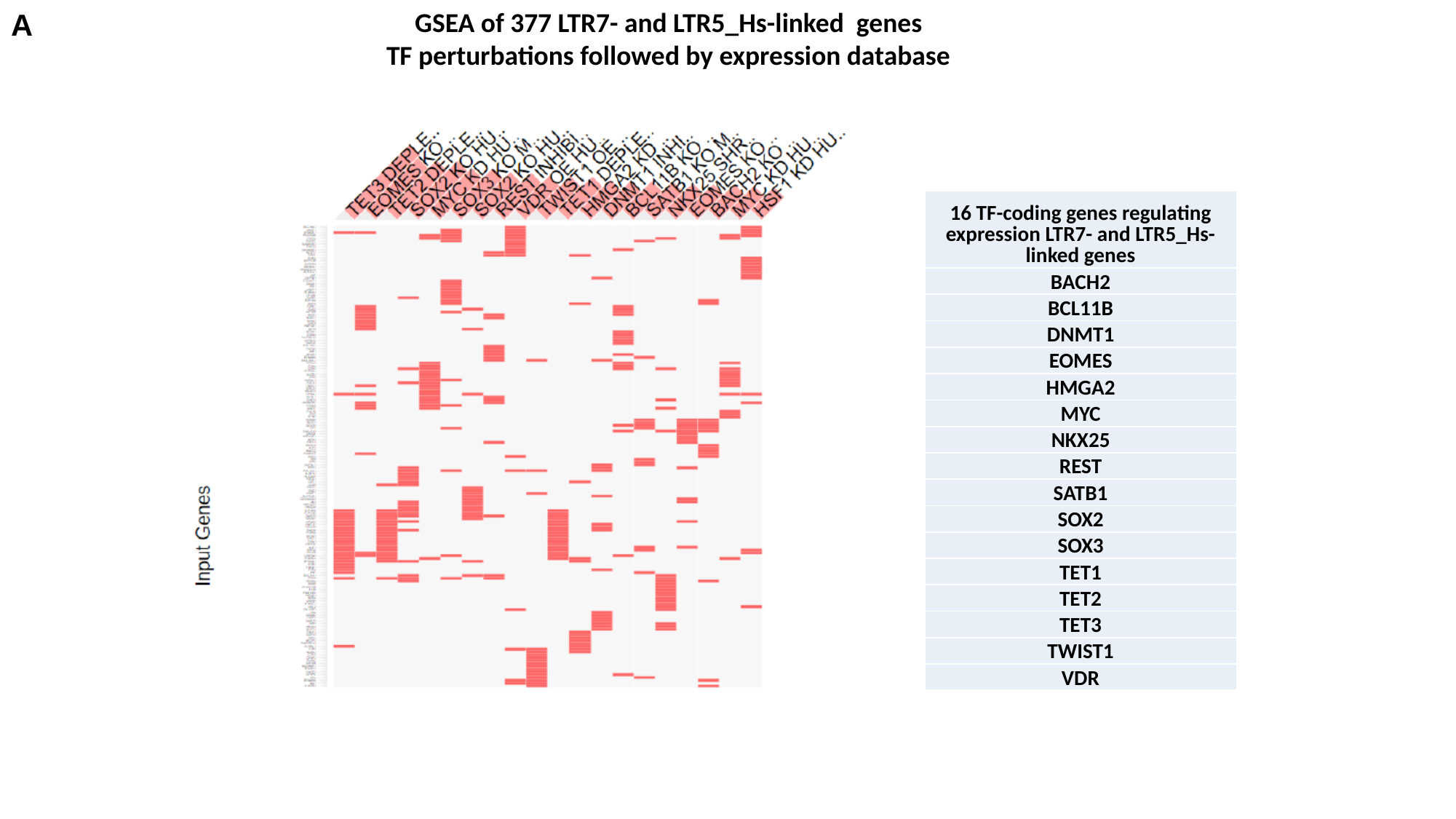

A
GSEA of 377 LTR7- and LTR5_Hs-linked genes
TF perturbations followed by expression database
| 16 TF-coding genes regulating expression LTR7- and LTR5\_Hs-linked genes |
| --- |
| BACH2 |
| BCL11B |
| DNMT1 |
| EOMES |
| HMGA2 |
| MYC |
| NKX25 |
| REST |
| SATB1 |
| SOX2 |
| SOX3 |
| TET1 |
| TET2 |
| TET3 |
| TWIST1 |
| VDR |

### Slide 3
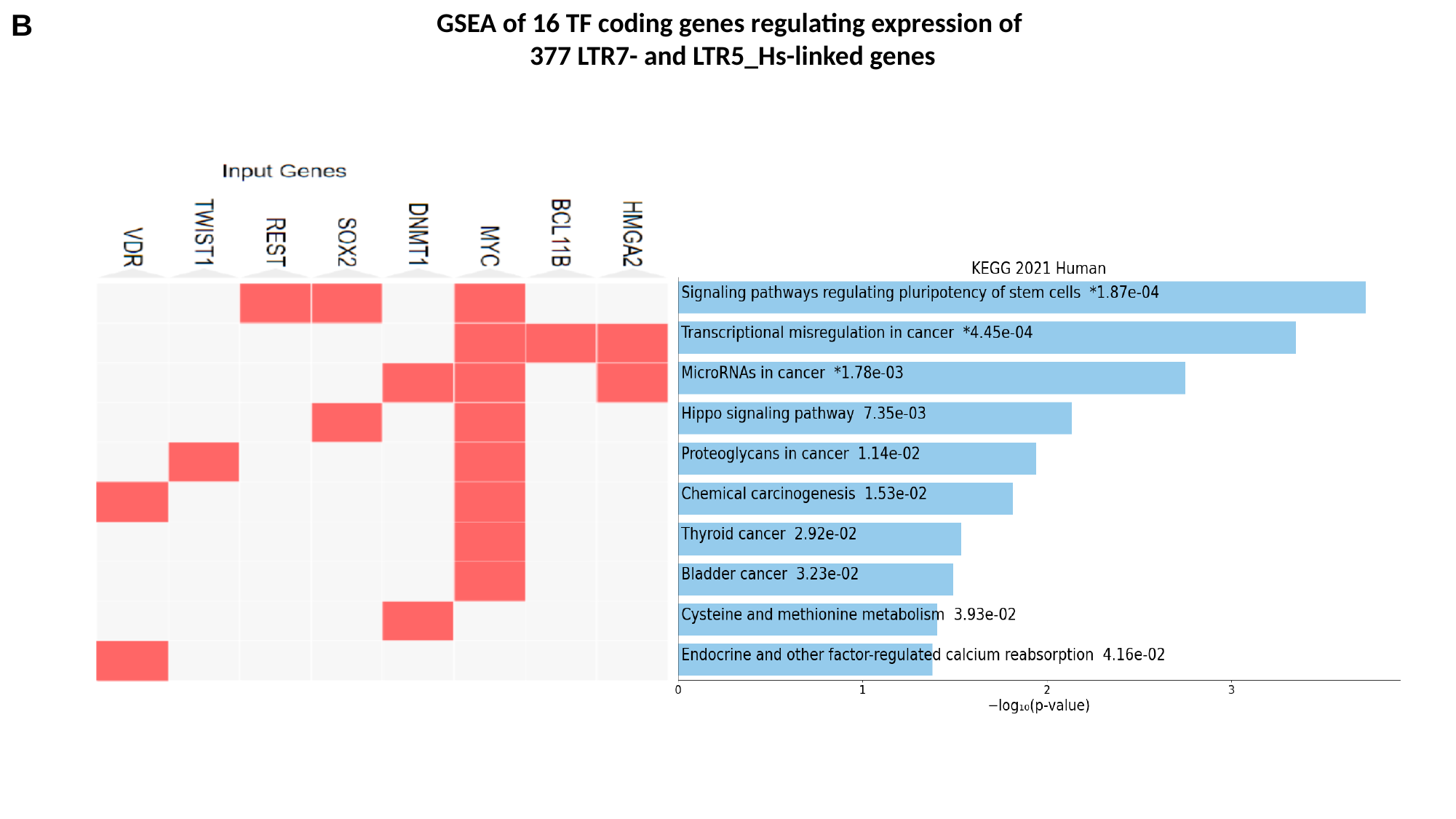

B
GSEA of 16 TF coding genes regulating expression of
377 LTR7- and LTR5_Hs-linked genes

### Slide 4
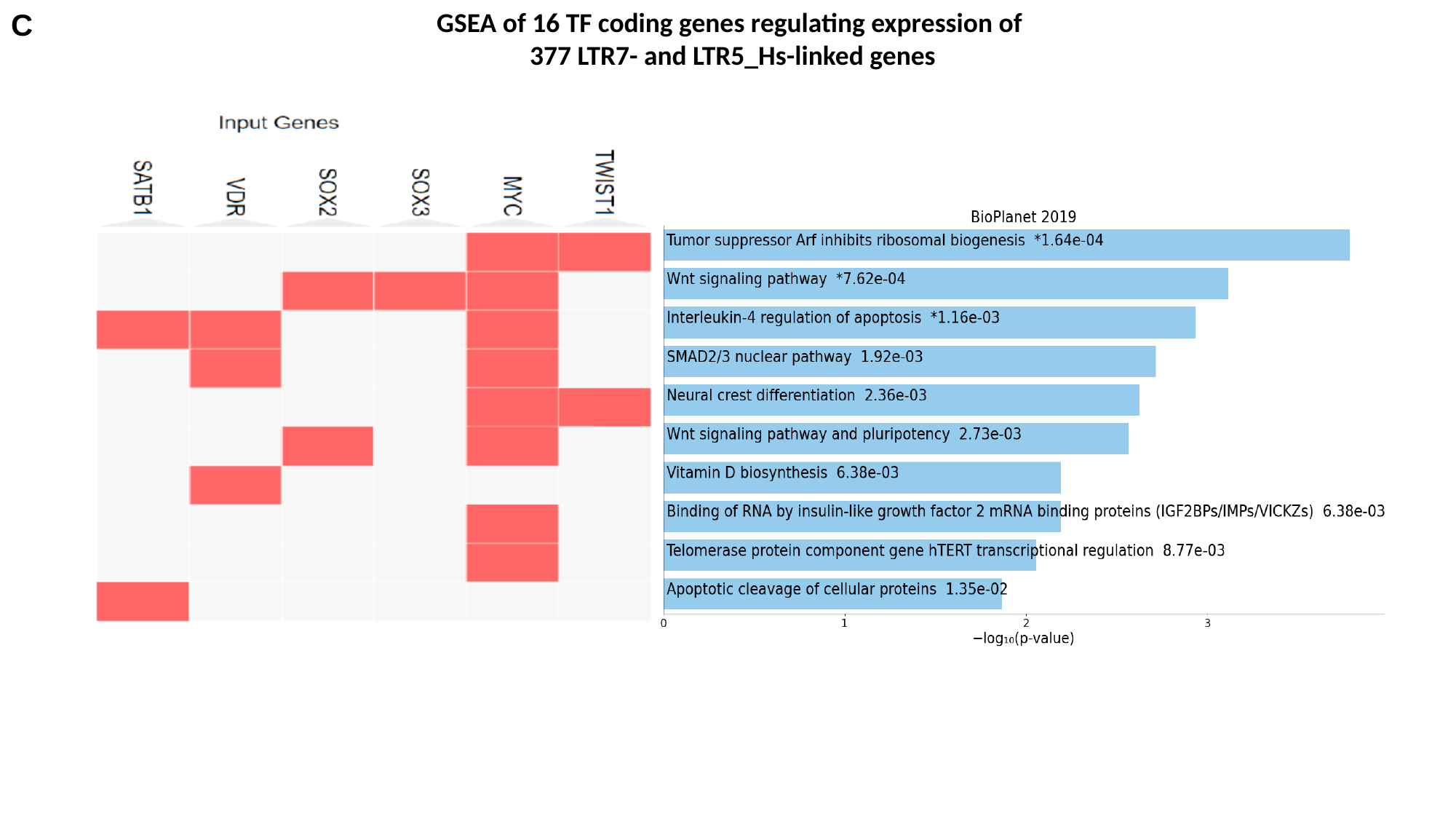

C
GSEA of 16 TF coding genes regulating expression of
377 LTR7- and LTR5_Hs-linked genes

### Slide 5
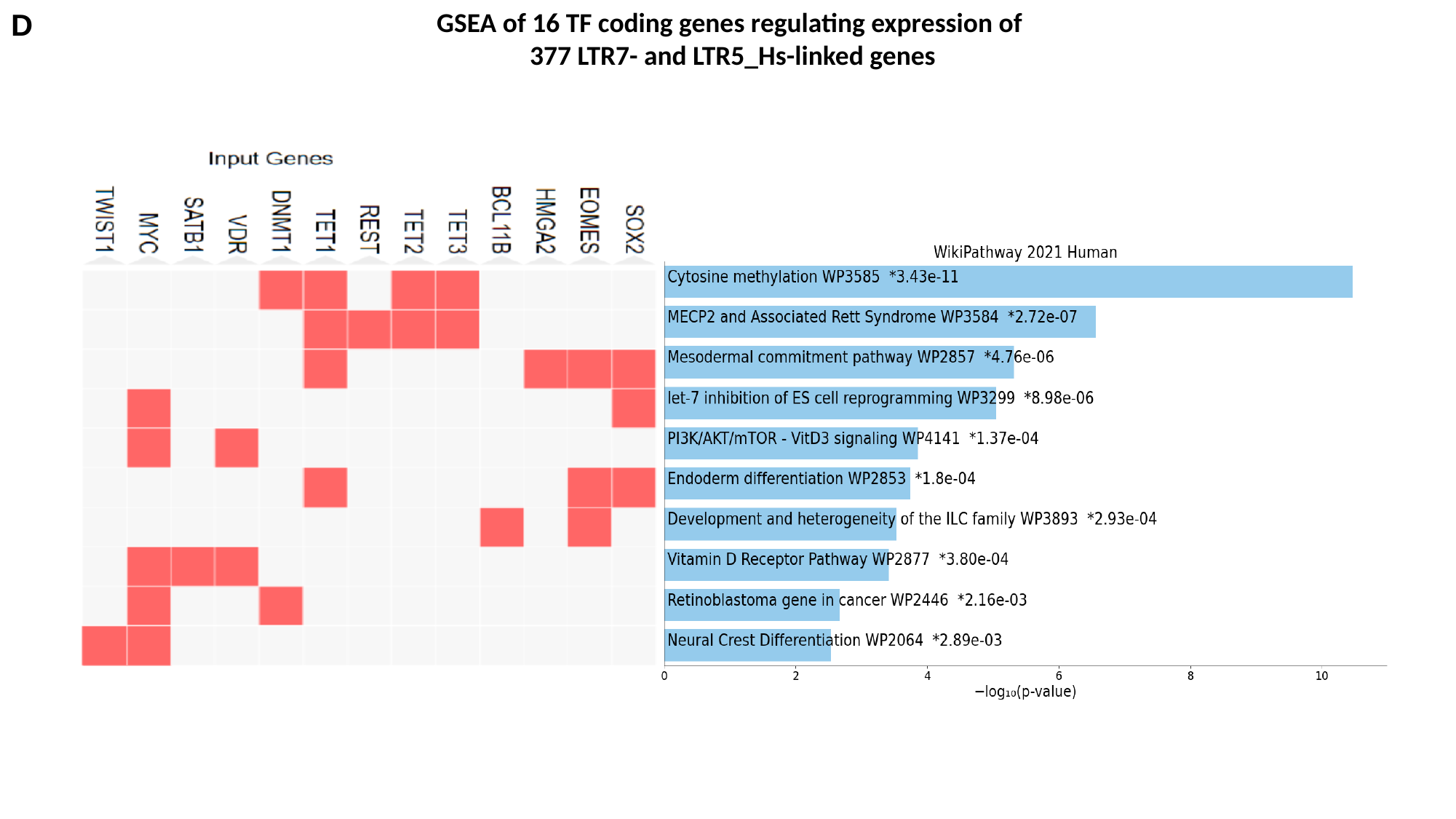

D
GSEA of 16 TF coding genes regulating expression of
377 LTR7- and LTR5_Hs-linked genes

### Slide 6
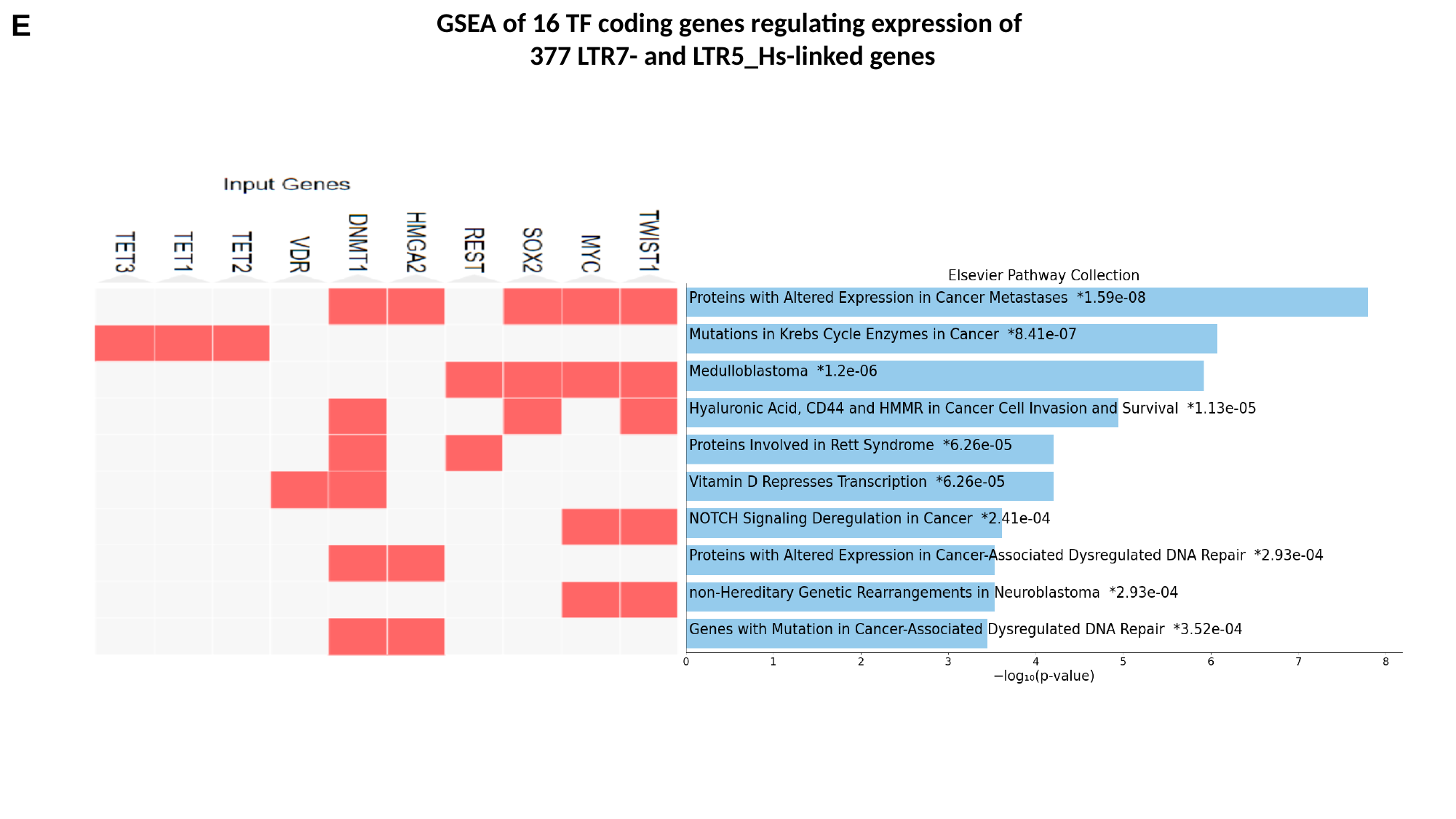

E
GSEA of 16 TF coding genes regulating expression of
377 LTR7- and LTR5_Hs-linked genes

### Slide 7
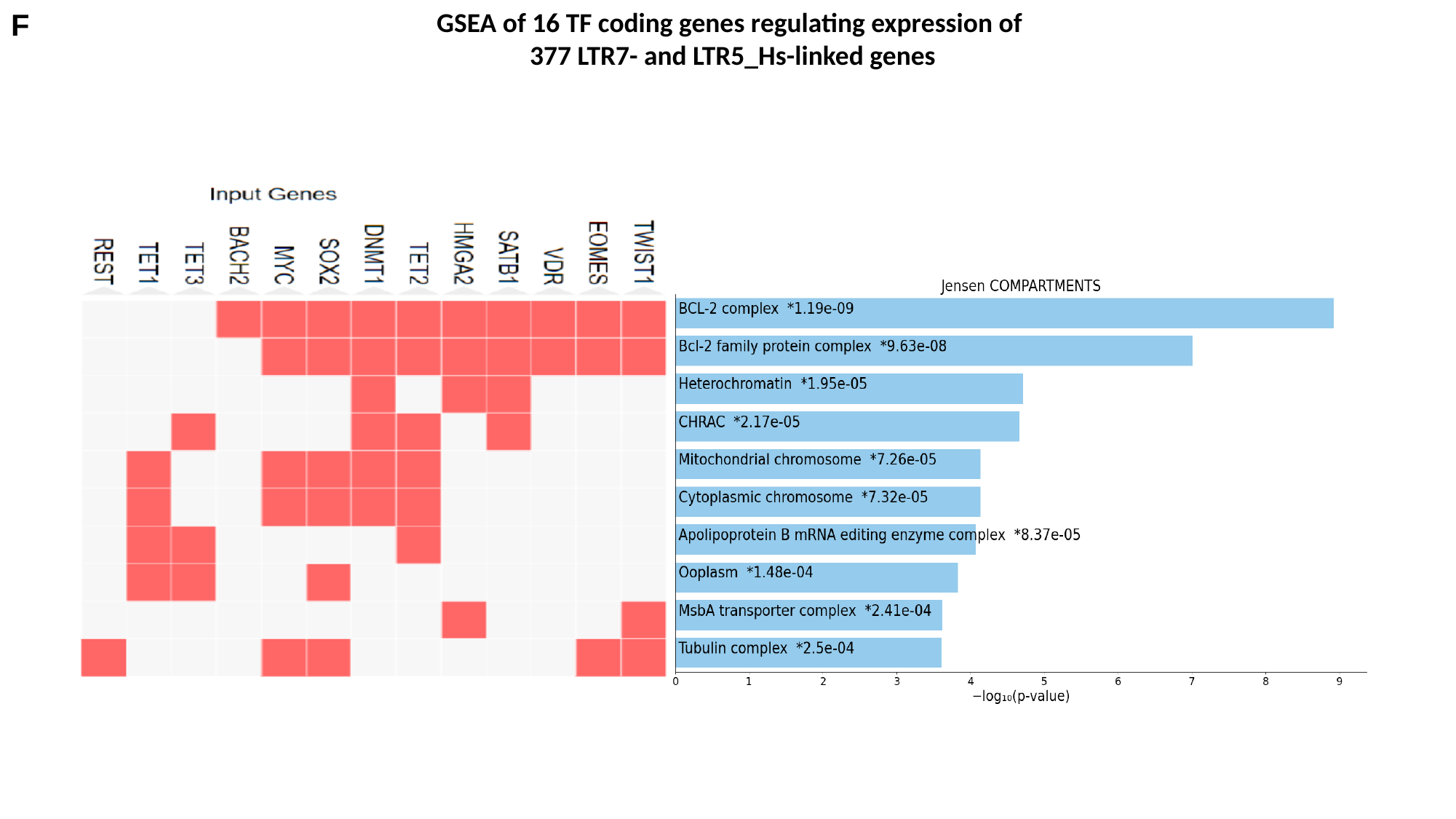

F
GSEA of 16 TF coding genes regulating expression of
377 LTR7- and LTR5_Hs-linked genes

### Slide 8
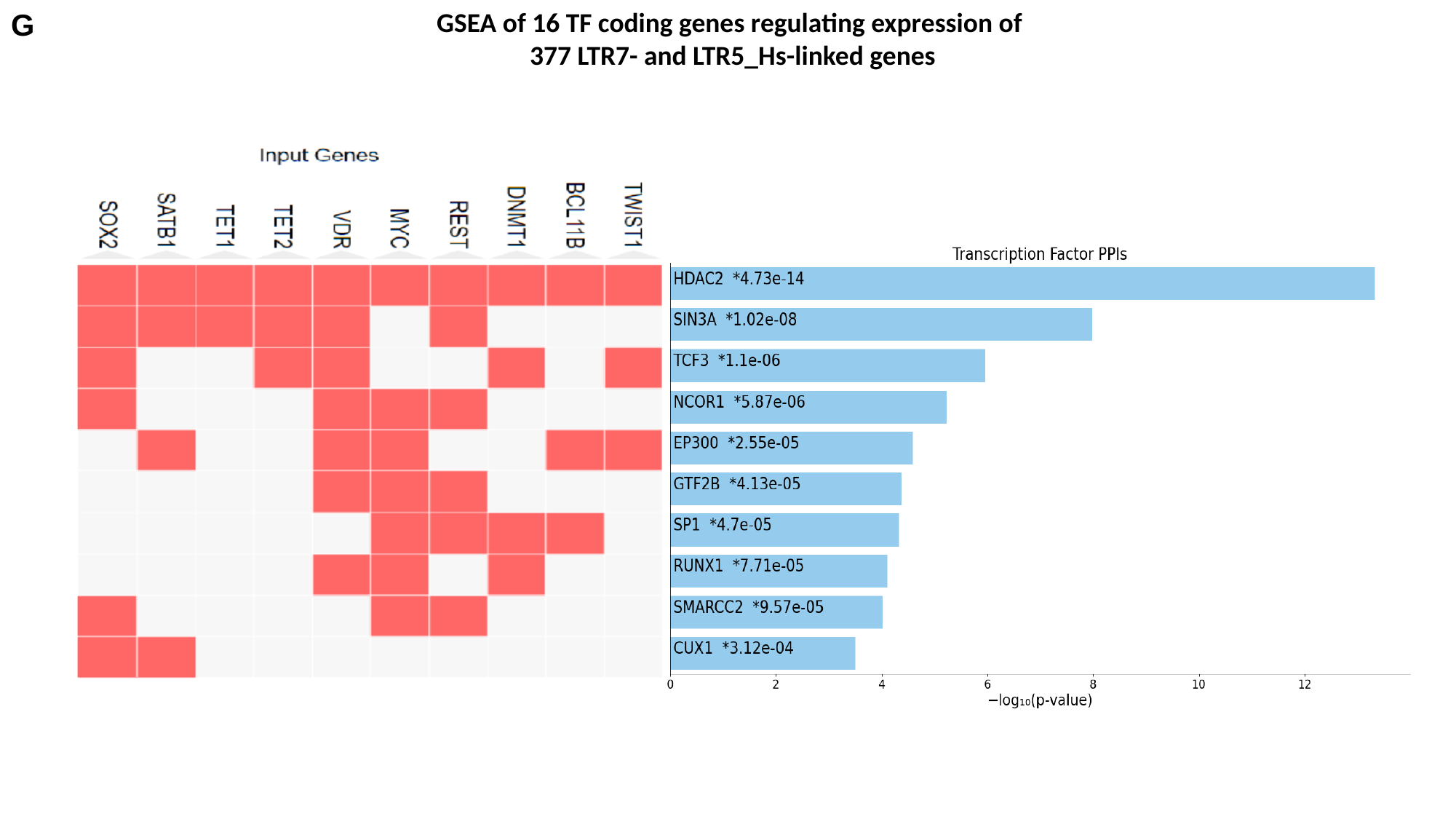

G
GSEA of 16 TF coding genes regulating expression of
377 LTR7- and LTR5_Hs-linked genes

### Slide 9
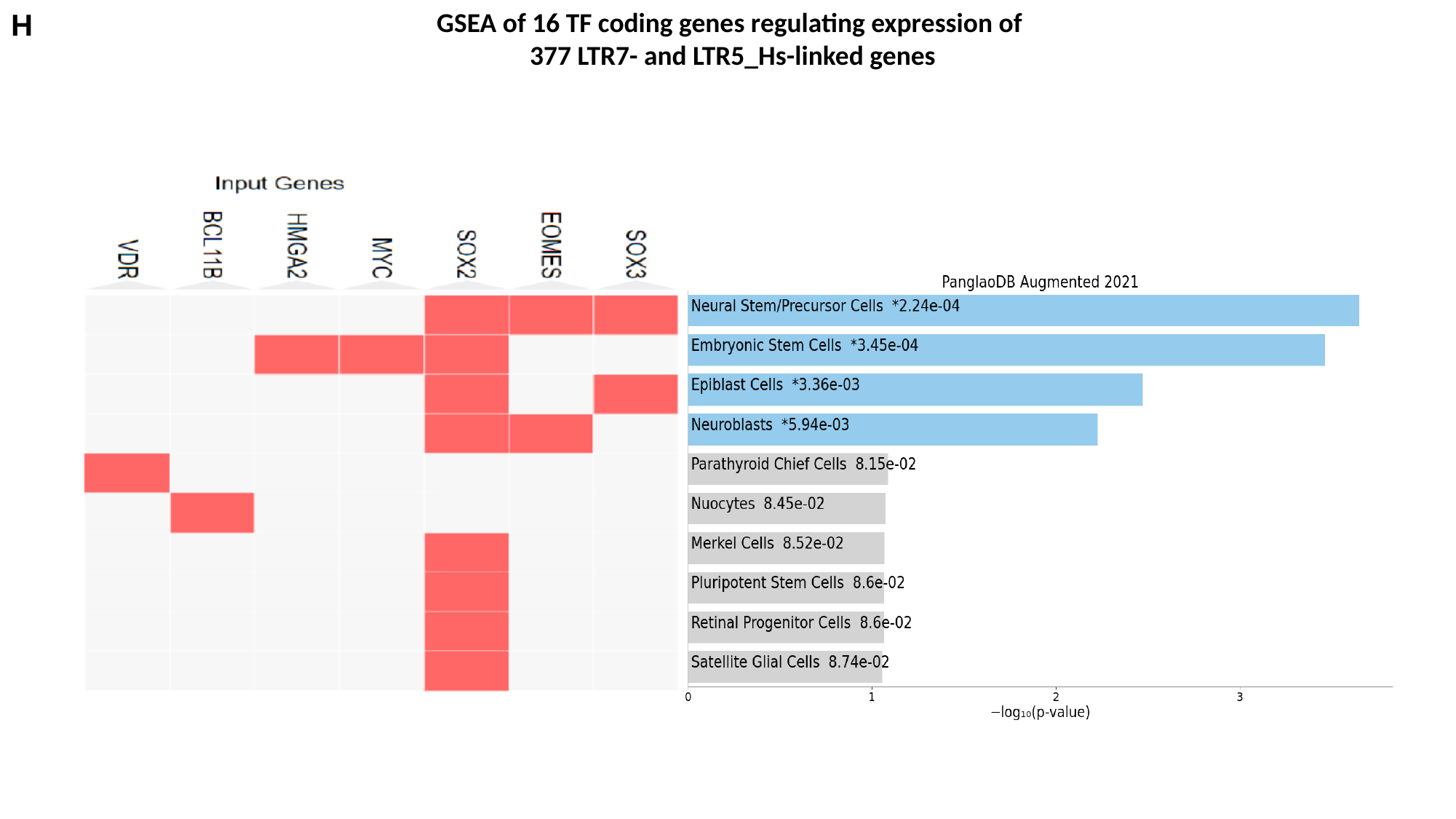

H
GSEA of 16 TF coding genes regulating expression of
377 LTR7- and LTR5_Hs-linked genes

### Slide 10
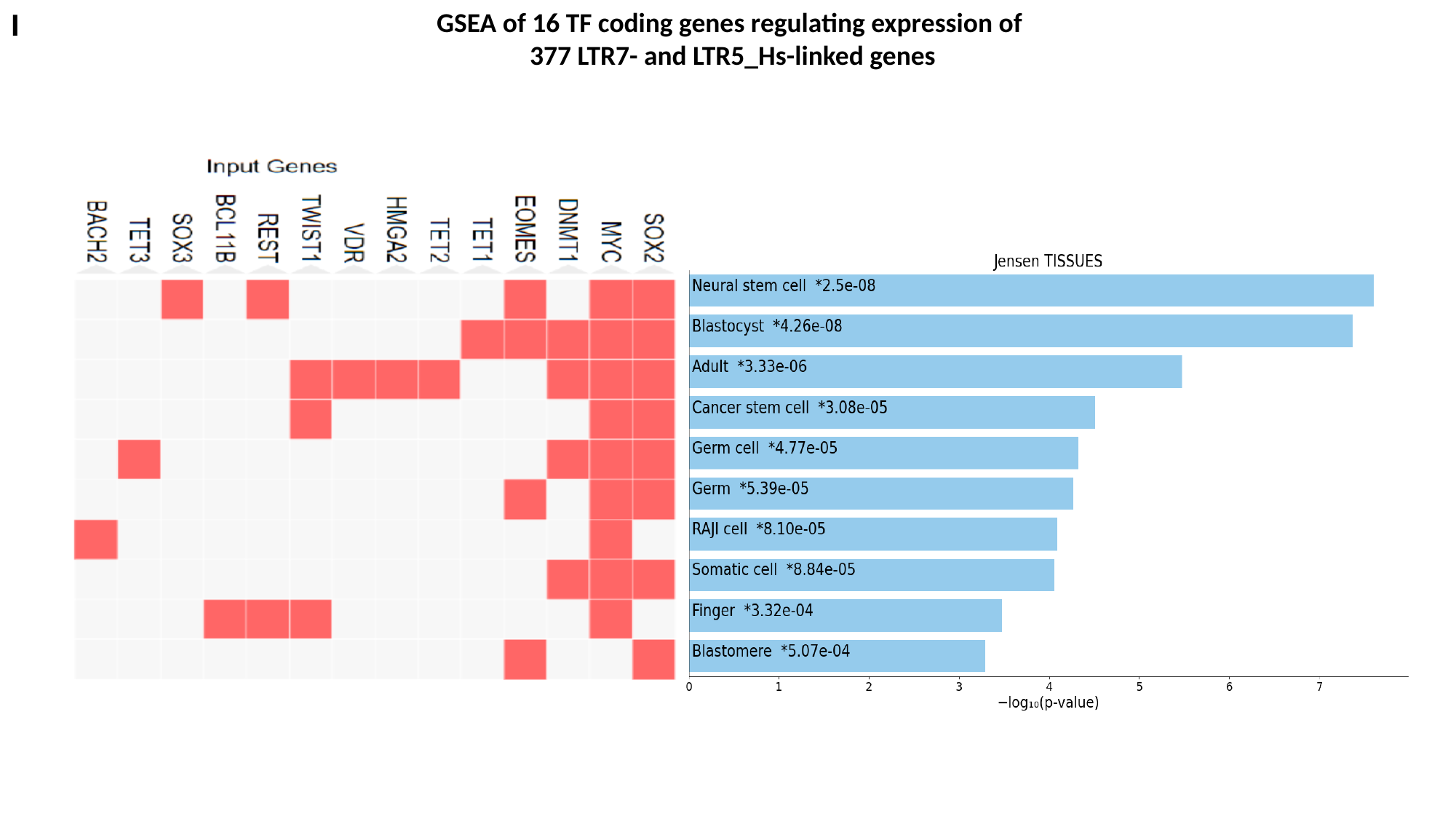

I
GSEA of 16 TF coding genes regulating expression of
377 LTR7- and LTR5_Hs-linked genes

### Slide 11
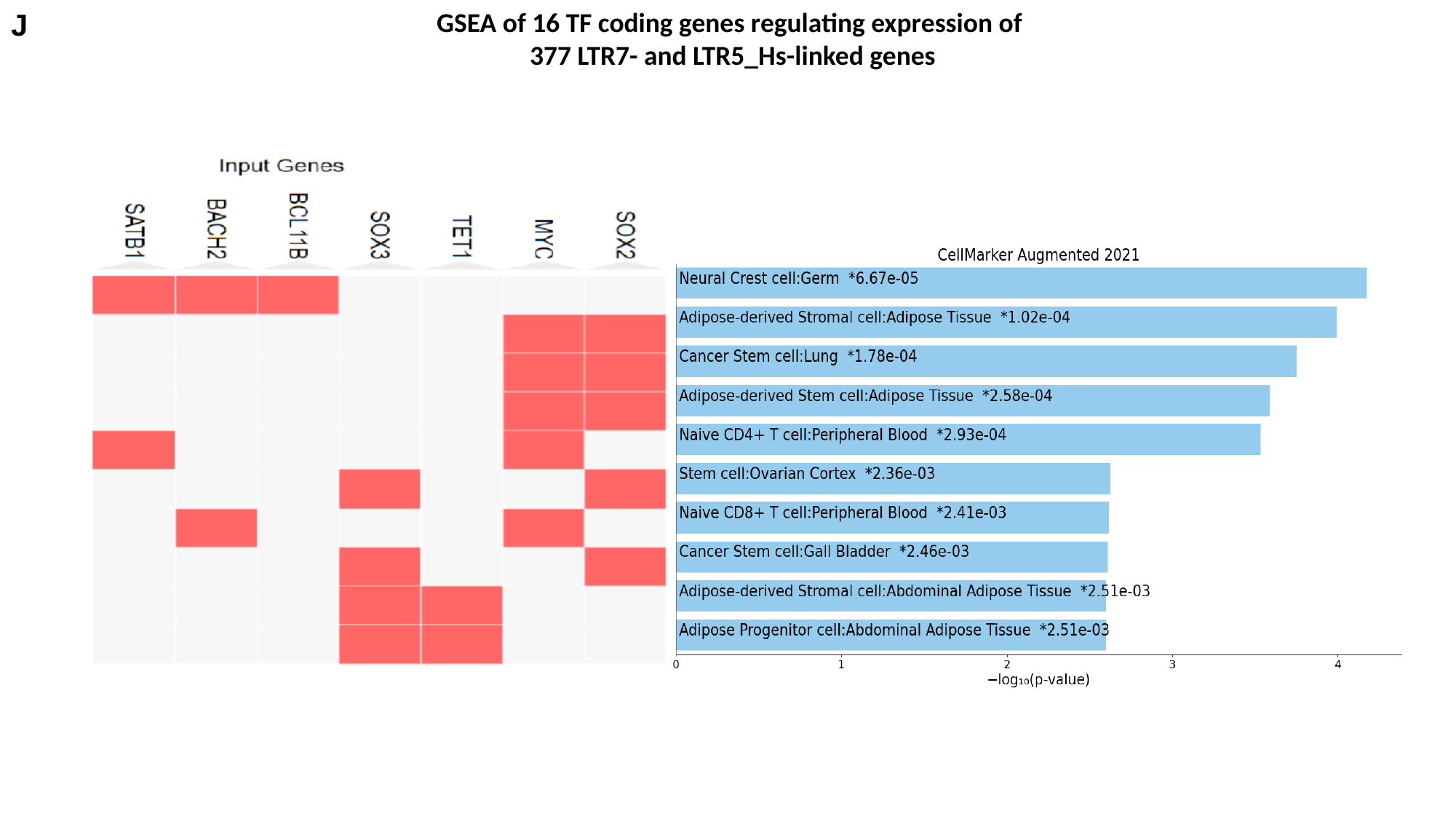

J
GSEA of 16 TF coding genes regulating expression of
377 LTR7- and LTR5_Hs-linked genes

### Slide 12
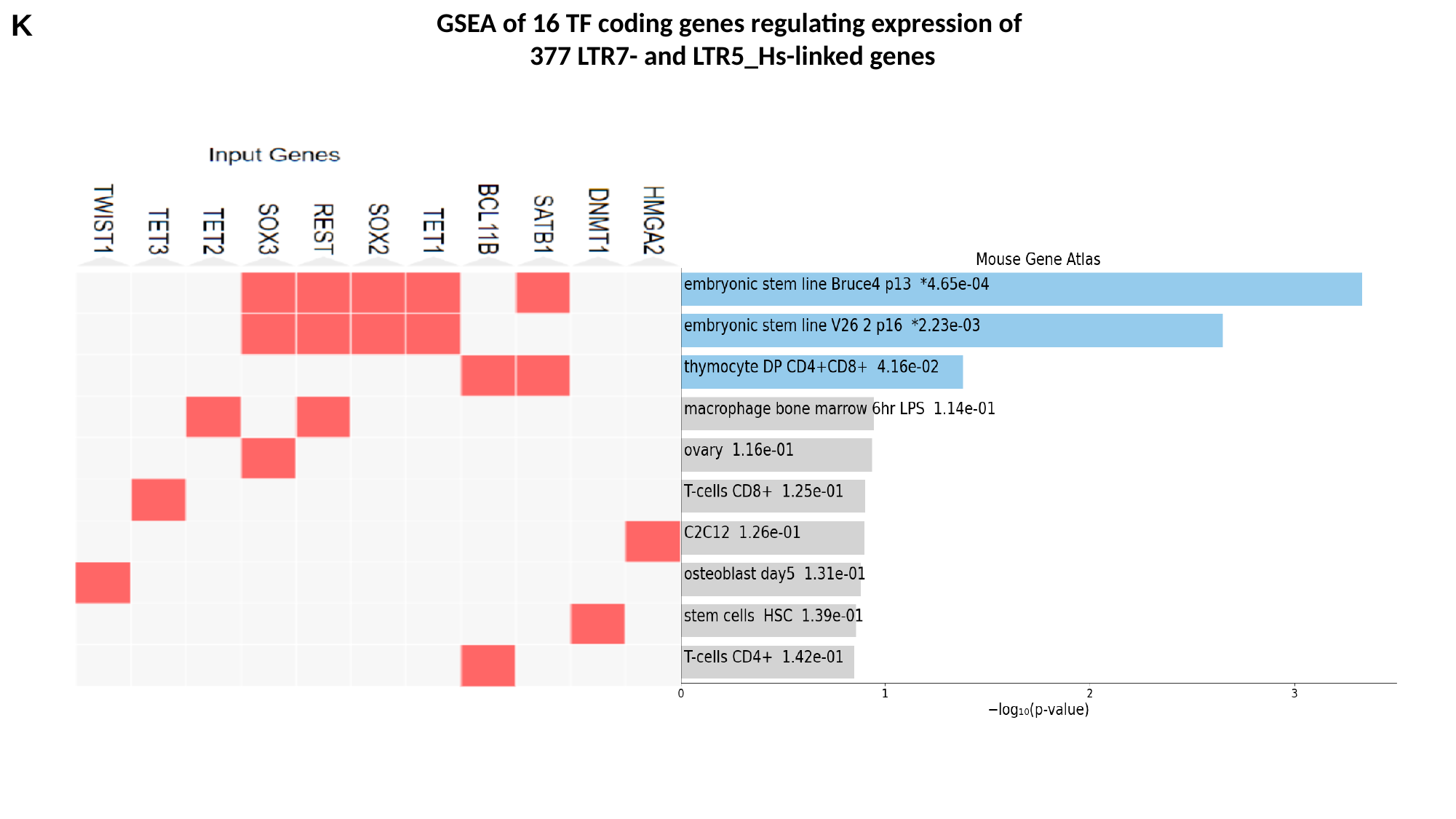

K
GSEA of 16 TF coding genes regulating expression of
377 LTR7- and LTR5_Hs-linked genes

### Slide 13
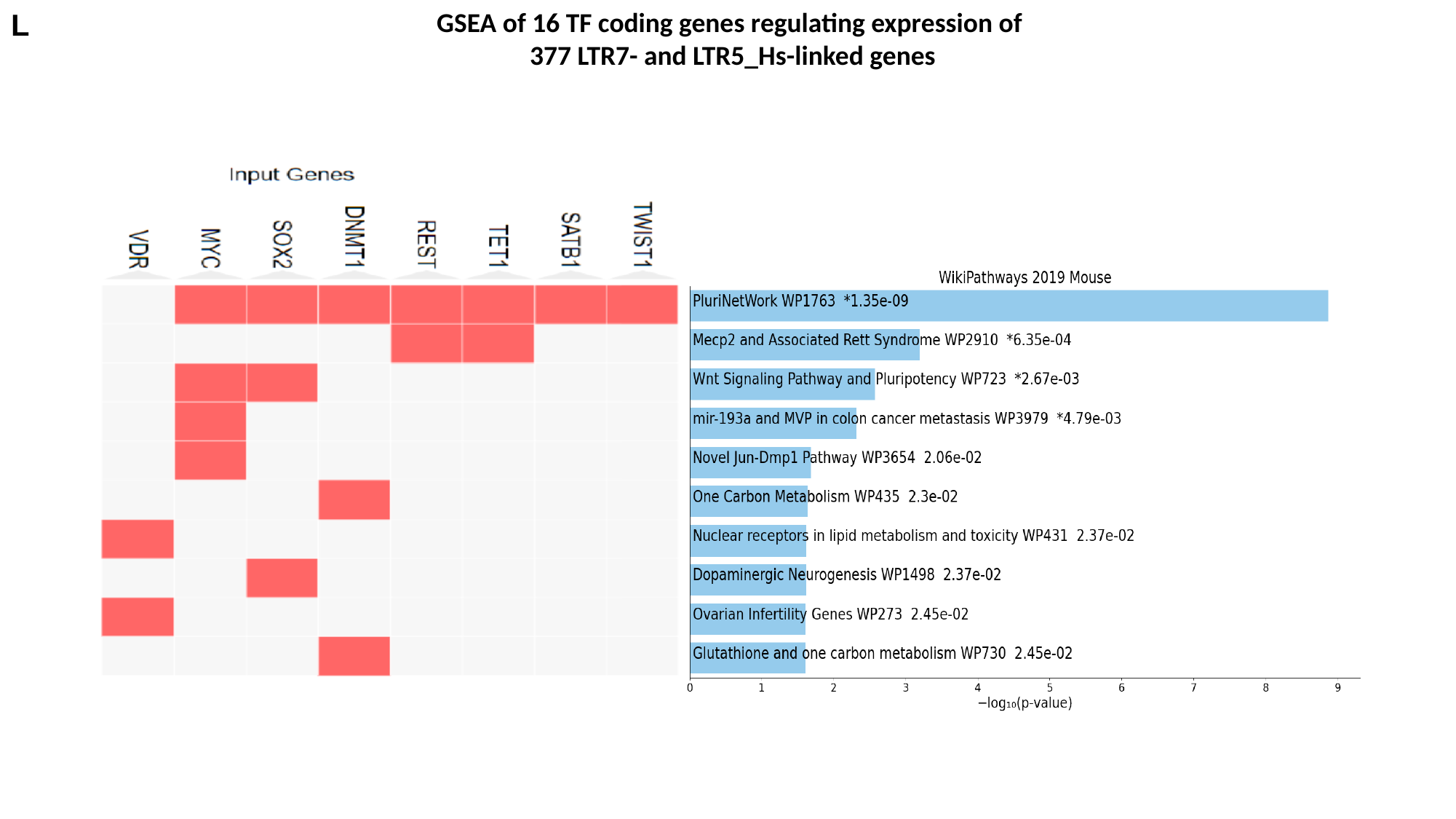

L
GSEA of 16 TF coding genes regulating expression of
377 LTR7- and LTR5_Hs-linked genes

### Slide 14
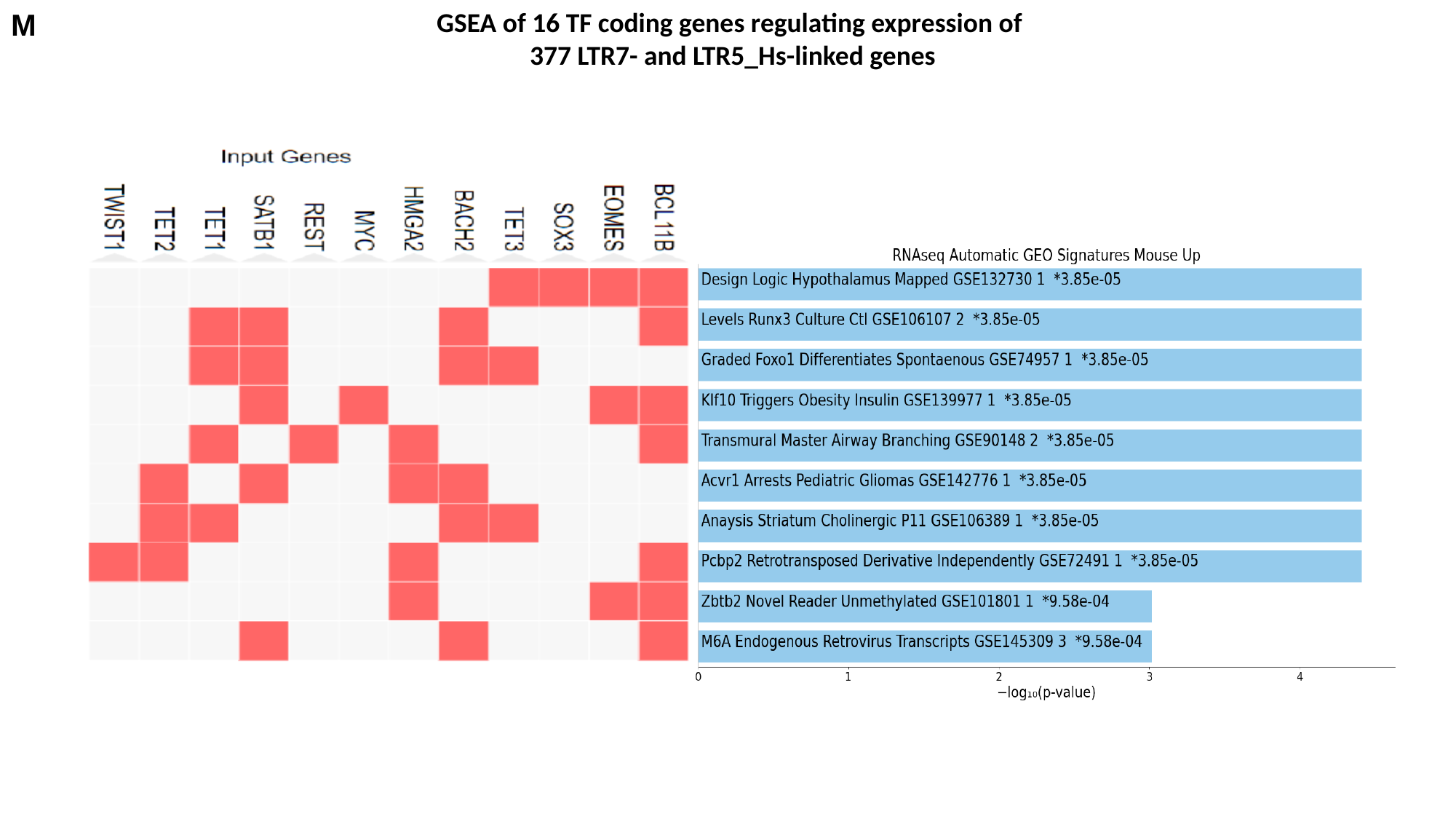

M
GSEA of 16 TF coding genes regulating expression of
377 LTR7- and LTR5_Hs-linked genes
